# Lifelong restriction of dietary valine has sex-specific benefits for health and lifespan in mice

**DOI:** 10.1101/2025.08.31.673254

**Authors:** Mariah F. Calubag, Ismail Ademi, Cara L. Green, Dulmalika N.H. Manchanayake, Hashan S. M. Jayarathne, Ryan N. Marshall, Sandra M. Le, Penelope Lialios, Lucia E. Breuer, Shoshana Yakar, Reji Babygirija, Michelle M. Sonsalla, Isaac Grunow, Chung-Yang Yeh, Yang Liu, Bailey A. Knopf, Sarah Yandell, Charles I. Opara, William A. Ricke, Teresa T. Liu, Mark P. Keller, Alan D. Attie, Mariana Sadagurski, Dudley W. Lamming

## Abstract

Dietary protein is a key regulator of metabolic health in humans and rodents. Many of the benefits of protein restriction are mediated by reduced intake of dietary branched-chain amino acids (BCAAs; leucine, valine and isoleucine), and restriction of the BCAAs is sufficient to extend healthspan and lifespan in mice. While the BCAAs have often been considered as a group, it has become apparent that they have distinct metabolic roles, and we recently found that restriction of isoleucine is sufficient to extend the healthspan and lifespan of male and female mice. Here, we test the effect of lifelong restriction of the BCAA valine on healthy aging. We find that valine restriction (Val-R) improves metabolic health in C57BL/6J mice, promotes leanness and glycemic control across ages, and reduces frailty, cancer prevalence, and senescent cell burden in multiple tissues in both sexes. Val-R reduces glial activation in the brain in a male-specific manner, and extends the lifespan of male, but not female, mice by 23%. To investigate the molecular mechanisms engaged by Val-R with aging, we conducted multi-tissue transcriptional profiling and gene network analysis. While Val-R had a greater molecular impact in the liver, muscle, and brown adipose tissue of females, the enrichment of genes associated with phenotypic traits was stronger in males. Assessing novel gene relationships across tissues, we identified a liver gene module enriched in mitochondrial-related pathways as a central hub. Assessing mitochondrial function, we identified a Val-R-induced male-specific increase in mitochondrial respiration. Our results demonstrate for the first time that Val-R improves multiple aspects of healthspan in mice of both sexes and extends lifespan in males, and suggests that interventions that mimic Val-R may have translational potential for aging and age-related diseases.

## Introduction

Dietary interventions have strong effects on health and lifespan, with calorie restriction extending lifespan in diverse species and improving health in humans ^1, 2^. Recently, it has become clear that interventions that alter dietary intake of specific macronutrients rather than reducing calories can also promote healthy aging. In particular, emerging evidence suggests dietary protein is a key regulator of healthy aging.

Protein consumption has traditionally been thought of as beneficial for healthy aging, as protein promotes satiety and supports weight loss ^3, 4^, and increased protein intake is often recommended for the elderly to combat sarcopenia ^5^. However, several retrospective and prospective cohort studies have shown that higher protein consumption is associated with diseases of aging including diabetes ^6, 7, 8, 9^ and sarcopenia ^10^. Randomized clinical trials have shown that protein restriction (PR) improves metabolic health, reducing adiposity and improving insulin sensitivity ^11, 12, 13^. Finally, multiple studies have shown that PR promotes metabolic health and even extends lifespan in flies and mice ^14, 15, 16, 17, 18^.

Many of the benefits of PR may be due to reduced intake of specific essential amino acids. While restriction of several different amino acids has been shown to improve health without reducing calorie intake ^19, 20, 21^, we have focused on the branched-chain amino acids (BCAAs; leucine, isoleucine and valine). Blood levels of BCAAs are specifically reduced by PR in humans ^12^, and we have found that restriction of the BCAAs improves metabolic health in C57BL/6J mice of both sexes and extends the lifespan of males by over 30% ^17^. Isoleucine is the most potent of the BCAAs in its impact on metabolic health, and restriction of isoleucine alone is sufficient to improve overall metabolic health as well as reduce molecular markers of aging in C57BL/6J mice ^22, 23^. Further, isoleucine restriction extends the lifespan of both flies and UM-HET3 mice ^24, 25, 26^.

While this previous work demonstrates an important effect of isoleucine on metabolism and aging, the impact of restricting either leucine or valine on healthy aging and lifespan has not been investigated. While leucine is a potent agonist of mTORC1, an amino acid-sensitive protein kinase that is a central regulator of metabolism and aging ^27, 28^, restriction of leucine in our hands is associated with negligible metabolic benefits, and leucine supplementation has been shown to not significantly impact lifespan ^22, 29, 30^. In contrast, recent work on valine has shown that valine is associated with cancer, inflammation, insulin resistance and glucotoxicity in mice as well as human and porcine cell culture models ^31, 32, 33, 34, 35^. We have also found that dietary restriction of valine alone can reverse diet-induced obesity and restore glucoregulatory control in C57BL/6J males ^22, 30^.

Here, we investigate the hypothesis that valine restriction (Val-R) increases the healthspan and lifespan of mice. We find that lifelong Val-R improves metabolic health and reduces frailty in C57BL/6J mice of both sexes, and increases the lifespan of male but not female mice when started at one month of age. While Val-R also improves cognitive function in female mice, we observed a stronger reduction in neuroinflammatory glia in Val-R-fed females. We identify sex-specific molecular effects of Val-R on the transcriptome of multiple tissues, including the PI3K-Akt signaling pathway, a pathway whose downregulation is involved in longevity in multiple organisms. Surprisingly though, at the protein level we observed increased Akt activity and activation of its downstream targets in the liver, including mTORC1, a central regulator of aging. An increase in activity of either Akt or mTORC1 would generally be associated with accelerated aging and reduced lifespan, prompting us to look deeper for potential mechanisms to explain the beneficial effects of Val-R on male longevity and healthspan. Analyzing transcriptional networks across tissues, we identified a hub module enriched for mitochondrial metabolism genes; examining mitochondrial activity in the liver, we found that Val-R induces a male-specific increase in mitochondrial respiration. This work suggests that lowering dietary valine should be explored as a potential intervention for age-related diseases and to gain insight into new longevity mechanisms. In conclusion, these results demonstrate the unique role of dietary valine, and expand the universe of dietary components that control healthy aging.

## Methods

### Animal care, housing and diet

All animal procedures were performed in accordance with institutional guidelines and approved by the Institutional Animal Care and Use Committee of the William S. Middleton Memorial Veterans Hospital, under institutional assurance number D16-00403 (Madison, WI, USA). Male and female C57BL/6J mice were purchased from The Jackson Laboratory (Bar Harbor, ME, USA) at 3 weeks of age. All mice were acclimated to the animal research facility for one week before entering studies. All animals were housed in static microisolator cages in a specific pathogen-free mouse facility with a 12:12 h light–dark cycle, maintained at approximately 22°C and housed 2-3 mice per cage.

Mice were fed amino acid-defined diets with either full valine (TD.140711; CTL) or a 67% restriction of valine (TD.160735; Val-R) (full diet compositions are provided in **Table S1**; Inotiv, Madison, WI, USA). Diets were started at 4 weeks of age and continued lifelong.

### Lifespan Study

Mice were randomized into diet groups and enrolled in the survival study at 4 weeks of age (n=25 mice/diet/sex), a group size that provides approximately 90% power to observe a 15% change in lifespan (α = 0.05) ^36^. Mice were euthanized for humane reasons if moribund, developed other problems such as excessive tumor burden or upon the recommendation by the facility veterinarian. Mice found dead were noted during daily inspection and refrigerated. Gross necropsy was performed on euthanized mice and on found dead mice in suitable condition, during which the abdominal and thoracic cavities were examined for the presence of solid tumors, splenomegaly or infection. Based on this inspection, the presence or absence of cancer was noted. A second cohort of mice was sacrificed at 24 months of age for cross-sectional analysis; all mice in this second cohort were also included in the lifespan analysis (n=12 mice/diet/sex). We ended up with n=7-10 mice/group/sex for organ analysis at 24 months of age. Mice were censored as of the date of death if removed for cross-sectional analysis, or if death was due to experimental error (n=1). The lifespan of all mice can be found in **Table S6.**

### Metabolic Phenotyping

Glucose, insulin and alanine tolerance tests were performed by fasting all mice for 4 hours or overnight (∼16 hours) and then injecting either glucose (1g/kg), insulin (0.75U/kg) or alanine (2g/kg) intraperitoneally ^37, 38^.

Blood glucose levels were determined at the indicated times using a Bayer Contour blood glucose meter (Bayer, Leverkusen, Germany) and test strips. Body composition was determined using an EchoMRI Body Composition Analyzer. For assay of multiple metabolic parameters (O_2_, CO_2_, food consumption, and activity tracking), mice were acclimatized to housing in a Columbus Instruments Oxymax/CLAMS-HC metabolic chamber system for approximately 24 hours, and data from a continuous 24-hour period was then recorded and analyzed.

### Physical fitness testing via rotarod and inverted cling assays

To assess motor coordination via a rotarod assay, mice were trained at a constant speed of 4 rpm the day before testing. On the day of testing, mice were placed on the rotarod for three rounds, at least 30 min apart, and the average time spent on the rotarod, and the maximum speed were recorded. During the testing, the rotarod started at a speed of 4 rpm with an acceleration of 0.5 rpm/s up to a maximum of 40 rpm. To assess grip strength via the inverted cling test, mice were placed on a wire frame and carefully inverted until the mice were hanging upside down. The timer then started, and the time until the mouse fell was recorded. The average time of three rounds of testing conducted at least 30 min apart was calculated.

### Frailty Index Scoring

Frailty was assessed longitudinally using a 30-item list frailty index whose measures are based on procedures outlined in ^39^; only mice enrolled in the full lifespan study, and not those enrolled for cross-sectional sacrifice, were scored. This frailty index reflects an accumulation of age-associated deficits similar to the Rockwood frailty index in humans ^40^. The items are scored from 0 (no deficit) to 0.5 (mild deficit) to 1 (severe deficit). The 29 criteria scored throughout life include alopecia, body weight, loss of fur color, dermatitis, loss of whiskers, coat condition, tumors, distended abdomen, kyphosis, tail stiffening, gait disorders, tremor, body condition score, vestibular disturbance, cataracts, corneal opacity, eye discharge/swelling, microphthalmia, vision loss, menace reflex, nasal discharge, malocclusions, rectal prolapse, vaginal/uterine/penile prolapse, diarrhea, breathing rate/depth, mouse grimace score, and piloerection. Grip strength (assessed by inverted cling time) was scored as a 30^th^ criteria up to 24 months of age. The scores for all items are averaged to give the frailty score. These tests were conducted at 12-13, 18, 24, 28, 31-32 and 32-33 months of age. Complete frailty scores can be found in **Tables S7 and S8**.

### Void Spot Assay

Void spot assays were performed as described previously ^26, 41^. Mice were individually placed in standard mouse cages with thick chromatography paper (Ahlstrom, Kaukauna, WI). During the study period (4 hours), mice were restricted from water. Chromatography papers were imaged with a BioRad ChemiDoc Imaging System (BioRad, Hercules, CA) using an ethidium bromide filter set and 0.5 second exposure to ultraviolet light. Images were imported into ImageJ and total void spots analyzed with VoidWhizzard.

### Novel Object Recognition (NOR) assay

A novel object recognition test (NOR) was performed in an open field where the movements of the mouse were recorded using a camera mounted above the field as previously described ^42^. Before each test, mice were acclimatized in the behavioral room for at least 30 min and were given a 5-minute habituation trial with no objects on the field. This was followed by a short-term memory test (STM) phase on the same day, consisting of one acquisition trial and one test trial. The next day, we conducted a long-term memory test (LTM) consisting of only a test trial. In the acquisition trial, the mice were allowed to explore two identical objects placed diagonally on opposite sides of the field for 5 minutes. One hour after the acquisition trial, STM was performed, and 24 hours later, LTM was performed. Both test trials were performed by replacing one of the identical objects of the acquisition trial with a novel object. The results were quantified using a discrimination index (DI), which represents the ratio of the duration of exploration for the novel object to the duration of exploration of the old object.

### Collection of tissues for molecular and histological analysis

Mice were euthanized in the fed state at 24 months of age, where they were fasted overnight starting the day prior to sacrifice; in the morning, mice were refed for 3 hours and then sacrificed. Following blood collection via submandibular bleeding, mice were euthanized by cervical dislocation and tissues were rapidly collected, weighed, and snap frozen in liquid nitrogen. A portion of the liver was directly embedded into Tissue-Tek Optimal Cutting Temperature (OCT) compound. It was sent to the University of Wisconsin-Madison Carbone Cancer Center Experimental Animal Pathology Laboratory (UWCCC EAPL) for cryosectioning and staining for Oil-Red-O (ORO). Another portion of the liver as well as a portion of the kidney and spleen was fixed in 10% formalin for 4 hours, transferred to 30% sucrose for 24 hours and then embedded in OCT and then cryosectioned and stained for SA-β-Gal activity. The inguinal white adipose tissue (iWAT) and brown adipose tissue (BAT) were fixed in 10% formalin for 24 hours, switched to 70% ethanol and then paraffin-embedded before being cryosectioned and stained for Hematoxylin and eosin (HE). Images of the liver, kidney, iWAT and BAT were taken using an EVOS microscope (Thermo Fisher Scientific Inc., Waltham, MA, USA) at a magnification of 40X as previously described ^43, 44^. Quantification fields were obtained for each tissue from each mouse and quantified using ImageJ (NIH, Bethesda, MD, USA).

For histological analysis, brains were fixed in 10% formalin for 24 hours and transferred to 30% sucrose. Brains were postfixed, dehydrated, and then sectioned coronally (30 μm) using a sliding microtome, followed by immunofluorescent analysis as described ^45^. For immunohistochemistry, brain sections were washed with PBS six times, blocked with 0.3% Triton X-100 and 3% normal donkey serum in PBS for 2 hours before staining was carried out overnight using rabbit anti-GFAP (1:1000; Millipore, Cat. No. ab5804 primary antibody. For goat anti-Iba1 (1:1000 Abcam Cat. No. ab5076), immunostaining brain sections were pretreated with 0.5% NaOH and 0.5% H_2_O_2_ in PBS for 20 minutes. After the primary antibody brain sections were incubated with AlexaFluor-conjugated secondary antibodies for 2 hours (Invitrogen). Microscopic images of the stained sections were obtained using an Olympus FluoView 500 and Zeiss LSM 800 Laser Scanning Confocal Microscope.

### Astrocyte and Microglia Morphology Analysis

Immunofluorescence images were taken using a multiphoton laser-scanning microscope (LSM 800, ZEISS) equipped with a 63X objective for the hippocampus. Stacked images along the optical axis (z-axis), were reconstructed to 3D using Fiji-Image J. These were analyzed for cellular morphology, skeleton and fractal analyses using an established protocol ^46, 47^. Skeletal and fractal analysis parameters were exported into separate Excel files and used for data analysis. All images used for analysis were taken with the same confocal settings (pinhole, digital gain, and digital offset). Image processing, three-dimensional reconstruction, and data analysis were performed in a blinded manner with respect to the experimental conditions.

### Micro-computed tomography

Right femurs from all animals were collected for micro-computed tomography (μCT). Bones were fixed in 10% formalin and placed on a rocker for 24 hours before being rinsed, placed in 70% ethanol, and stored at 4°C. All bones were scanned using the same instrument under the same conditions, following the American Society for Bone and Mineral Research guidelines ^48^. Using a high-resolution SkyScan micro-CT system (SkyScan 1172, Kontich, Belgium) with 10-MP digital detector, 10 W of energy (60 kV and 167 mA), and a pixel size of 9.7 microns, exposure 925 ms/frame rotation step 0.3 degrees with ×10 frame averaging, 0.5-mm aluminum filter (to increase the transmission), samples were scanned in the air with scan rotation of 180 degrees. Before morphometric analysis, global thresholding was applied. Image reconstruction was done using NRecon software (version 1.7.3.0; Bruker micro-CT, Kontich, Belgium). Data analysis was done using CTAn software (version 1.17.7.2+; Bruker micro-CT, Kontich, Belgium). 3D images were constructed using CT Vox software (version 3.3.0 r1403; Bruker micro-CT, Kontich, Belgium).

### Immunoblotting

Tissue samples from liver and muscle were lysed in cold RIPA buffer supplemented with phosphatase inhibitor and protease inhibitor cocktail tablets (Thermo Fisher Scientific, Waltham, MA, USA) as previously described ^17, 49^ using a FastPrep 24 (M.P. Biomedicals, Santa Ana, CA, USA) with screw cap microcentrifuge tubes (822-S) from (Dot Scientific, Burton, MI) and ceramic oxide bulk beads (10158-552) from VWR (Radnor, PA, USA). Protein lysates were then centrifuged at 13,300 rpm for 10 minutes and the supernatant was collected. Protein concentration was determined by Bradford (Pierce Biotechnology, Waltham, MA, USA). 10-20 μg protein was separated by SDS–PAGE (sodium dodecyl sulfate–polyacrylamide gel electrophoresis) on 10 and 16% resolving gels (Thermo Fisher Scientific, Waltham, MA, USA) and transferred to PVDF membrane (EMD Millipore, Burlington, MA, USA). pT389-S6K1 (9234), S6K1 (9202), pS240/244-S6 (2215), S6 (2217), pThr37/46 4E-BP1 (2855), 4E-BP1 (9644), eIF2α (5324), pS51-eIF2α (3597), AMPKa (5831), pAMPKa (4188), Beclin-1 (3495), LC3A/B (12741), AKT (pan) (C67E7), pAKT S473(4060L), p44/42 MAPK (Erk1/2) (9102), p-p44/42 MAPK (Erk1/2) (Thr202/Tyr204) (9101), p-FoxO1 (Thr24)/FoxO3a (Thr32) (9464), VDAC (4866), GAPDH (2118), and β-tubulin (2146) were purchased from Cell Signaling Technologies (CST, Danvers, MA, USA) and used at a dilution of 1:1000. p62 (American Research Products, #03-GP62-C) was also used at a dilution of 1:1000. Total OXPHOS Rodent Antibody Cocktail (Abcam, ab110413) was also used at a dilution of 1:1000. Imaging was performed using a Bio-Rad Chemidoc MP imaging station (Bio-Rad, Hercules, CA, USA). Quantification was performed by densitometry using NIH ImageJ software.

### Quantitative real-time PCR (qRT-PCR)

qRT-PCR was carried out according to protocols as previously described ^43^. using TRI Reagent according to the manufacturer’s protocol. The concentration and purity of RNA were determined by absorbance at 260/280 nm using Nanodrop (Thermo Fisher Scientific). 1 μg of RNA was used to generate cDNA (Superscript III; Invitrogen, Carlsbad, CA, USA). Oligo dT primers and primers for real-time PCR were obtained from Integrated DNA Technologies (IDT, Coralville, IA, USA). Reactions were run on an StepOne Plus machine (Applied Biosystems, Foster City, CA, USA) with Sybr Green PCR Master Mix (Invitrogen). Actin was used to normalize gene-specific reaction results.

The primers that were used are as follows:

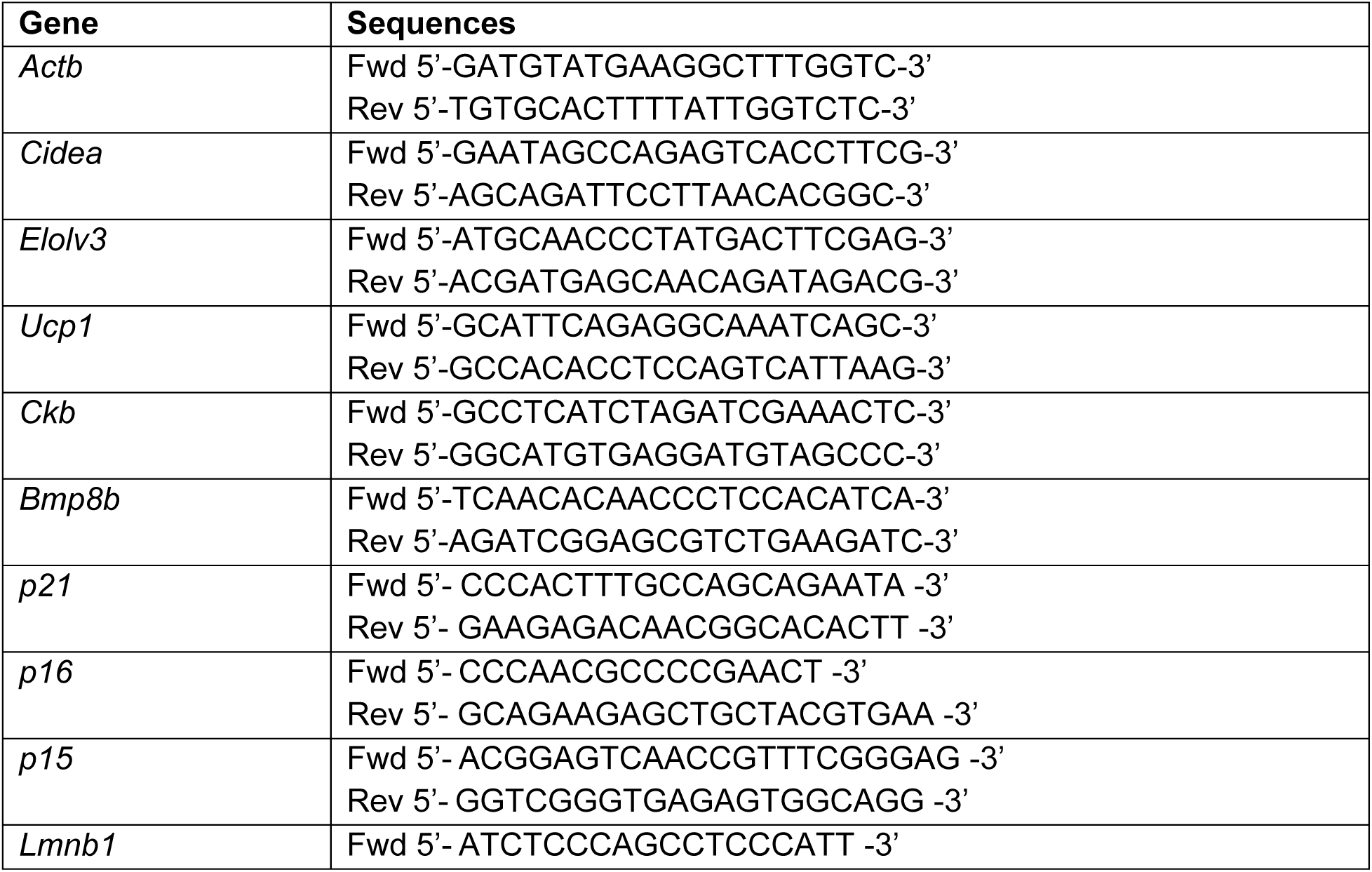
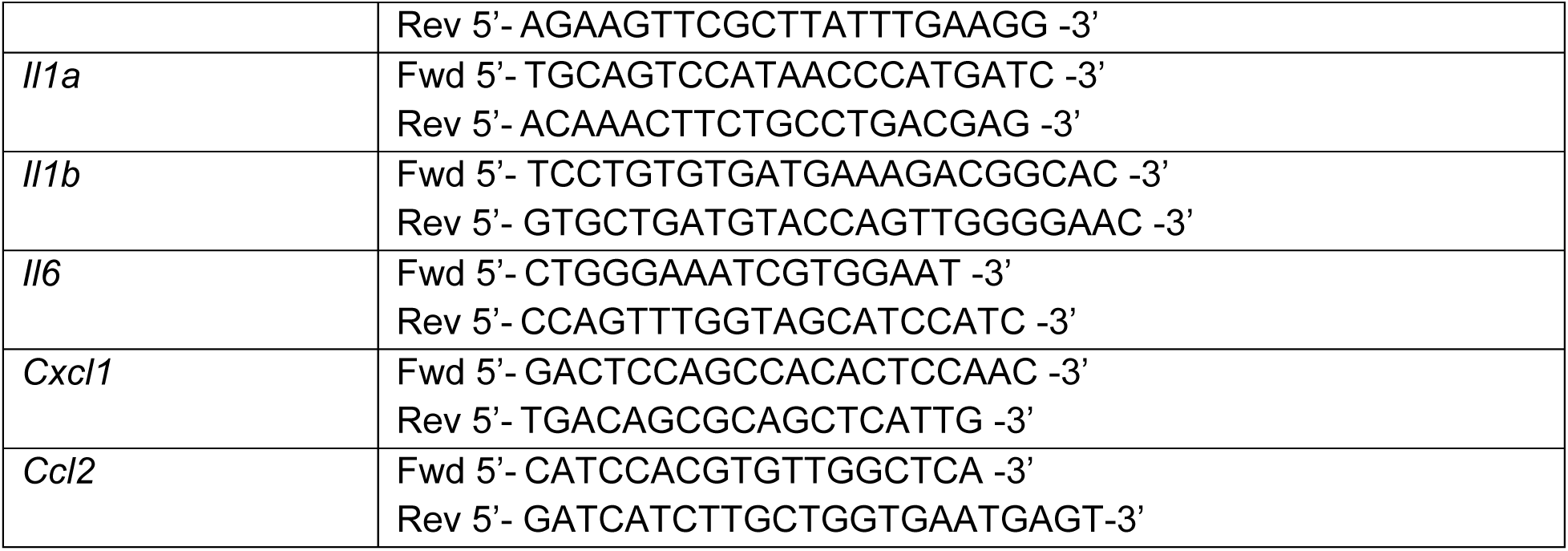

### Senescence-Associated B-Galactosidase Staining

Enhanced lysosomal biogenesis, a common feature of cellular senescence, can be detected by measuring β-galactosidase (lysosomal hydrolase) activity at pH 6.0 ^50^. This senescence-associated β-galactosidase (SA-β-Gal) activity appears to be restricted to senescent cells at a low pH ^51, 52^. Fresh tissue from mice was collected and fixed in 10% neutral buffered formalin (NBF) on ice for 3-4 hr. Tissues were then transferred to 30% sucrose at 4°C for 24 hours before being embedded in O.C.T. compound in a cryomold and stored at −80°C. Before cryosectioning, tissues were equilibrated at −20°C and then cryosectioned into 5-7 µm sections before being attached to Superfrost Plus slides. Fresh SA-β-Gal staining solution at a pH of 6 was prepared as previously described. Tissue slides were stained with SA-β-Gal staining solution for 18-24 hrs at 37°C in a non-CO2 incubator and then rinsed three times with PBS. To prevent crystal formation a parafilm Coplin jar was used to prevent evaporative loss of staining solution. Stained sections were imaged using the EVOS microscope (Thermo Fisher Scientific Inc., Waltham, MA, USA) at a magnification of 40X as previously described. The percent of SA-βgal-positive area for each sample was quantified using ImageJ.

### ELISA assays and kits

Blood plasma for FGF21 and insulin was obtained at 19 months of age in the fasted state. Blood FGF21 levels were assayed by a mouse/rat FGF-21 quantikine ELISA kit (MF2100) from R&D Systems (Minneapolis, MN, USA). Plasma insulin was quantified using an ultra-sensitive mouse insulin ELISA kit (90080), from Crystal Chem (Elk Grove Village, IL, USA). Triglycerides were measured by Triglyceride Colorimetric Assay Kit (Cayman Chemical Company, Ann Arbor, MI, USA; Item No. 10010303) using plasma and tissue collected at sacrifice.

### Transcriptomics

RNA was extracted from the liver, BAT and muscle using the PureLink RNA mini kit (Invitrogen, 12183025) with DNase (Invitrogen, 12185010) following manufacturer’s instructions. The concentration and purity of RNA was determined using a NanoDrop 2000c spectrophotometer (Thermo Fisher Scientific, Waltham, MA) and RNA was diluted to 100–400 ng/mL for sequencing. Total RNA was submitted to the University of Wisconsin-Madison Biotechnology Center Gene Expression Center (RRID:SCR_017757) & DNA Sequencing Facility (RRID:SCR_017759) for RNA quality assessment on an Agilent Biomek Plate Reader (A260/A280) and Agilent 4200Tapestation (RIN). RNA libraries were prepared using the NEBNext® Ultra™ II Directional RNA Library Prep Kit (Illumina, New England Biolabs GmbH, Frankfurt, DE) with a 500ng total RNA input. Paired-end 150bp sequencing was done on an Illumina NovaSeq X Plus sequencer. Adapter-trimmed strand-specific 2×150 bp Illumina reads were processed with Skewer v0.1.123 (Jiang et al., 2014) to remove sequencing adapters and low-quality bases ^53^. Reads were aligned to the *Mus musculus* GRCm39 reference genome (NCBI assembly accession GCA_000001635.9) using STAR v2.7.11b (Dobin et al., 2013) with splice-aware alignment and transcript annotations from Ensembl release 110 ^54^. STAR was run with the options --twopassMode Basic to improve splice junction discovery and --outSAMtype BAM SortedByCoordinate to produce coordinate-sorted BAM files suitable for downstream analysis. Expression quantification at the gene and transcript levels was performed with RSEM v1.3.1 (Li and Dewey, 2011) using the STAR-aligned BAM files as input. The RSEM reference was prepared using the corresponding Ensembl transcript annotations ^55^.

Analysis of significantly differentially expressed genes (DEGs) was completed in R version 4.4.3 using *edgeR* and *limma* packages. Gene names were converted to gene symbol and Entrez ID formats using the *mygene* package. PCA plots were generated using the *mixomics* package. DEGs were used to identify enriched pathways, both Gene Ontology (for Biological Processes) and KEGG enriched pathways using an adjusted p-value cut-off of 0.05 (**Tables S2-S5**). All genes, log2 fold-changes and corresponding unadjusted and Benjamini-Hochberg adjusted p values can be found in **Tables S2-S5**.

WGCNA analysis was conducted in R (Version 4.4.3) using the WGCNA package. We analyzed males and females separately. First, we filtered by gene expression variance, keeping the top 50% variable genes for each tissue to remove “noisy” genes. We combined genes from all three tissues, which were demarcated to indicate their tissue origin. We then checked that all genes had enough samples and there were no clear outliers before running the analysis. We started with 25197 genes for females and 22928 for males. WGCNA analysis identifies significant gene modules and their correlations with phenotypes. Once gene modules were identified, they were enriched for KEGG Pathways. We then separated out genes by their original tissue type and re-ran the KEGG enrichment analysis to identify tissue specific pathways that were related to phenotypes of interest.

### Cytoscape network

For the inter-tissue network analysis, we first matched the mice used for transcriptomics in BAT, Liver and Muscle, yielding 28 total mice (6 and 6 for male mice on control and ValR diets, and 7 and 9 female mice on control and ValR diets, respectively). Co-expression gene modules were computed in R using the WGCNA package ^56^ as we have previously described following rankz-transformation of gene expression data ^57^. Among all sequenced transcripts, an average of ∼94% were included in 18, 24 and 22 modules for BAT, Liver and Muscle, respectively. Gene expression and WGCNA module gene membership is available at https://connect.doit.wisc.edu/dlamming_valine_study/ME-ME correlations were used to construct a correlation network in Cytoscape (v3.10.4) where the nodes are co-expression gene modules denoted by their ME.colorname. The color of each module reflects the tissue of origin, as indicated in the legend. Edges connecting nodes represent the correlation between the Module Eigengene (ME) for the modules identified in BAT, Liver and Muscle. Edge color indicates direction of correlation (red = positive, blue = negative), thickness is controlled by the strength of correlation (|correlation|, range, 0.37 to 1.0) and transparency is set by the -log_10_ P-value (range, 1 to 9). The relative position of the nodes and curved edges were determined by the yFiles Radial Layout option in Cytoscape.

### Mitochondrial Respiratory Analysis

Liver mitochondrial oxygen consumption rates (OCR) for respiratory complexes I, II, and IV were measured in frozen liver samples using a Seahorse XFe96 Analyzer, as previously described ^58, 59^. Briefly, 50 mg of liver tissue was thawed in 3 mL of ice-cold 1× mitochondrial assay solution (MAS; 70 mM sucrose, 220 mM mannitol, 5 mM KH₂PO₄, 5 mM MgCl₂, 1 mM EGTA, 2 mM HEPES, pH 7.4), finely minced with scissors, and homogenized on ice using 15 strokes of a tight-fitting Dounce homogenizer. Homogenates were centrifuged at 1,000 × g for 10 min at 4°C to remove debris, and the supernatant was subsequently centrifuged at 10,000 × g to isolate crude mitochondria. The mitochondrial pellet was washed twice in 1× MAS buffer and resuspended in 100 μL ice-cold MAS buffer with vortexing. Protein concentration was determined, and 3 μg of mitochondrial protein was loaded per well in triplicate.

Complex I driven respiration was assessed using NADH (1 mM), and complex II driven respiration was measured using succinate (5 mM) in the presence of rotenone (2 μM). Rotenone (2 μM) and antimycin A (4 μM) were subsequently injected to inhibit complexes I and III, respectively. Complex IV activity was stimulated with ascorbate (1 mM), and non-mitochondrial respiration was determined following the addition of sodium azide (40 mM).

To assess intrinsic mitochondrial capacity, OCR values were normalized to citrate synthase (CS) activity ^60^. CS activity was measured as previously described ^61^ and adapted to a 96-well format ^62^. In duplicate, 10 μL of mitochondrial protein was added to 186 μL of assay buffer (50 mM potassium phosphate, pH 7.4, 100 μM DTNB, 115 μM acetyl-CoA) and baseline absorbance was recorded at 15 s intervals for 3 min. The reaction was initiated by addition of 4 μL of 5 mM oxaloacetate (final concentration 100 μM), and absorbance was monitored for an additional 3 min. Enzyme activity (nmol·min⁻¹·mg⁻¹) was calculated from absorbance values, corrected for pathlength, with a within-plate coefficient of variation of 3.6% across technical replicates.

### Blinding

Investigators were blinded to diet groups during data collection whenever feasible, but this was not always possible or feasible as cages were clearly marked to indicate the diet provided, diets were color-coded to prevent feeding mistakes, and the size and body composition of the mice was altered by the diet. Researchers were blinded to group allocations during image analysis. During analysis of other datasets, the investigators are not blinded as the results are objective quantifications. Investigators were not blinded during necropsies.

### Statistics

Data are presented as the mean ± SEM unless otherwise specified. Statistical analyses were performed using one-way or two-way ANOVA followed by Tukey–Kramer post hoc test, as specified in the figure legends. Outliers were excluded using the Robust Regression Outlier Test (ROUT) in Graphpad Prism (v10), Q=1%, and are indicated by an asterisk(*) and blue colored font in the Source Data. Lifespan comparisons were calculated by log-rank test. Maximum lifespan calculations were made by generating a cutoff of the top 25% longest lived animals in each sex, coupled with Boschloo’s Test (Wang-Allison) for significance testing between groups. Healthspan by FAMY (Frailty Adjusted Mouse Years) and GRAIL (Gauging Robust Aging when Increasing Lifespan) were calculated as described ^63^. Other statistical details are available in the figure legends. Energy expenditure differences were detected using analysis of covariance (ANCOVA). ANCOVA analysis assumes a linear relationship between the variables and their covariates. If the slope is equal between groups, then the regression lines are parallel, and elevation is then tested to determine any differences (i.e., if slopes are statistically significantly different, elevation will not be determined). In all figures, n represents the number of biologically independent animals. Sample sizes were chosen based on our previously published experimental results with the effects of dietary interventions. Data distribution was assumed to be normal, but this was not formally tested.

### Randomization

All studies were performed on animals or on tissues collected from animals. Young animals of each sex were randomized into groups of equivalent weight, housed 3 animals per cage, before the beginning of the *in vivo* studies.

## Results

### Valine restriction improves the metabolic health of male and female mice

We began our lifespan study by randomizing male and female C57BL/6J mice to one of two amino acid (AA)-defined diets starting at 4 weeks of age and followed them longitudinally (**Fig. 1A**). Briefly, our control (CTL) diet contained all twenty common AAs; the diet composition reflects that of a natural chow in which 21% of calories are derived from protein. We also utilized a diet in which the level of valine was reduced by 67% (valine-restricted, Val-R). These diets are isocaloric, with identical levels of fat and carbohydrates; the reduction in valine was balanced by a proportional increase in non-essential AAs, keeping the percentage of calories derived from AAs constant. The full composition of these diets is summarized in **Table S1**.

**Figure 1:**
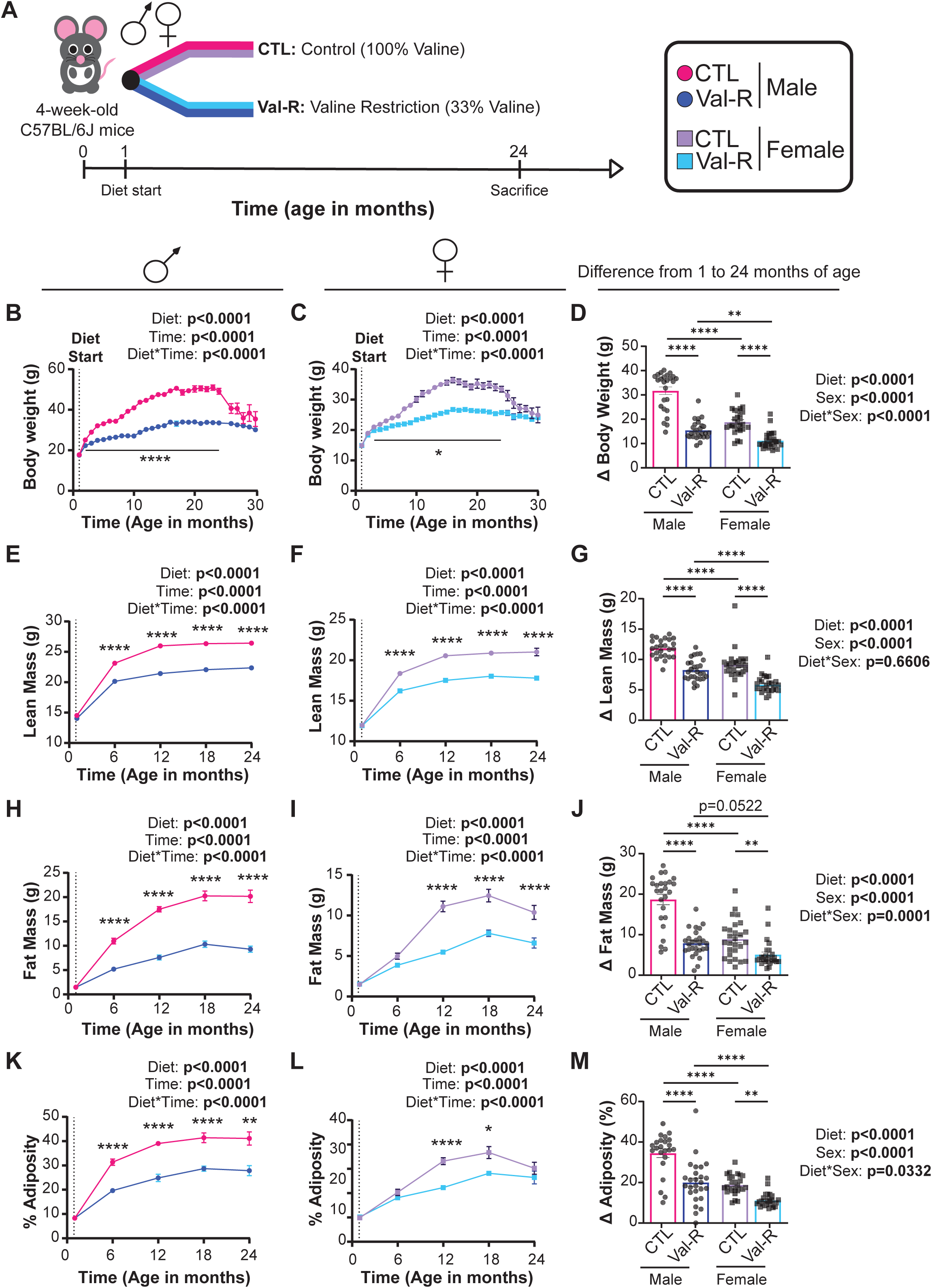
Val-R attenuates body weight and fat mass accretion in both males and females. (A) Experimental design. (B-D) Body weight of male (B) and female (C) mice over 30 months and change in body weight from 1 month of age until 24 months of age (D). (E-G) Lean mass of male (E) and female (F) mice over 30 months and change in lean mass from 1 month of age until 24 months of age (G). (H-J) Fat mass of male (H) and female (I) mice over 30 months and change in body weight from 1 month of age until 24 months of age (J). (K-M) Percent adiposity of male (K) and female (L) mice over 30 months and change in body weight from 1 month of age until 24 months of age (M). (B-M) n=37 mice/group; two-way ANOVA, *p<0.05, **p<0.01, ***p<0.001, ****p<0.0001; statistics for the overall effects of time or sex, diet, and the interaction represent the p value from a two-way ANOVA. Data represented as mean ± SEM.

We measured body weight monthly and assessed body composition every 6 months. Val-R-fed mice of both sexes gained weight more slowly than their CTL-fed counterparts, with a highly significant difference at 24 months of age (**Figs. 1B-D**). The lower weight of Val-R-fed mice was the result of reduced accretion of both lean mass and fat mass; the greater impact on fat mass resulted in an overall reduction in adiposity in both sexes (**Figs. 1E-M**). In addition to being leaner and lighter, Val-R fed mice had reduced iWAT and eWAT mass, as well as a reduced iWAT to eWAT ratio, but increased mass of the quadricep muscle relative to body weight (**Supplementary Fig. 1**).

We also assessed the effects of Val-R on bone morphology using micro-CT. We found no differences in femur and tibia length in Val-R-fed mice of both sexes, demonstrating no effect of diet on skeletal growth (**Supplementary Figs. 2A-C**). In the femur, Val-R significantly decreased total cross-sectional area (T.Ar) with a proportional decrease in marrow area (M.Ar), in male but not female mice, indicating inhibition of radial bone expansion (**Supplementary Figs. 2D-M**). Reduction in T.Ar in male mice was associated with reduced mean polar moment of inertia (**Supplementary Figs. 2E, 2G**). In the trabecular bone compartment at the femur distal metaphysis, Val-R-fed female mice showed reduced trabecular thickness, suggesting reduced bone remodeling (**Supplementary Figs. 2J-M**).

**Figure 2:**
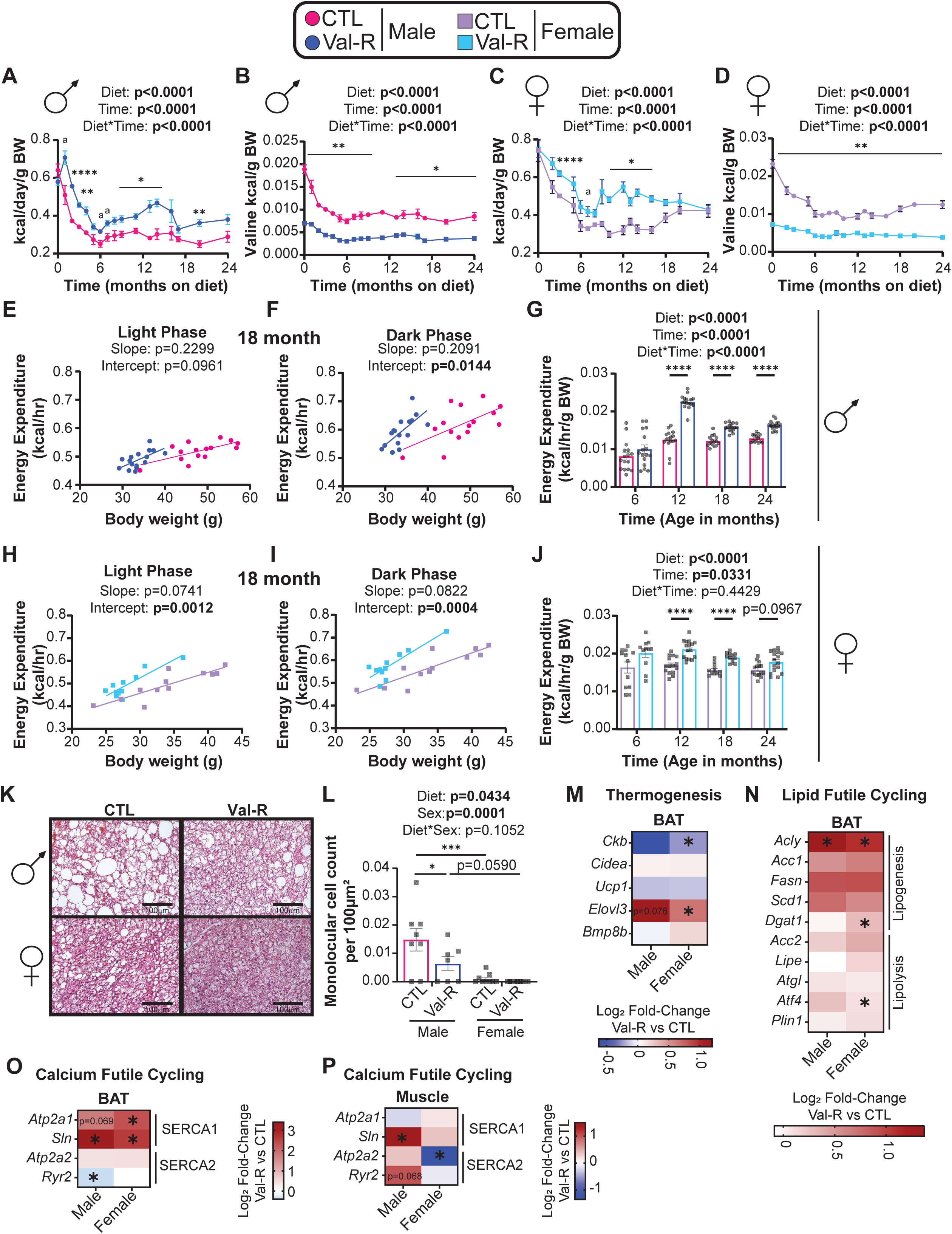
Val-R increases energy expenditure via the induction of thermogenesis in BAT. (A) Kilocalorie intake per day per gram of body weight (kcal/day/g BW) measured over 24 months on diet in males (n=4-20 mice/group). (B) Kilocalories derived from valine per gram of body weight (Valine kcal/g BW) over 24 months on diet in males (n=4-20 mice/group). (C-D) Kcal/day/g BW (C) and Valine kcal/g BW (D) over 24 months on diet in female mice (n=4-20 mice/group). (E-F) Energy expenditure of male mice as a function of body weight in the light (E) and dark (F) phase (n=15-16 mice/group). (G) Average energy expenditure normalized to body weight at 6, 12, 18 and 24 months of age in male mice (n=14-16 mice/group). (H-I) Energy expenditure of female mice as a function of body weight in the light (H) and dark (I) phase (n=11-12 mice/group). (J) Average energy expenditure normalized to body weight at 6, 12, 18 and 24 months of age in female mice (n=14-16 mice/group). (K-L) Representative images (K) and quantified number of monolocular cells per 100µm^2^ in BAT (L). (M-O) Log_2_ fold-change of gene expression from RNA sequencing analysis in the BAT of pre-selected genes relating to thermogenesis (M), lipogenesis and lipolysis (N) and SERCA1 and SERCA2 (O). (P) Log_2_ fold-change of gene expression from RNA sequencing analysis in the muscle of pre-selected genes related to SERCA1 and SERCA2. n=6-10 mice/group. (A-D, G, J) statistics for the overall effects of time, diet, and the interaction represent the p-value from a two-way ANOVA analysis. (E-F, H-I) data for each individual mouse is plotted; simple linear regression (ANCOVA) was calculated to determine if the slopes or elevations are equal; if the slopes are significantly different, differences in elevation cannot be determined. (L) statistics for the overall effects of sex, diet, and the interaction represent the p value from a two-way ANOVA. (A-D, G, J, L) a=p<0.1, *p<0.05, **p<0.01, ***p<0.001, ****p<0.0001 from a Sidak’s post-test examining the effect of parameters identified as significant in the two-way ANOVA. Data represented as mean ± SEM.

While decreased weight and adiposity can result from reduced calorie intake, this was not the case; in fact, Val-R-fed mice consume more calories (but less valine) relative to their body weight than CTL-fed mice (**Figs. 2A-D**). We therefore examined energy balance in detail using metabolic chambers. In both males and females, Val-R significantly increased energy expenditure at 18 months of age (**Figs. 2E-F & 2H-I**); we observed a similar overall effect of diet on energy expenditure throughout the lifespan, reaching statistical significance at 12, 18, and in males, 24 months of age (**Figs. 2G, 2J**). Although the CTL and Val-R diets have an identical percentage of calories derived from amino acids, carbohydrates, and fats, there was an overall effect of diet on the respiratory exchange ratio (RER) in both sexes, with RER higher in Val-R-fed males than CTL-fed males throughout their life, and RER higher in Val-R-fed females prior to 24 months of age (**Supplementary Figs. 3A-B**). The increased energy expenditure of Val-R-fed mice was not the result of increased activity; spontaneous activity was not altered by a Val-R diet in males and was lower, not higher, in Val-R fed females than in CTL-fed females (**Supplementary Figs. 3C-D**).

The effects of PR on energy balance are mediated by the energy balance hormone fibroblast growth factor 21 (FGF21), in part by promoting the beiging of inguinal white adipose tissue (iWAT) ^16, 64, 65, 66, 67, 68, 69^. However, we observed no increase in FGF21 in Val-R fed mice (**Extended Fig. 1A**). We also observed no change in the phosphorylation of eIF2α in the liver or muscle (**Supplementary Figs. 3E-L**), which is upstream of FGF21 ^70, 71, 72^, suggesting that an increase in FGF21 does not mediate the metabolic effects of Val-R. While we previously reported that Val-R induces beiging in young C57BL/6J mice ^22^, in these aged mice, consistent with the lack of induction of FGF21 in the present study, the iWAT adipocytes of Val-R-fed mice at 24 months of age were monolocular and of similar size to that of CTL-fed mice (**Extended Figs. 1B-C**). Despite this lack of change in the adipocyte morphology, there was an overall significant effect of diet on thermogenic gene expression, with a significant increase in the expression of *Elovl3* in Val-R-fed mice of both sexes (**Extended Figs. 1D-E**). We also saw an overall significant effect of diet effect on the expression of UCP1 protein, with a statistically significant increase in UCP1 in Val-R-fed males (**Extended Figs. 1F-G**). This suggests that Val-R may induce some changes that promote thermogenesis in iWAT.

To further understand how Val-R affects thermogenesis, we next turned to the brown adipose tissue (BAT). We found that Val-R-fed mice have a reduced number of monolocular cells in the BAT, which was significantly different in males (**Figs. 2K-L**), suggesting a shift away from lipid storage and towards utilization of lipids for thermogenesis. In support of this, we observed increased expression of *Elovl3* in the BAT along with an overall increase in lipogenesis-related genes with a modest increase in lipolysis-related genes in the BAT of both sexes (**Figs. 2M-N**), suggesting an upregulation of futile lipid cycling to promote thermogenesis. We also observed a significant increase in SERCA1-related genes, suggesting an upregulation of SERCA1-mediated futile calcium cycling in the BAT (**Fig. 2O**). This suggests that Val-R mice of both sexes also engage UCP1-independent mechanisms in the BAT to promote thermogenesis. Interestingly, we also observed a male-specific increase in *Sln* of SERCA1-mediated and a non-significant increase in *Ryr2* (p=0.068) of SERCA2-mediated futile calcium cycling in the muscle (**Fig. 2P**), suggesting that Val-R-fed males may engage muscle thermogenesis in addition to BAT thermogenesis to increase energy expenditure. Overall, this data suggests that the increased energy expenditure of Val-R fed mice is mediated by a multi-tissue increase in canonical and non-canonical thermogenesis.

We examined the effect of Val-R on glycemic control longitudinally throughout life. Val-R-fed males displayed improved glucose tolerance as early as 3 months of age and throughout life, even at 24 months of age (**Figs. 3A-C, Supplementary Figs. 4A-B, E-F, I-J, M-N**). Val-R-fed males tended to be more sensitive to I.P. administration of insulin than CTL-fed males, which was significant at 12 months of age (**Figs. 3D-F**). While we observed no significant difference in glucose-stimulated insulin secretion (GSIS) nor HOMA2-IR (**Figs. 3G-H**), we did observe an improvement in HOMA2 %B, suggesting Val-R improved pancreatic beta cell function (**Fig. 3I**). Additionally, we tested for suppression of hepatic gluconeogenesis by performing an alanine tolerance test (ATT). We found a significant overall effect of diet, suggesting that Val-R improves hepatic insulin sensitivity in males (**Supplementary Fig. 4Q**).

**Figure 3:**
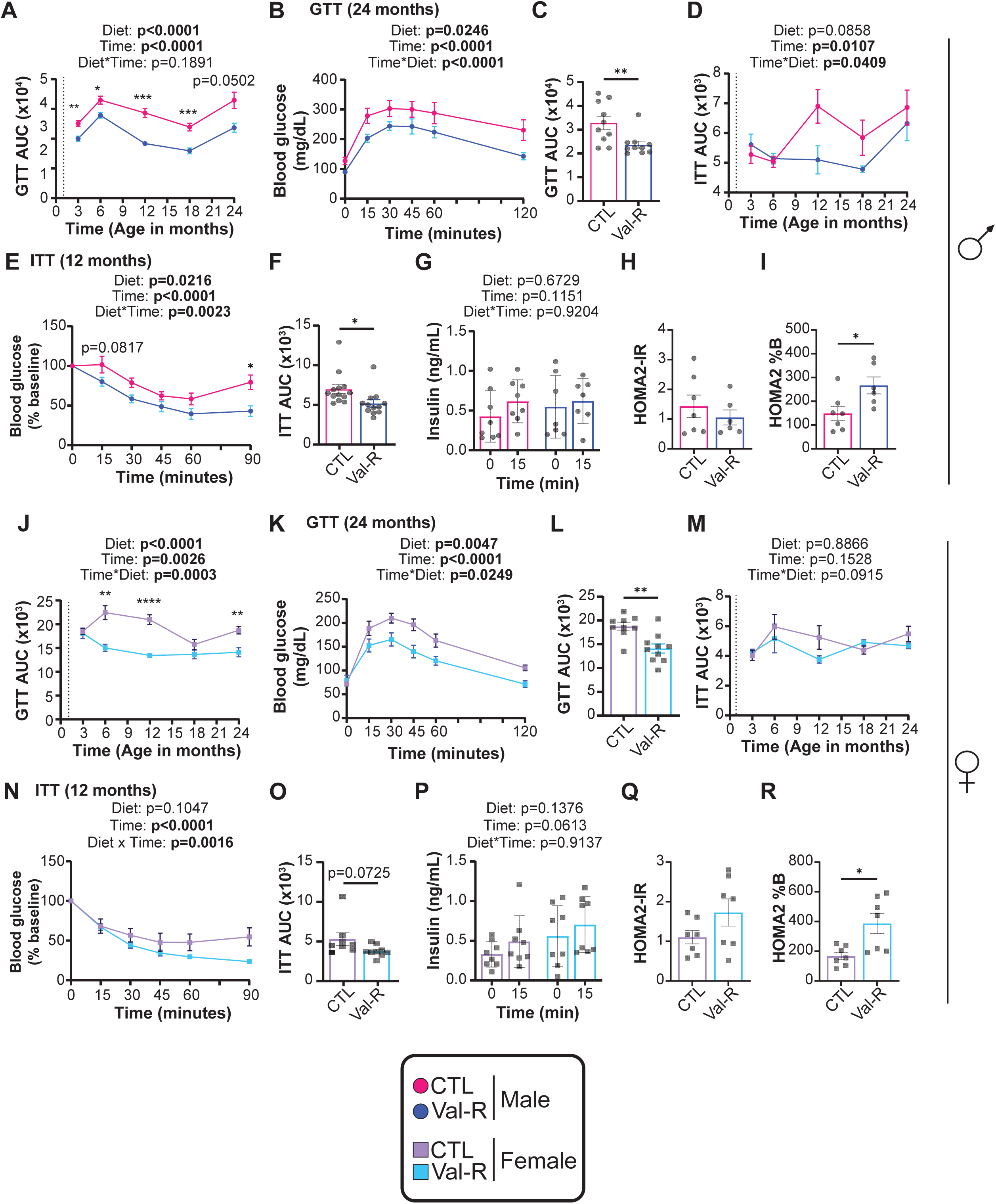
V**al-R improves glucose regulation.** (A) Glucose tolerance area under the curve (GTT AUC) over time in male mice (n=11-12 mice/group). (B-C) A glucose tolerance test (B) performed at 24 months of age and its GTT AUC (C) in male mice (n=11-12 mice/group). (D) Insulin tolerance area under the curve (ITT AUC) over time in male mice (n=9-13 mice/group). (E-F) An insulin tolerance test (E) performed at 12 months of age and its ITT AUC (F) in male mice (n=12-13 mice/group). (G) A glucose stimulated insulin secretion (GSIS) assay performed at 19 months of age (n=8 mice/group). (H-I) HOMA2-IR (H) and HOMA2 %B (I) calculated at 19 months of age in male mice (n=6-7 mice/group). (J) GTT AUC over time in female mice. (K-L) A GTT (K) performed at 24 months of age and its GTT AUC (L) in female mice (n=9-12 mice/group). (M) ITT AUC over time in female mice (n=8-12 mice/group). (N-O) An ITT (N) performed at 12 months of age and its ITT AUC (O) in female mice (n=9 mice/group). (P) A GSIS assay performed at 19 months of age in female mice (n=8 mice/group). (Q-R) HOMA2-IR (Q) and HOMA2 %B (R) calculated at 19 months of age in female mice (n=7-8 mice/group). (A-B, D-E, G, J-K, M-N, P) statistics for the overall effects of time, diet, and the interaction represent the p value from a two-way ANOVA analysis. (C, F, H-I, L, O, Q-R) Student’s t-test. (A-R) *p<0.05, **p<0.01, ***p<0.001, ****p<0.0001 from a Sidak’s post-test examining the effect of parameters identified as significant in the two-way ANOVA or student’s t-test. Data represented as mean ± SEM.

Val-R-fed females similarly had improved glucose tolerance throughout most of their life (**Figs. 3J-L, Supplementary Figs. 4C-D, G-H, K-L, O-P**). In contrast to males, there was no overall effect of Val-R on the response to I.P. administration of insulin (**Figs. 3 M-O**). As in males, the Val-R diet had no significant differences in GSIS nor HOMA2-IR in females, but increased beta cell function as measured via HOMA2 %B (**Fig. 3P-R**). Also as in males, a Val-R diet had an overall effect of diet on alanine tolerance suggestive of improved hepatic insulin sensitivity (**Supplementary Fig. 4R**). Overall, we found that Val-R-fed mice of both sexes had improved glycemic control relative to their CTL-fed counterparts.

Hepatic insulin sensitivity is influenced by hepatic lipid deposition. At 24 months of age, we assessed fatty liver using Oil-Red-O staining. We found that there was an overall effect of diet on lipid droplet size in both sexes, with Val-R-fed males having significantly smaller hepatic lipid droplets than CTL-fed males (**Extended Figs. 2A-B**). While the overall area of lipids droplets decreased in response to a Val-R diet (**Extended Fig. 2C**), the total number of lipid droplets increased in both sexes (**Extended Fig. 2D**). We observed an overall effect of diet on hepatic and plasma triglycerides in both sexes, with Val-R-fed males having significantly less hepatic triglycerides and significantly more plasma triglycerides than CTL-fed males (**Extended Figs. 2E-F**). This signature of reduced hepatic triglycerides with increased circulating triglycerides is consistent with normal, healthy conditions in which the liver stores little triglycerides and exports the fatty acids to be used by other tissues, such as the muscle and adipose tissue ^73^.

Overall, we find that consumption of a Val-R diet results in improved metabolic health. Val-R reduces fat mass and adiposity, likely through an increase in energy expenditure mediated by increased BAT thermogenesis, improves glycemic control, and protects from hepatic steatosis.

### Tissue- and sex-specific molecular effects of valine restriction

To investigate the molecular impact of Val-R across tissues, we performed transcriptional profiling of brown adipose tissue (BAT), liver and muscle of CTL-fed and Val-R-fed mice of both sexes at 24 months of age. As anticipated, principal component analysis (PCA) showed that gene expression profiles grouped strongly by tissue type (**Fig. 4A**). We visualized the top 50 most variable genes across samples and found that, overall, the muscle and BAT had much more similar gene expression patterns than the liver, with the exception of *Ucp1* which was much more highly expressed in BAT than in liver or muscle (**Fig. 4B**).

**Figure 4:**
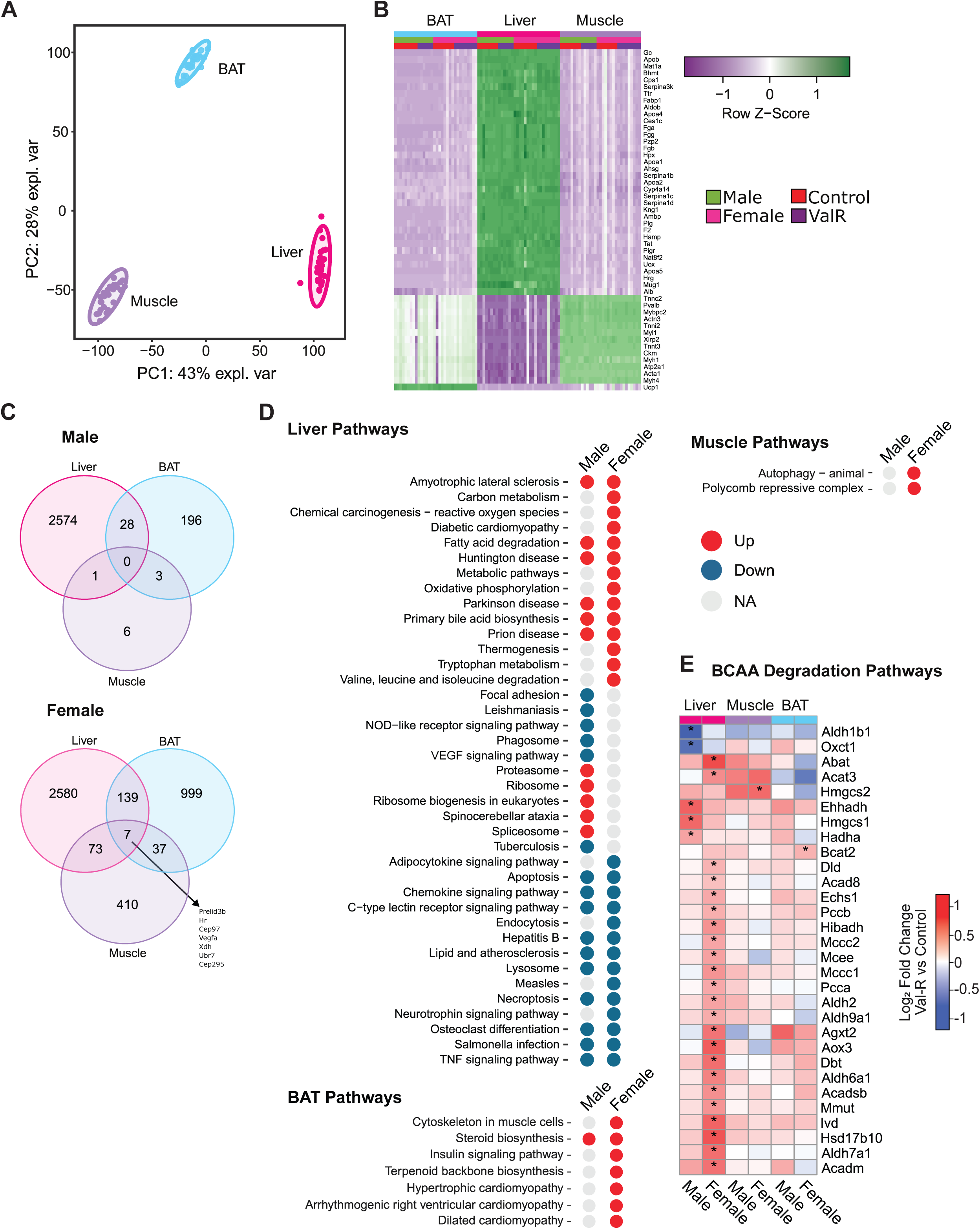
Multi-tissue transcriptomic analysis of male and female mice on Val-R. (A) A principal component analysis (PCA). (B) Top 50 differentially expressed genes (DEGs) in the liver, muscle and adipose tissue. (C) Venn diagram of number of gene changes by tissue in male and female mice. (D) KEGG pathway analysis in liver, muscle and BAT in male and female mice. (E) Altered genes in the BCAA degradation pathway in the liver, muscle and BAT. (A-E) n=6-10 mice/group; *p<0.05, moderated t-test.

The response to Val-R was highly influenced by sex and tissue. Liver had by far the greatest number of differentially expressed genes in response to Val-R, with over 2,500 genes differentially expressed in both sexes. There were nearly 1,000 genes differentially expressed in Val-R-fed female BAT, but only 196 were differentially expressed in Val-R-fed male BAT. Similarly, over 400 genes were differentially expressed in Val-R-fed female muscle, but only 6 genes were differentially expressed in Val-R-fed male muscle. In males, no genes were significantly differentially expressed in response to Val-R across all three tissues, while in females, only seven genes were shared between liver, BAT and muscle (**Fig. 4C**).

When we investigated the pathways enriched for each of the tissues, and compared males and females, we found many pathways altered by Val-R in the liver, with fewer pathways altered in muscle and BAT, and a strong sex-specific response in those tissues (**Fig. 4D**). In liver, there was an upregulation of the “Valine, leucine, and isoleucine degradation” pathway in females in Val-R mice. We further examined this pathway at the gene level; interestingly, while we see an overall upregulation of this pathway across both males and females in all tissues, the strongest changes, as well as the most sex-specific, occurred in liver, with more modest effects in muscle and BAT (**Fig. 4E**).

In the liver, there was an upregulation of “Fatty acid degradation” and “Primary bile acid biosynthesis” in both male and female Val-R-fed mice. Other upregulated liver pathways in both sexes included “Amyotrophic Lateral Sclerosis,” “Parkinson’s disease,” “Prion disease,” and “Huntington’s disease.” Several immune pathways, including the chemokine, TNF and C-type lectin receptor signaling pathways as well as “Salmonella infection” were downregulated in the livers of both sexes. Other non-immune downregulated pathways in both sexes include “Apoptosis,” “Hepatitis B,” “Lipid and atherosclerosis,” “Lysosome,” “Necroptosis,” and “Osteoclast differentiation.” There were also numerous changes that were sex-specific, with Val-R-fed females having an upregulation of several metabolic pathways, such as “Oxidative phosphorylation,” “Thermogenesis,” “Carbon metabolism,” and “Tryptophan metabolism;” there was also an upregulation of “Reactive Oxygen Species” and “Diabetic cardiomyopathy”. Val-R-fed females also displayed a downregulation of the “Adipocytokine signaling pathway.” In male livers, we found an upregulation of “Ribosome” and “Ribosome biogenesis in eukaryotes” as well as a downregulation of “VEGF signaling” and several immune related pathways (**Fig. 4D**).

In the muscle, only “Autophagy” and “Polycomb repressive complex” was upregulated in females in response to Val-R, while in the BAT, several metabolic pathways including “Steroid biosynthesis” in both sexes and “Insulin signaling pathway” in the females were upregulated (**Fig. 4D**). Most pathways altered in muscle and BAT were altered only in females. As noted in **Fig. 4E**, we observed altered expression of several genes involved in BCAA degradation in all three tissues, with the most significant changes in the livers.

Amino acids in general and the BCAAs in particular are agonists of mTORC1; we have previously shown that protein or BCAA restriction reduces mTORC1 signaling in the muscle and liver as well as other tissues ^23, 42, 74, 75^. While one of the pathways identified from transcriptional profiling was consistent with decreased mTORC1, i.e as downregulation of “Lysosome” in the liver, the male-specific increase in “Ribosome and Ribosome biogenesis in Eukaryotes” (**Fig. 4D**), suggested that hepatic mTORC1 signaling might actually increase in Val-R-fed males.

To clarify the effect of Val-R on mTORC1, we first examined the genes in the “Ribosome” and “Ribosome biogenesis in eukaryotes” pathways. We saw a significant increase of most ribosomal genes in the livers of male Val-R-fed mice as well as several genes in the livers of the Val-R-fed females (**Extended Figs. 3A-B**). In agreement with this, at the protein level, we found a significant effect of diet on mTORC1 signaling in both males and females, with a statistically significant increase in the phosphorylation of the mTORC1 substrate S6K1 T389 in both sexes (**Extended Figs. 3C-D, F-G**). In the livers of males, but not females, we found an overall significant effect of Val-R on autophagy, with a non-significant increase in the level of LC3 and LC3 cleavage (p=0.0987) (**Extended Figs. 3E and 3H**). These results agree with our transcriptional profiling of the liver, and collectively, suggests that despite the decrease in dietary valine, a Val-R diet increases the activity of mTORC1 in the liver.

In the muscle, there were several upregulated ribosomal genes in the Val-R females (**Extended Figs. 3A-B**). However, in contrast to the liver, when we looked at mTORC1 and autophagy at the protein level, there was no statistically significant effect of diet in either pathway in male muscle (**Supplementary Figs. 5A-C**). In females, we saw a trend (p=0.0594) towards reduced mTORC1 signaling and no significant difference in autophagy in Val-R fed females (**Supplementary Figs. 5D-F**). We did see a slight increase in LC3AB abundance in females, but it was not significant; however, this may explain the increase in autophagy in our female muscle transcriptomic data.

### Valine restriction reduces neuroinflammation in both sexes and improves short-term memory in females

Restriction of dietary protein or BCAAs preserves cognition in the context of Alzheimer’s disease ^42, 76, 77^. To assess the effects of Val-R on cognitive function during normal aging, we performed a novel object recognition (NOR) test on the mice at 27 months of age. We found that there was a diet-sex interaction (p=0.0543) on short-term memory, with Val-R fed females, but not males, showing an improved ability to recognize the novel object compared to their CTL-fed counterparts (**Fig. 5A**). In contrast, neither sex displayed improvements in long-term memory with Val-R feeding (**Fig. 5B**).

**Figure 5:**
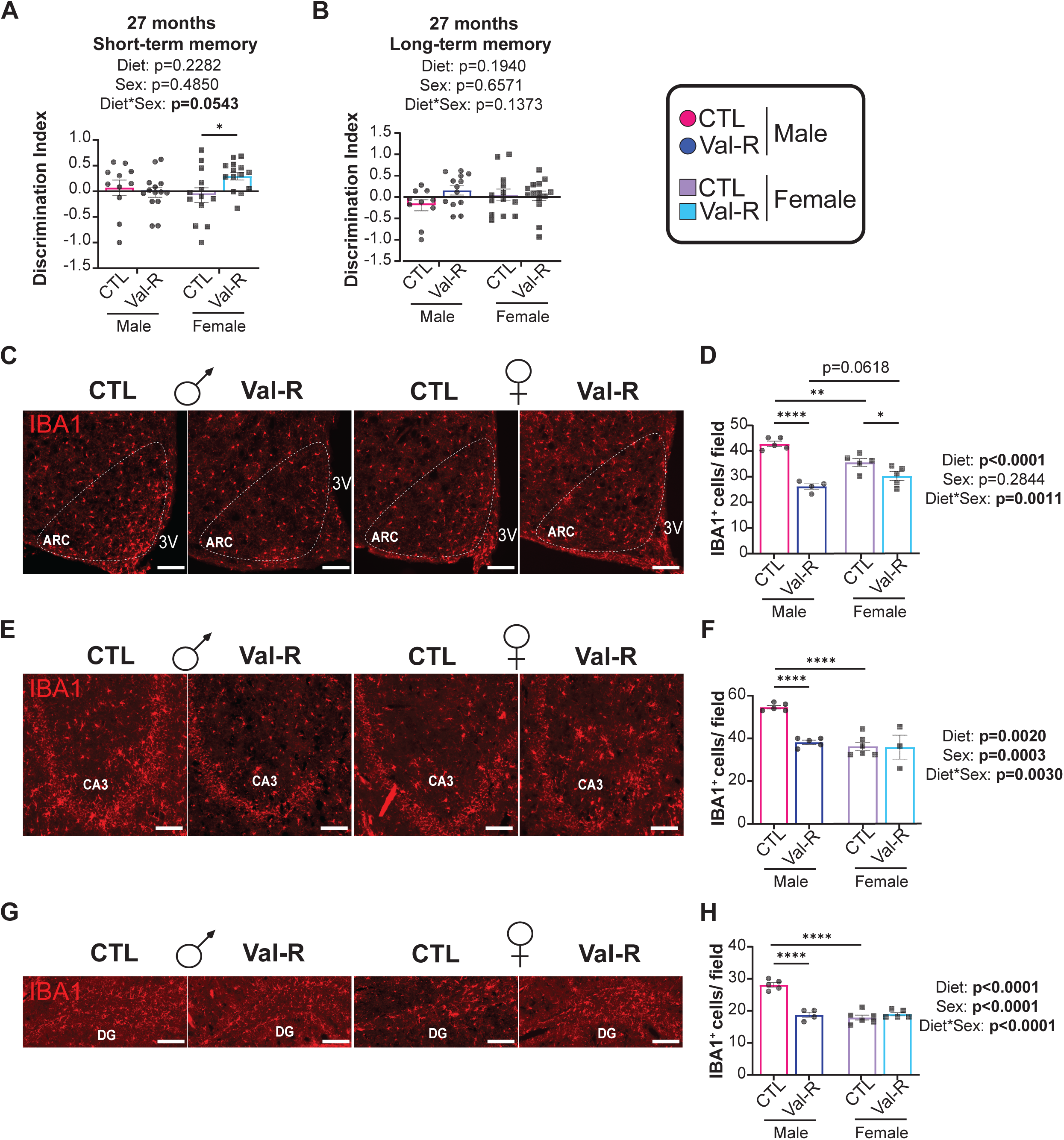
Val-R improves cognition in female mice and reduces neuroinflammation in Val-R-fed males and females. (A-B) Novel object recognition test discrimination index in the short-term memory test of males and females (A), and the long-term memory test in males and females (B) (n=10-14 mice/group). (C, E, G) Representative images of Iba1 staining in the Arc, CA3 and DG of the brain. Scale bar represents 200 µm. (D, F, H) Quantified staining of Iba1 in the Arc (D), CA3 (F) and DG (H) (n=3-6 mice/group). (A-B, D, F, H) statistics for the overall effects of sex, diet, and the interaction represent the p value from a two-way ANOVA analysis; *p<0.05, **p<0.01, ****p<0.0001 from a Sidak’s post-test examining the effect of parameters identified as significant in the two-way ANOVA. Data represented as mean ± SEM.

Neuroinflammation commonly increases with age and is associated with cognitive decline as well as age-related diseases such as Alzheimer’s disease ^78, 79, 80, 81^. We assessed neuroinflammation by quantifying the density of microglia and astrocytes through immunostaining brain sections with anti-glial fibrillary acidic protein (GFAP), an astrocyte marker, or anti-ionized calcium binding adaptor molecule 1 (IBA-1), a microglia marker. We looked at the arcuate nucleus (Arc) of the hypothalamus, a region critical for energy homeostasis as well as the sub-regions of the hippocampus, the CA3 region and the dentate gyrus (DG), which play a role in memory acquisition and memory retrieval. We found that Val-R reduces IBA1 in all three regions of the brain in male mice (**Figs. 5C-H**), and reduced GFAP in the Arc and CA3 of Val-R males (**Supplementary Figs. 6A-F**). Val-R also reduced IBA1 in the Arc of females, and significantly reduced GFAP in the DG of females (**Figs. 5C-D and Supplementary Figs. 6C-D**). However, this is most likely due to the significant overall effect of sex on IBA1 staining of CA3 and DG, and GFAP staining of all regions, due to the substantially lower levels of IBA and GFAP in females than males (**Figs. 5C-H and Supplementary Figs. 6A-F**). We also saw a significant diet*sex interaction in IBA1 staining of the Arc, which we interpret as the effect of Val-R being greater in males than in females (**Figs. 5C-H**).

To further assess the neuroinflammatory state of the brain, we performed skeletal morphological analysis of the microglia and astrocytes in the hippocampus as structural indicators of activation/reactivity ^46, 47^. Activation of microglia or reactivity of astrocytes is associated with chronic, damaging neuroinflammation that disrupts neuronal homeostasis and impairs brain function ^82, 83, 84^. Using the microglia skeletal analysis, we found a diet*sex interaction on process length and bounding circle diameter (**Extended Figs. 4A-F**), with a Val-R diet increasing these in males but not females, suggesting that a Val-R diet promotes an overall more ramified or resting state in males only. Females in general displayed shorter process lengths and reduced fractional dimension compared to males, along with increased lacunarity, (greater structural gaps) in Val-R-fed conditions (**Extended Figs. 4A-F**), suggesting that females as a whole may have somewhat more activated microglia than males. Using the astrocyte skeletal analysis, we found a diet*sex interaction for total number of endpoints and process lengths, with a Val-R diet reducing these in male, but not female mice (**Supplementary Figs. 6G-K**). These reductions are consistent with decreased astrocytic hypertrophy and reactivity in males. Overall, we find that a Val-R diet has sex-specific effects on glial morphology, supporting function and reduced neuroinflammatory tone selectively in males.

#### Valine restriction improves healthspan and lifespan

We comprehensively assessed healthspan in mice as they age. Mice and humans become increasingly frail with age, and starting at approximately 12 months of age, we utilized a widely-adopted mouse frailty index ^39^ to assess the impact of Val-R on frailty in both sexes. We observed that in both males and females, Val-R feeding resulted in a lower frailty index as compared to their CTL-fed counterparts at multiple timepoints across age (**Figs. 6A-B**). The reduced frailty in Val-R fed mice was primarily the result of decreased deficits in the “Physical/Musculoskeletal” and “Discomfort” categories (**Supplementary Figs. 7A-L**).

**Figure 6:**
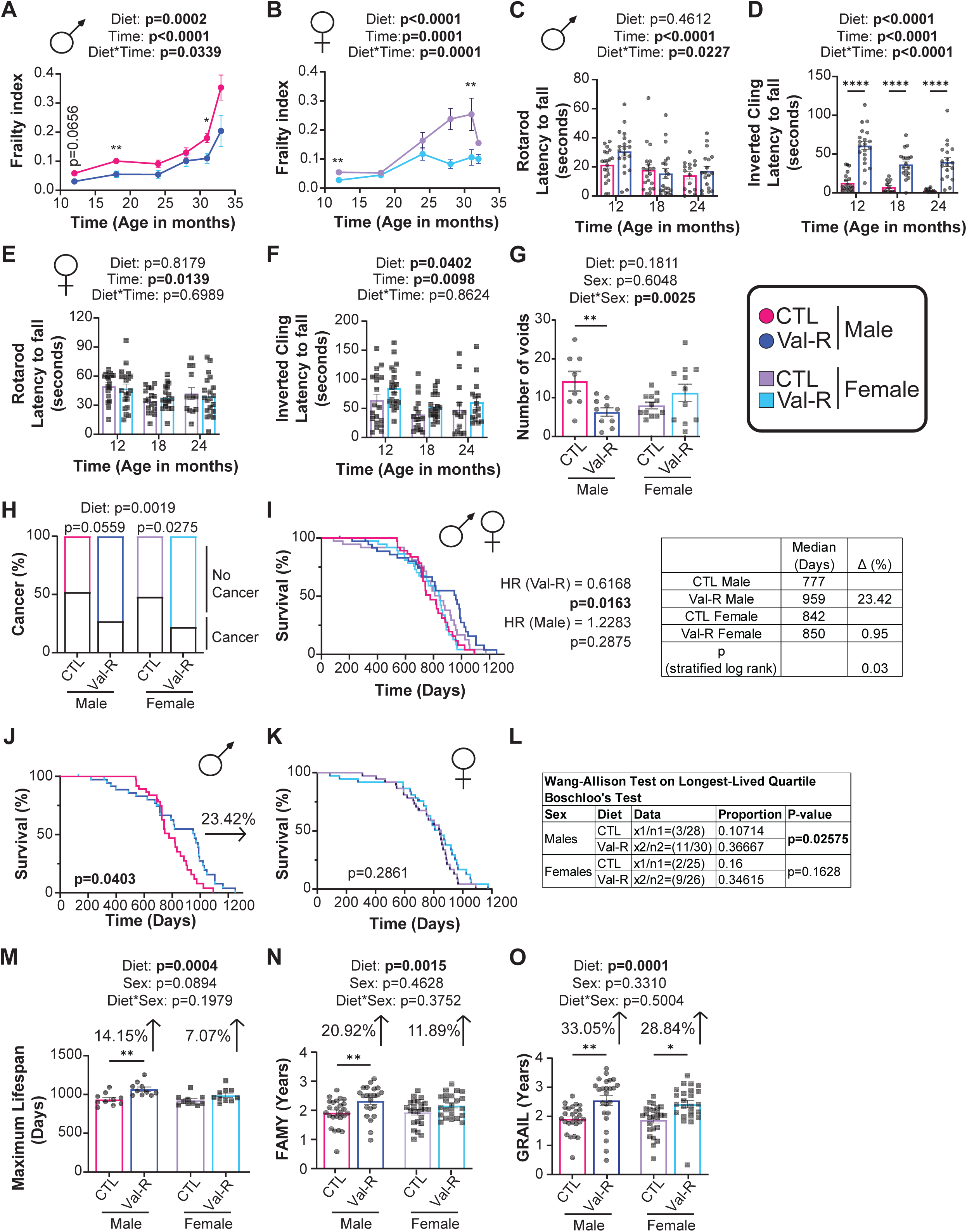
Val-R improves health of both males and females and extends lifespan in male mice. (A-B) Frailty index scoring of male (A) and female (B) mice from 12 to 32 months of age in mice. (C) Rotarod latency to fall in seconds at 12, 18 and 24 months of age in male mice (n=25 mice/group). (D) Inverted cling latency to fall in seconds at 12, 18 and 24 months of age in male mice. (E) Rotarod latency to fall in seconds at 12, 18 and 24 months of age in female mice (n=15-20 mice/group). (F) Inverted cling latency to fall in seconds at 12, 18 and 24 months of age in female mice (n=15-20 mice/group). (G) Void spot assay conducted in 27-month-old male and female mice (n=8-12 mice/group). (H) Percent of male and female mice with and without cancer observed at necropsy from lifespan in (I-K), Statistic from Fischer’s exact test. (I) Kaplan-Meier plots showing the survival of male and female (n=37 biologically independent animals for both groups). Table of median lifespans for mice on each diet, percent change between Val-R-fed and Control-fed mice of the same sex, and the two-sided Gehan-Breslow-Wilcoxon p-value between Val-R-fed and CTL-fed mice of the same sex. (J) Kaplan-Meier plots showing the survival of male (n=37 biologically independent animals for each diet). (K) Kaplan-Meier plots showing the survival of female (n=37 biologically independent animals for each diet). (L) Table of maximum lifespan calculations were made by generating a cutoff of the top 25% longest lived animals in each sex, coupled with Boschloo’s Test (Wang-Allison) for significance testing between groups. (M) Maximum lifespan of top 10 longest lived mice per group per sex (n=10 mice/group). (N) Frailty-Adjusted Mouse Years (FAMY) in years was calculated using the survival and frailty data plotted in panels A-B and J-K (n=24-25 mice/group). (O) Gauging Robust Aging when Increasing Lifespan (GRAIL) in years calculated using the survival and frailty data plotted in panels A-B and J-K (n=24-25 mice/group). (A-G, M-O) statistics for the overall effects of time or sex, diet, and the interaction represent the p value from a two-way ANOVA analysis; *p<0.05, **p<0.01, ***p<0.001, ****p<0.0001 from a Sidak’s post-test examining the effect of parameters identified as significant in the two-way ANOVA. Data represented as mean ± SEM.

We also performed rotarod and inverted cling assays to assess muscle coordination and grip strength, respectively. In both males and females, we found no difference in muscle coordination with Val-R feeding as measured by rotarod performance (**Figs. 6C, 6E, Supplementary Figs. 8A-C, E-F**), while age did seem to reduce performance regardless of diet. In Val-R-fed males, we observed a significant overall improvement in grip strength as measured by an inverted cling assay, which reached statistical significance at every age measured (**Fig. 6D**). Analyzing performance on these assays using weight as a covariate, however, suggests the effects of Val-R on inverted cling performance are primarily the effect of lower weight (**Supplementary Figs. 8A-F**). In Val-R-fed females, we saw a significant overall impact of diet on inverted cling performance that did not reach significance at any single age (**Fig. 7F**). When analyzed by ANCOVA with weight as a covariate, Val-R-fed females performed worse on the rotarod than CTL-fed females at all time points, reaching statistical significance at 12 and 24 months of age, and Val-R-fed females had slightly worse, though not significantly so, inverted cling performance than CTL-fed females at all ages (**Supplementary Figs. 8G-L**). Additionally, when using ANCOVA to test for body weight as a covariate for inverted cling performance, we found that this test could not be performed at 24 months of age in males because their slopes reached statistical significance, and therefore, could not be compared (**Supplementary Figs. 8E-F**). To address the issue of comparing the groups using ANCOVA, we normalized cling time by body weight. We found that Val-R-fed males, but not females, had increased cling time normalized to body weight at every timepoint (**Supplementary Figs. 8M-N**). We still observed an overall effect of diet in both sexes (**Supplementary Figs. 8M-N**). Overall, these results are consistent with Val-R having minimal effects on muscles, except perhaps promoting strength in aged mice.

**Figure 7:**
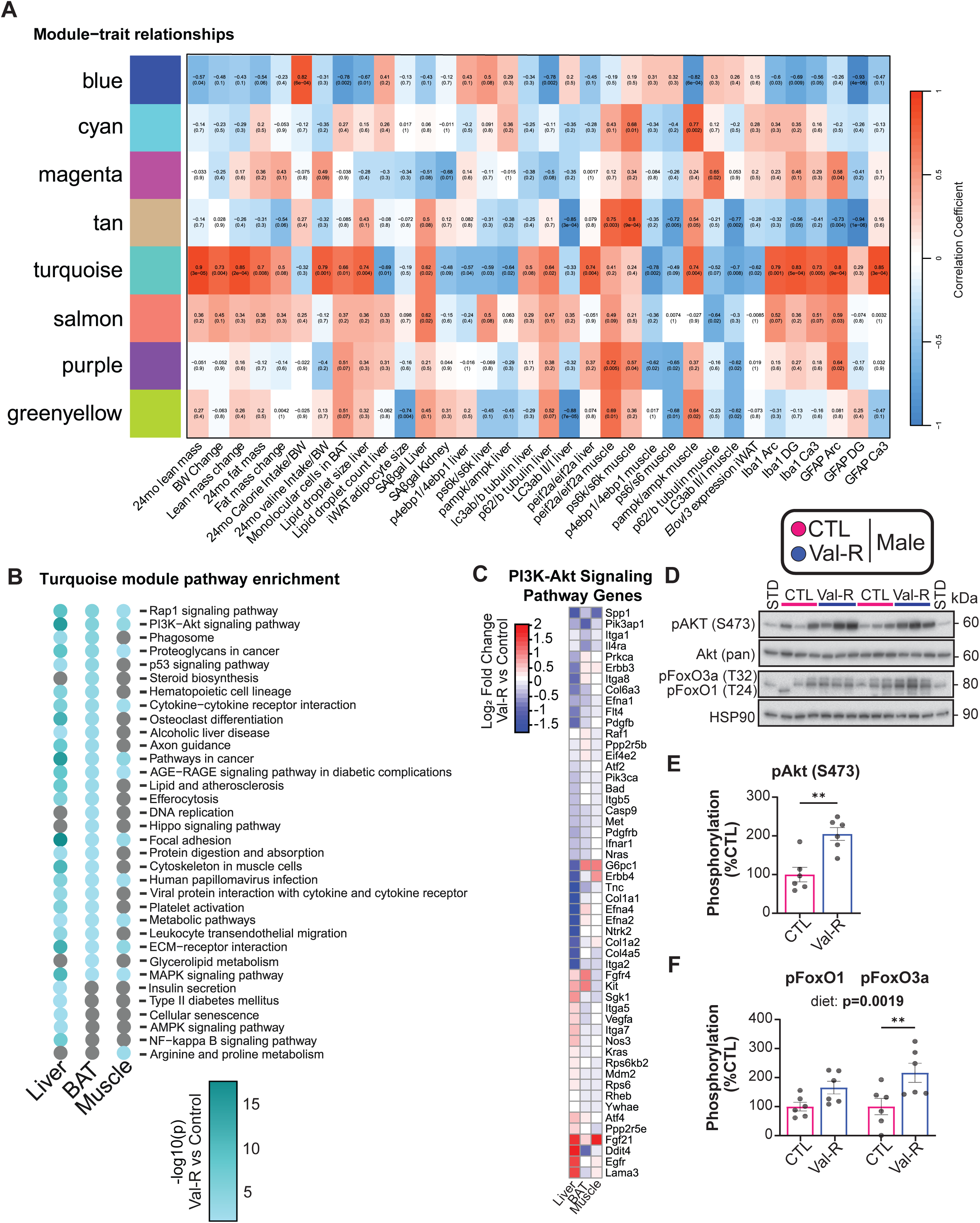
WGCNA analysis of selected modules and pathways in males. (A) Pearson correlation coefficient between the gene modules and selected phenotypic traits, numbers in brackets indicate the corresponding p values in males (n=6-9 mice/group). (B) Selected KEGG pathway enrichment of turquoise module. Gray dots indicate no alterations in that pathway for that tissue. (C) Altered genes in the PI3K-AKT signaling pathway in the liver, muscle and BAT. Genes shown were significantly altered (Benjamini-Hochberg (BH) adjusted p<0.05) by Val-R in at least one tissue in males or females. (D) Western blots of the analyzed proteins in male livers. (E) Phosphorylation of AKT S473 normalized to the expression of AKT. (F) Phosphorylation of FoxO1 and FoxO3a normalized to HSP90. (E-F) n=6 mice/group; *p<0.05, t-test. Data represented as mean ± SEM.

Lower urinary tract dysfunction increases with age ^85, 86^. We assessed urinary frequency, and we found that Val-R-fed males had significantly less urinary spotting compared to their CTL-fed counterparts (**Fig. 6G**); no difference was observed in females (**Fig. 6G**).

Cancer is a major cause of death in C57BL6/J mice ^87, 88, 89^. Upon natural death or meeting criteria for euthanasia, we performed a gross necropsy in mice followed to the end of their life. We found that when we pooled males and females together (as there was no significant effect of sex), there was a significant overall effect of diet, with Val-R fed mice having a lower prevalence of cancer observed at necropsy. Assessing the effect of diet in each sex, we found the prevalence of cancer observed was reduced significantly in female Val-R mice, and non-significantly (p=0.0559) in male Val-R mice (**Fig. 6H**). Most of the cancers observed were most likely hepatocellular carcinoma, or lymphoma and leukemia as indicated by splenomegaly, but were not formally assessed by a pathologist.

Lastly, in agreement with the reduction in frailty, improved health metrics and reduced cancer incidence, we found that the lifelong consumption of a Val-R diet extends lifespan (p=0.03, stratified log-rank test) when considering the effect on both sexes (**Fig. 6I**). Cox regression likewise indicated a significant effect of diet on survival (hazard rate (HR) = 0.0163), and no interaction of diet with sex was detected (**Fig. 6I**). When assessing the sexes separately, we found a Val-R diet increased the median lifespan of males by 23.42% (log-rank, p=0.0403), but not females; we similarly observed a significant extension of the maximum lifespan of males (Wang-Allison, p=0.02575), but not females by a Val-R diet (**Fig. 6I-L**). Investigating the survival of the top 10 longest-lived animals in each group, we found the top 10 Val-R males had a statistically significant 14.5% increase in maximal lifespan relative to the top 10 of CTL-fed males; the top 10 Val-R-fed females had a non-significant 7.07% increase in maximum lifespan compared to the top 10 CTL-fed females (**Fig**. **6M**).

Lastly, using the cumulative data on longevity, frailty, healthspan and hallmarks of aging collected during this lifespan study, we assessed FAMY (Frailty Adjusted Mouse Years) and GRAIL (Gauging Robust Aging when Increasing Lifespan), new summary statistics that are analogous to Quality Adjusted Life Years (QALY) in humans ^63^. In response to Val-R, we see increases in FAMY and GRAIL in males, with a 20.29% increase in FAMY and a 33.05% increase in GRAIL (**Fig**. **6N-O**). In females, while we do see an 11.89% increase in FAMY, it is not statistically significant; however, GRAIL increases by 28.84% (**Fig**. **6N-O**). Overall, these data demonstrate that Val-R increases healthspan in both males and females.

We also assessed a key hallmark of aging: cellular senescence ^90, 91^. We hypothesized that with the improved healthspan of both sexes and male-specific lifespan extension, we would see a reduction in senescence. Staining for the cellular senescence marker SA-β-Gal, we found that in response to Val-R, there was a significant diet effect in the liver, as well as an overall reduction in senescence and senescence-associated secretory phenotype (SASP) genes (**Extended Figs. 5A-C**). The genes we chose were based on CoreScence genes, which is a core senescence gene set that generalizes across species, tissues, and senescence contexts, as well as uses senescence signatures that are expressed in at least 5 senescence data sets, such as SenMayo and SenSig ^92, 93, 94^. We similarly found a significant effect of diet in the kidney, with a diet*sex interaction consistent with a reduction of cellular senescence by Val-R in females but not males as assessed by SA-β-Gal, as well as a downregulation of senescent and SASP genes (**Extended Figs. 5D-F**). We also surveyed senescence and SASP genes in several other tissues; we found an overall downregulation of these genes in both sexes in iWAT and BAT, and specifically in males in eWAT (**Extended Figs. 5G-I**). Lastly, we found no consistent change in senescence and SASP genes in muscle (**Extended Fig. 5J**); however, this may be due to the small fraction of proliferation-competent cells in muscle.

In summary, we find that lifelong Val-R results in both general and sex-specific improvements in both health and lifespan. Val-R reduces body weight and adiposity, which may be mediated by an increase in energy expenditure, as well as promoting blood sugar control across the lifespan. We see that Val-R reduces hepatic steatosis in both sexes, but particularly in males. We found that Val-R improves cognition in females while reducing neuroinflammation in both sexes and reducing activation of microglia and reactivity of astrocytes in the male hippocampus. Val-R reduces frailty, cancer prevalence, and senescence in both sexes as well as improves urinary function in males. Val-R extends median lifespan by 23.42% and maximum lifespan by 14.15% in males, while not significantly increasing median or maximum lifespan in females. We also confirmed an overall improvement in healthspan in response to Val-R in both sexes through the assessment of FAMY and GRAIL.

### Val-R upregulates hepatic mitochondrial respiration

To gain more mechanistic insight into the effects of Val-R, we performed a weighted gene co-expression network analysis (WGCNA). Selected traits and pathways for males and females are shown in **Fig. 7** and **Extended Fig. 6**; full tables and pathways can be found in **Tables S9-S12**. In males, we observed that the turquoise module had several strongly correlated phenotypes, including body composition, valine intake, hepatic lipid droplet size and brain inflammation traits (**Fig. 7A**). Pathway enrichment on the genes in the turquoise module showed that the genes in this module were enriched for changes in processes related metabolism and longevity pathways, such as the “PI3K-Akt pathway”, “MAPK signaling”, “AGE-RAGE signaling pathway in diabetic complications” and “metabolic pathways” (**Figs. 7B**). We investigated the significantly altered genes within the PI3K-Akt and MAPK signaling pathways, as both of these pathways have been shown to regulate longevity. There was not a clear effect of Val-R on gene expression of MAPK signaling genes in any tissue, nor was there an effect on the phosphorylation of MAPK/ERK in the livers of either sex (**Supplementary Fig. 9A-F**). However, we did observe an overall downregulation of genes in the PI3K-Akt signaling pathway in males across tissues (**Fig. 7C**), suggesting a potential role in the downregulation of PI3K-Akt signaling in the lifespan extension in males.

In females, we observed that the blue module had a strong negative correlation with autophagy and a positive correlation with lean mass (**Extended Fig. 6A**). When we performed pathway enrichment on the genes clustered into the blue module, we found only 3 pathways that had changes in all three tissues (“hematopoietic cell lineage”, “intestinal immune network for IgA production” and “cytokine-cytokine receptor interaction”) that do not have known associations with aging (**Extended Fig. 6B**). However, we saw that the “PI3k-Akt signaling” pathway was also enriched in liver and muscle. We found largely no effect on genes in the PI3K-Akt pathway in response to Val-R in females (**Extended Fig. 6C**).

Because the liver showed the greatest changes in the PI3K-Akt signaling pathway, we looked at the phosphorylation of AKT S473; surprisingly, despite the decreased expression of many pathway genes, hepatic AKT S473 phosphorylation was increased in Val-R males (**Figs. 7D-E**). The phosphorylation of the AKT substrates FOXO1 and FOXO3a were also increased, supporting an increase in AKT activity (**Figs. 7D, 7F**). The increased hepatic mTORC1 we observed previously can likely be explained by the increase in AKT activity, as mTORC1 is activated by PI3K-AKT activity. In females, although there was a trend towards increased phosphorylation of AKT S473 (p=0.0749), there was no increase in the phosphorylation of FOXO1 or FOXO3a (**Extended Figs. 6D-F**), further confirming no change in the PI3K-AKT pathway in females. Overall, this data suggests that the longer lifespan of males is not the result of downregulated PI3K-AKT signaling.

To determine what may be mediating the lifespan extension we see in males, we used WGCNA modules and correlated the module eigengene (ME) for each distinct gene module with other module MEs, and used these correlations to construct a network in Cytoscape where the nodes are MEs and edges correlaDtion between nodes (**Fig. 8A**). This analysis allows for the identification of a root node, a cluster of genes that serves as a central hub connecting other gene clusters (i.e. nodes) within and across tissues. Interestingly, we found that the module “L.Yellow” (i.e., Liver yellow) serves as the root node, a cluster of genes that serves as a central hub connecting other nodes within and across tissues, in our data set (**Fig. 8A**). KEGG pathway enrichment revealed that many of the L.Yellow module genes relate to the mitochondria, including such pathways as “Fatty acid degradation,” “Fatty acid metabolism,” “Glyoxylate and dicarboxylate metabolism,” “Chemical carcinogenesis − reactive oxygen species,” and “Pyruvate metabolism” (**Fig. 8B**). To further assess the mitochondrial metabolism in the liver, we assessed genes related to mitochondrial regulation, electron transport chain (ETC), fatty acid metabolism and TCA cycle (**Supplementary Fig. 10**). Interestingly, while we found changes in both sexes when it comes to β-Oxidation and the TCA cycle, we found a female specific increase nuclear-encoded genes related to the ETC.

**Figure 8:**
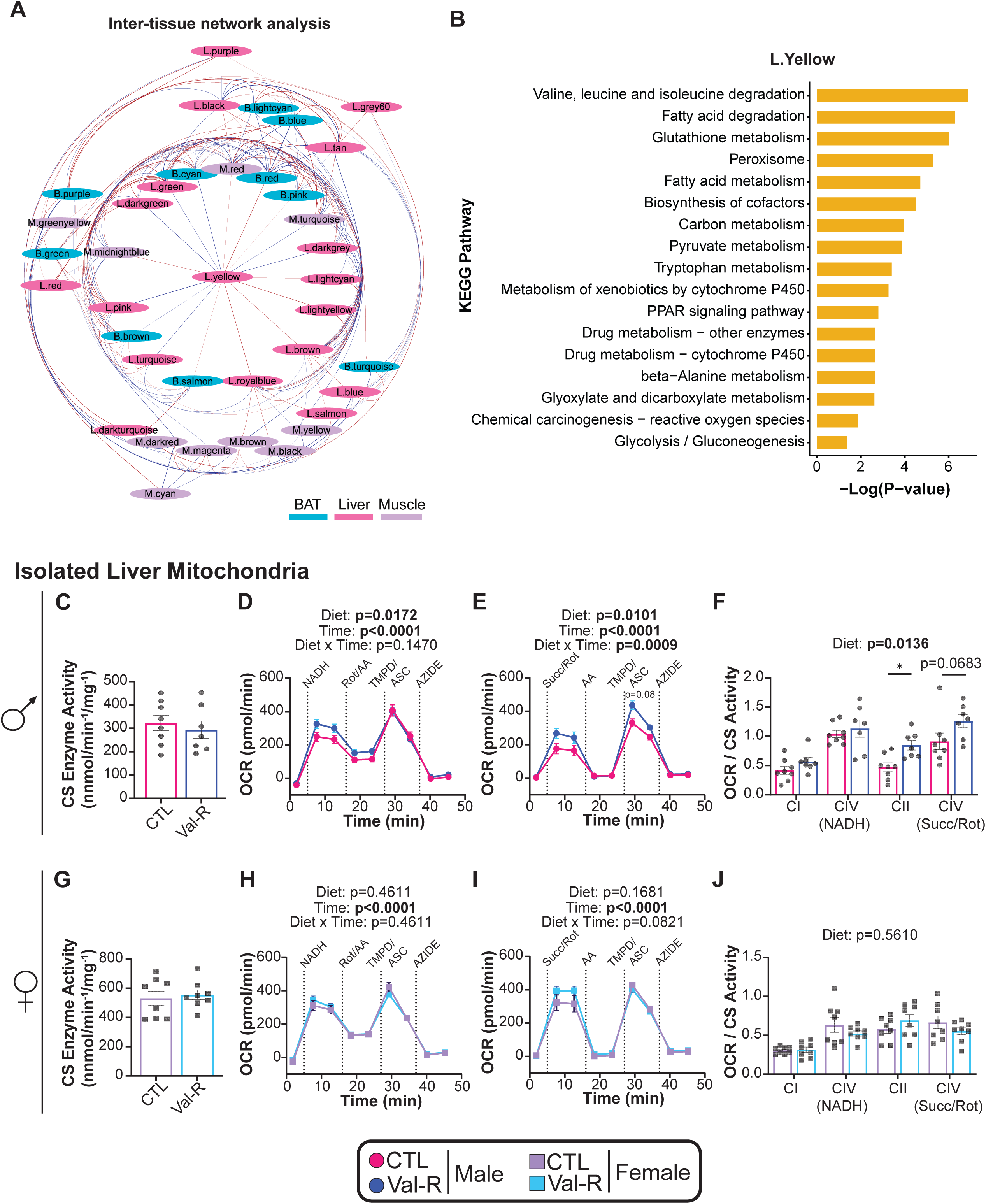
Val-R alters mitochondrial metabolism and respiration. (A) ME-ME correlations were used to construct a correlation network in Cytoscape. Color indicates direction of correlation (red = positive, blue = negative), and thickness is proportional to the strength of correlation (|correlation| range, 0.37 to 1.0). (B) KEGG pathway enrichment of the L.Yellow Node. (C) Citrate synthase (CS) enzyme activity in male isolated liver mitochondria. (D) Oxygen consumption rate (OCR) of complex I-driven respiration using NADH and Complex IV-driven respiration using ascorbate in male isolated liver mitochondria. (E) OCR of Complex II-driven respiration using succinate in the presence of rotenone and Complex IV-driven respiration using ascorbate in male isolated liver mitochondria. (F) OCR normalized to CS activity in male isolated liver mitochondria. (G) CS enzyme activity in female isolated liver mitochondria. (H) OCR of complex I-driven respiration using NADH and Complex IV-driven respiration using ascorbate in female isolated liver mitochondria. (I) OCR of Complex II-driven respiration using succinate in the presence of rotenone and Complex IV-driven respiration using ascorbate in female isolated liver mitochondria. (J) OCR normalized to CS activity in female isolated liver mitochondria. (C,G) n=7-8 mice/group; student’s t-test. (D-F, H-J) statistics for the overall effects of time or complex, diet, and the interaction represent the p value from a two-way ANOVA analysis; *p<0.05 from a Sidak’s post-test examining the effect of parameters identified as significant in the two-way ANOVA. Data represented as mean ± SEM.

To better understand how Val-R affects mitochondrial function, we isolated liver mitochondria. While there was no difference in citrate synthase in male liver (**Fig. 8C**), suggesting that there was no diet-induced difference in mitochondrial abundance, when we assessed CI, CII and CIV-related mitochondrial respiration we observed a significant diet effect when using either NADH or succinate/rotenone as substrates (**Figs. 8D-E**). When we quantified the oxygen consumption rate (OCR) normalized to mitochondrial abundance, we found a significant effect of diet, with a significant increase in CII and trending increase in succinate/rotenone-derived CIV (p=0.0683)-mediated respiration in Val-R-fed males (**Fig. 8F**). We also assessed the expression of OXPHOS protein abundance via western blot and observed no effect of diet in either sex (**Extended Fig. 7**), suggesting the Val-R induced changes in male mitochondrial complex activity are not driven by changes in complex abundance.

## Discussion

Dietary protein has emerged as a key regulator of longevity and health in both humans and rodents ^6, 11, 12, 15, 16, 17, 26, 44, 95^. Restriction of protein, BCAAs or isoleucine alone is sufficient to promote healthy aging and extend lifespan in mice, while blood levels of BCAAs are associated with diabetes, insulin resistance, and mortality in humans ^22, 24, 25, 26, 74, 96, 97^. Valine has recently been linked to adverse metabolic effects as well as cancer and inflammation ^22, 31, 32, 33, 34, 35, 98^, and here we explored the hypothesis that valine restriction (Val-R) would improve healthspan and extend longevity. We found that lifelong Val-R blunted weight gain and reduced adiposity, improved glycemic control and energy balance in both sexes, extended lifespan in males, and reduced frailty in both sexes.

Val-R produces some of the same effects as other amino acid restricted diets, but there are also effects unique to Val-R. Like both isoleucine restriction and methionine restriction, Val-R extends male lifespan, but unlike the others, Val-R fails to extend female lifespan ^19, 20, 24, 26, 99, 100, 101, 102, 103, 104^. All three interventions improve glucose tolerance, reduce frailty, protect against liver pathologies and cancer, and increase energy expenditure, partly through the induction of thermogenesis ^20, 22, 26, 105, 106^. Val-R uniquely fails to improve rotarod performance ^20, 26^, and induces distinct effects on cognition and neuroinflammation that have not been fully characterized in other dietary interventions.

At the molecular level, Val-R is distinct from isoleucine and methionine restriction. While isoleucine, methionine and short-term Val-R all increase circulating FGF21, long-term Val-R does not ^22, 26, 30, 105, 106^. Unlike isoleucine restriction and methionine restriction, which suppresses mTORC1, Val-R increases hepatic mTORC1 activity while leaving mTORC1 in the muscle unchanged ^23, 105, 107^. Moreover, Val-R decreases hepatic autophagy, an unexpected finding given the positive links between autophagy and longevity ^108, 109, 110, 111^. Differences in amino acid catabolism may underlie some of these effects: methionine catabolism yields the methyl donor SAM, while isoleucine catabolism generates both acetyl-CoA and succinyl-CoA, and valine catabolism yields only succinyl-CoA. As shown here, Val-R may also activate hepatic mTORC1 via increased PI3K-AKT signaling.

The response to Val-R was strongly sex-specific. While we observed an overall increase in genes within the BCAA degradation pathway in all three tissues, females displayed a greater increase, particularly in the liver. One possible explanation for this difference is that due to inante difference in BCAA catabolism, Val-R males may be effectively more valine restricted, with Val-R females retaining greater valine availability, possibly explaining the extension of male but not female lifespan. Differences in BCAA catabolism between the sexes has been observed in previous studies ^26, 112, 113, 114^. Our transcriptional analysis found that “Valine, leucine, and isoleucine degradation” were significantly upregulated in the livers of Val-R-fed females, but not in Val-R-fed males. Although frailty is closely correlated with longevity in both mice and humans ^115, 116, 117^, Val-R reduced frailty in females without extending their lifespan. We previously observed a similar uncoupling in isoleucine-restricted UM-HET3 females ^26, 63^. Perhaps while 67% restriction of dietary valine is sufficient to improve multiple health metrics in female mice, it is insufficient to extend female lifespan; if so, future studies could explore if a greater degree of restriction will extend female lifespan.

When we specifically investigated genes related to BCAA degradation, we found that the strongest changes in these genes were in the liver, with smaller changes in muscle and BAT. Notably, Val-R upregulated numerous genes that play a role in lipid and fatty acid metabolism, supporting the finding of other groups that valine alters fatty acid metabolism to impact fatty liver disease ^34, 118, 119, 120^. Further, we see a diet effect in reducing lipid droplet size and count in our male and female Val-R-fed mice. We did not expect this, as muscle is canonically the primary site of BCAA catabolism, and is traditionally thought of as very sexually dimorphic; and at the molecular level, liver, muscle and adipose have similar levels of sexually dimorphic gene expression ^121^. Understanding sex-specific differences in BCAA and AA metabolism may be important to understanding the basis for the beneficial and sex-specific impacts of Val-R and other diets on healthy aging.

Despite weight-normalized preservation of muscle mass, Val-R did not consistently improve functional performance. Inverted cling tests suggested modest benefits, but these were largely attributable to reduced body weight. We also saw benefits of Val-R on frailty, particularly with the physical/musculoskeletal scoring. We did not measure muscle strength, quality, or fiber type, which will be critical to examine in future studies examining how valine impacts muscle.

WGCNA analysis implicated PI3K-AKT, AGE-RAGE and MAPK signaling across liver, muscle and adipose tissue, with changes in males consistent with improved health and longevity ^122, 123, 124, 125, 126, 127, 128, 129, 130, 131^. Cellular senescence and related pathways (i.e. p53 signaling and NF Kappa B signaling) as well as type 2 diabetes and insulin secretion were also altered, particularly in the liver, further supporting the role of valine on health and aging; and these findings are supported by our recent finding that BCAA restriction inhibits hepatic senescence ^132^. Here, we found that long-term Val-R reduces senescent cell burden, not only in the liver, but across multiple tissues, demonstrating that Val-R protects from one of the hallmarks of aging.

When we looked at the genes in the PI3k-AKT signaling and MAPK signaling pathways in males, we observed an overall downregulation of PI3K-AKT signaling pathway genes by Val-R, with no overall effect on MAPK signaling genes. In females, we found an overall weaker correlation of modules to phenotypic traits, and in the blue module where the strongest associations were found, we observed only a partial enrichment for PI3K-AKT and MAPK signaling. There was no observable pattern in significant genes altered in the PI3K-AKT signaling pathway in Val-R-fed females. In contrast to this effect on gene expression, we found that AKT signaling was upregulated at the protein level, with increased phosphorylation of AKT S473 and FOXOs. This effect may be due to increased insulin sensitivity, but as decreased – not increased – PI3K-AKT signaling is associated with lifespan ^123, 133^, it seems unlikely this effect drives increased longevity.

Analysis of our transcriptomics data implicated liver mitochondria as a potential “hub” driving multi-tissue changes in gene expression in response to Val-R. Therefore, we isolated liver mitochondria and assessed their oxygen consumption rate. We found a male-specific increase in CII- and a non-significant increase in CIV (p=0.0683)-related respiration, with no changes in OCR in females, correlating with the male-specific lifespan extension. Increased CII activity promotes lifespan in *C. elegans* ^134^, CII activity is increased by rapamycin in flies ^135^, and perturbations of mitochondrial activity more generally have been linked to lifespan extension via stress response pathways, e.g. via mitohormesis ^136, 137^. How valine restriction alters mitochondrial function and whether increased CII activity is required for the increased lifespan of Val-R-fed males remains to be determined. Additionally, determining the effects of Val-R on mitochondrial function in other tissues, such as the brain, may also provide insight into the mechanisms by which Val-R improves other aspects of health, like neuroinflammation, in a sex-specific manner.

There are several limitations of this work. First, we examined only a single level of restriction. Studies of CR, PR, and geroprotective drugs like rapamycin have shown that different levels of restriction or drugs can yield different responses ^15, 95, 138, 139, 140^. Different levels of Val-R may be able to extend female lifespan and healthspan, and examining the graded response to Val-R may provide new insights into biological mechanisms as well as the basis for the sex-specific effects we observed. Next, we analyzed longevity in only a single inbred strain of mice, C57BL/6J, and commenced our study when the mice were only 4 weeks old. There were advantages to this approach – in particular, the strain and age of onset were chosen to match conditions under which PR and BCAA restriction were previously shown to extend male lifespan by over 30% ^17^. However, it is possible that some of the benefits we see here may result from early-life effects; future studies should examine if Val-R initiated in mid- or late-life similarly promotes healthy aging. Different strains of mice often have distinct responses to lifespan interventions ^138, 141^, and interventions that begin later in life are more likely to be translatable than interventions begun in extremely young animals. Future studies of Val-R should be conducted in other strains of mice – for instance, genetically heterogeneous UM-HET3 mice which better mimic the genetic variability found in the human population – and begin Val-R in fully adult mice.

Our omics analysis was limited to transcriptional profiling, and further unbiased molecular analysis techniques such as metabolomics in multiple tissues would likely provide additional insights into the metabolic processes and mechanisms that drive aging that are modified by Val-R. Furthermore, although Val-R had promising effects on neuroinflammation, the only cognitive assay we performed was NOR. In future studies, it will be important to perform a more complete array of behavioral phenotyping assays, including those such as the Barnes maze or Morris water maze that provide information about spatial memory and hippocampal cognition, that may be more directly impacted by the effects of Val-R on hippocampal neuroinflammation.

Lastly, the diets we used here are extremely well-controlled, and are isocaloric, matched for the quantity and source of carbohydrates and fats, and isonitrogenous. However, choices still had to be made; the diets used here are based on the AA profile of a whey-based diet; and the reduction of valine was balanced by a small increase in a number of non-essential AAs. The protein-to-carbohydrate ratio, the type and levels of dietary carbohydrates and fats consumed, or the ratio of different amino acids to valine could all play a role in the responses we observed. The optimal level of protein may vary by age – it is generally believed that older people need to consume more protein – and varying levels of valine at different ages could lead to different results.

In conclusion, we have shown that dietary restriction of valine can promote healthy aging in both sexes, and extend the lifespan of male, but not female mice. Our results are consistent with a growing consensus showing that higher levels of valine are associated with negative impacts on healthy aging, including insulin resistance, cardiovascular disease, and cancer. Additional research will be required to identify the optimal level of dietary valine, especially across mice of different sexes, ages, and genetic background, and to identify if there are negative consequences to valine restriction. Our results highlight how protein quality, the specific amino acids that make up the protein, is as important in mediating healthy aging as total protein or the number of calories. While additional tests are needed to fully understand how valine affects health in humans, our results support the idea that lowering the amount of dietary valine may promote healthy aging.

## Data Availability Statement

RNA-seq data have been deposited with the Gene Expression Omnibus and are accessible through accession number GSE298741. The data that support the plots within this article and other findings of this study, including full scans of western blot images, are provided as Source Data files.

## Author Contributions

Experiments were performed in the Lamming laboratory, except for the analysis of the brain IHC, which was performed in the Sadagurski lab and the microCT analysis on the femur bone, which was performed in the Yakar lab. MC, MS, SY, WAR, ADA, and DWL conceived the experiments and secured funding. MFC, IA, PL, SML, LEB, RAM, HSMJ, DNHM, SY, TL, CLG, RB, MMS, YL, IG, CYY, YL, and BK performed the experiments. MC, DNHM, HSMJ, RAM, CLG, TL, CIO, MPK, MS and DWL analyzed the data. MFC, CLG, MPK, MS, and DWL wrote and edited the manuscript.

