## Extended Figures for "Lifelong restriction of dietary valine has sex-specific benefits for health and lifespan in mice"

### Extended Figure 1

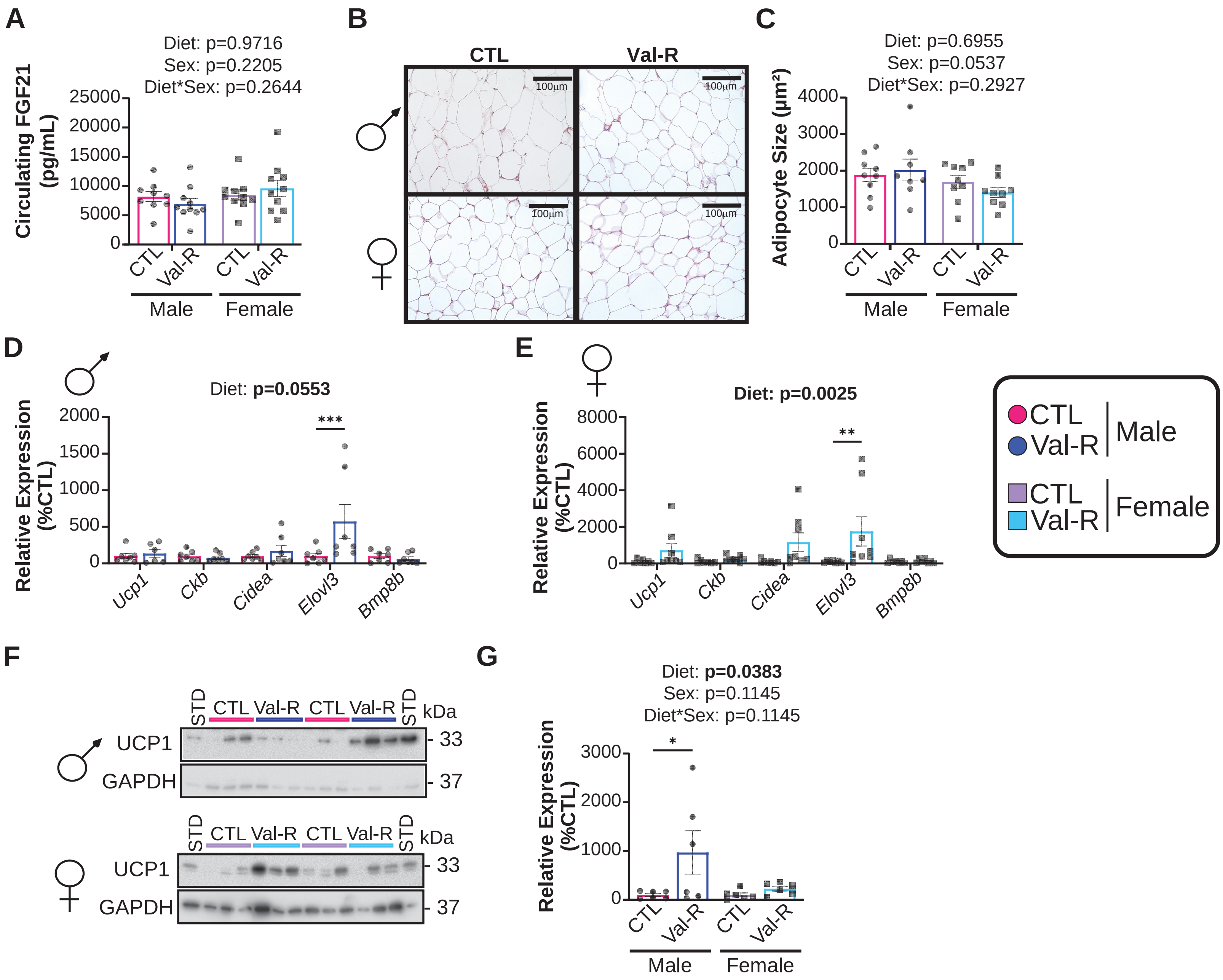

#### Extended Figure Legends

##### Extended Figure 1: FGF21-UCP1 axis in the iWAT of Val-R-fed mice.

(A) Circulating FGF21 at 19 months of age. (n=10-12 mice/group) (B) Hematoxylin and eosin (HE) staining (representative images; scale bar=100µm, 40X magnification) from iWAT of male and female mice. (C) Quantified adipocyte size ($\mu m^2$) from HE-stained iWAT images from male and female mice (n=6-9 mice/group). (D-E) The mRNA expression of thermogenic genes was quantified in the iWAT of male (D) and female (E) mice (n=7-8 mice/group). (F-G) Western blots of UCP1 protein normalized to GAPDH in male and female iWAT (F) and quantification (G), analysis via two-way ANOVA. (A, C-E, G) statistics for the overall effects of sex, diet, and the interaction represent the p value from a two-way ANOVA, *p<0.05, **p<0.01, ***p<0.001 from a Sidak's post-test examining the effect of parameters identified as significant in the two-way ANOVA. Data represented as mean ± SEM.

### Extended Figure 2

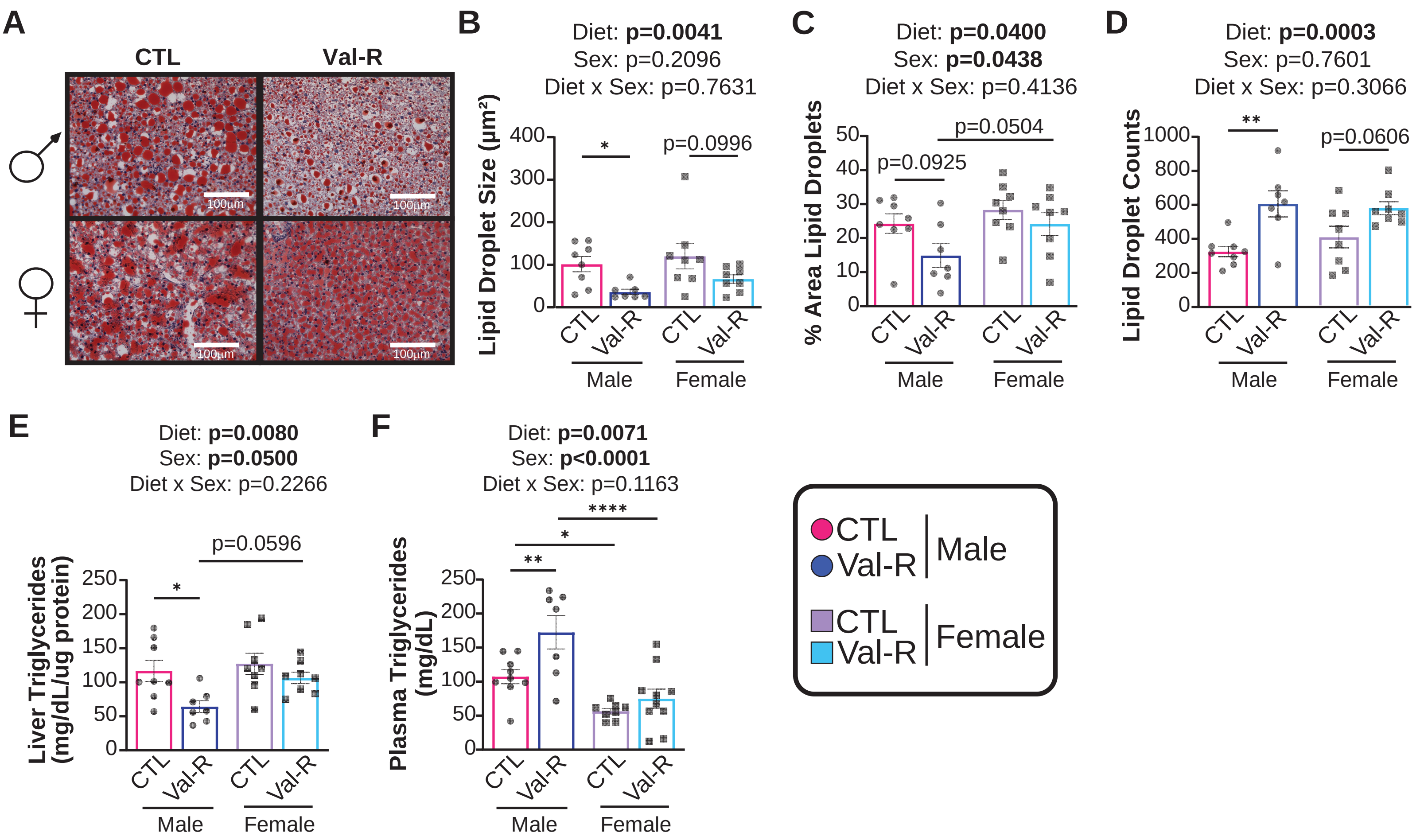

**Extended Figure 2: Val-R protects from hepatic steatosis**

(A) Oil-Red-O (ORO) staining (representative images; scale bar=100µm, 40X magnification) from the liver of male and female mice (n=7-8 mice/group). (B-D) Quantified lipid droplet size ($\mu m^2$) (B), percent of area of lipid droplets (C) and lipid droplet count ( D) calculated from ORO-stained liver images (n=7-8 mice/group). (E) Liver triglycerides normalized to protein at 24 months of age. (n=7-8 mice/group) (F) Circulating triglycerides at 24 months of age. (n=7-9 mice/group) (B-F) statistics for the overall effects of sex, diet, and the interaction represent the p value from a two-way ANOVA. *$p<0.05$, **$p<0.01$, ****$p<0.0001$ from Sidak's post-test examining the effect of parameters identified as significant in the two-way ANOVA. Data represented as mean ± SEM.

### Extended Figure 3

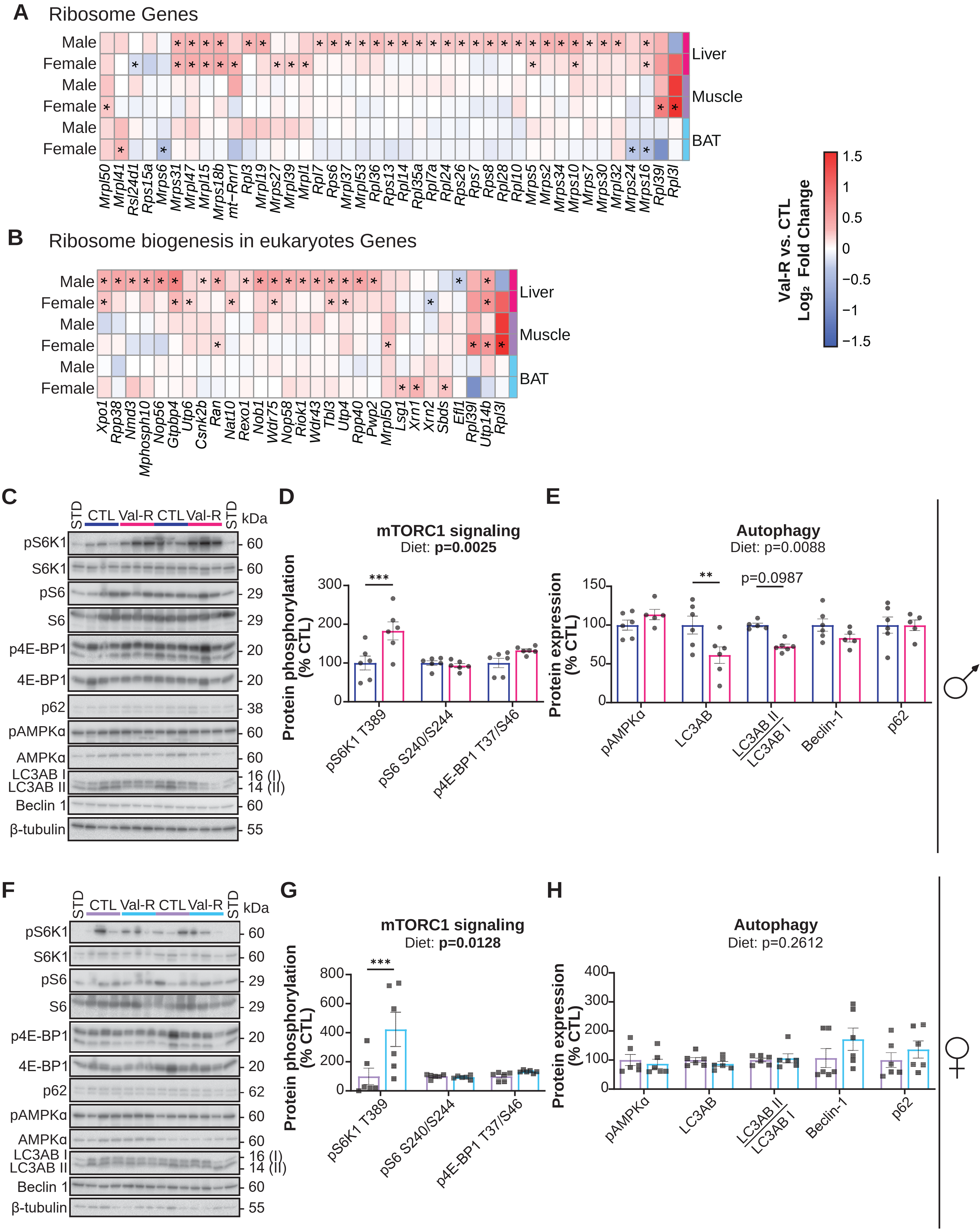

**Extended Figure 3: Val-R induced hepatic mTORC1 signaling**

(A-B) Altered genes in the “Ribosome” (A) and “Ribosome biogenesis in eukaryotes” (B) pathways in the liver, muscle and BAT. n=6-10 mice/group. (C) Western blots of the analyzed proteins in male livers. (D) Phosphorylation of S6K1 T389, S6 S240/S244 and 4E-BP1 T37/S46, normalized to the expression of the respective protein. (E) Phosphorylation of AMPKα normalized to the expression of AMPKα and protein expression of LC3AB, Beclin-1 and p62 normalized to expression of β-tubulin (n=5-6 mice/group). (F) Western blots of the analyzed proteins in female livers. (G) Phosphorylation of S6K1 T389, S6 S240/S244 and 4E-BP1 T37/S46 normalized to their total protein. (H) Phosphorylation of AMPKα normalized to the expression of AMPKα and protein expression of LC3AB, Beclin-1 and p62 normalized to expression of β-tubulin (n=6 mice/group). (D-E, G-H) statistics for the overall effects of gene or sex, diet, and the interaction represent the p value from a two-way ANOVA analysis; $^{**}p<0.01$, $^{***}p<0.001$ from a Sidak’s post-test examining the effect of parameters identified as significant in the two-way ANOVA. Data represented as mean ± SEM.

### Extended Figure 4

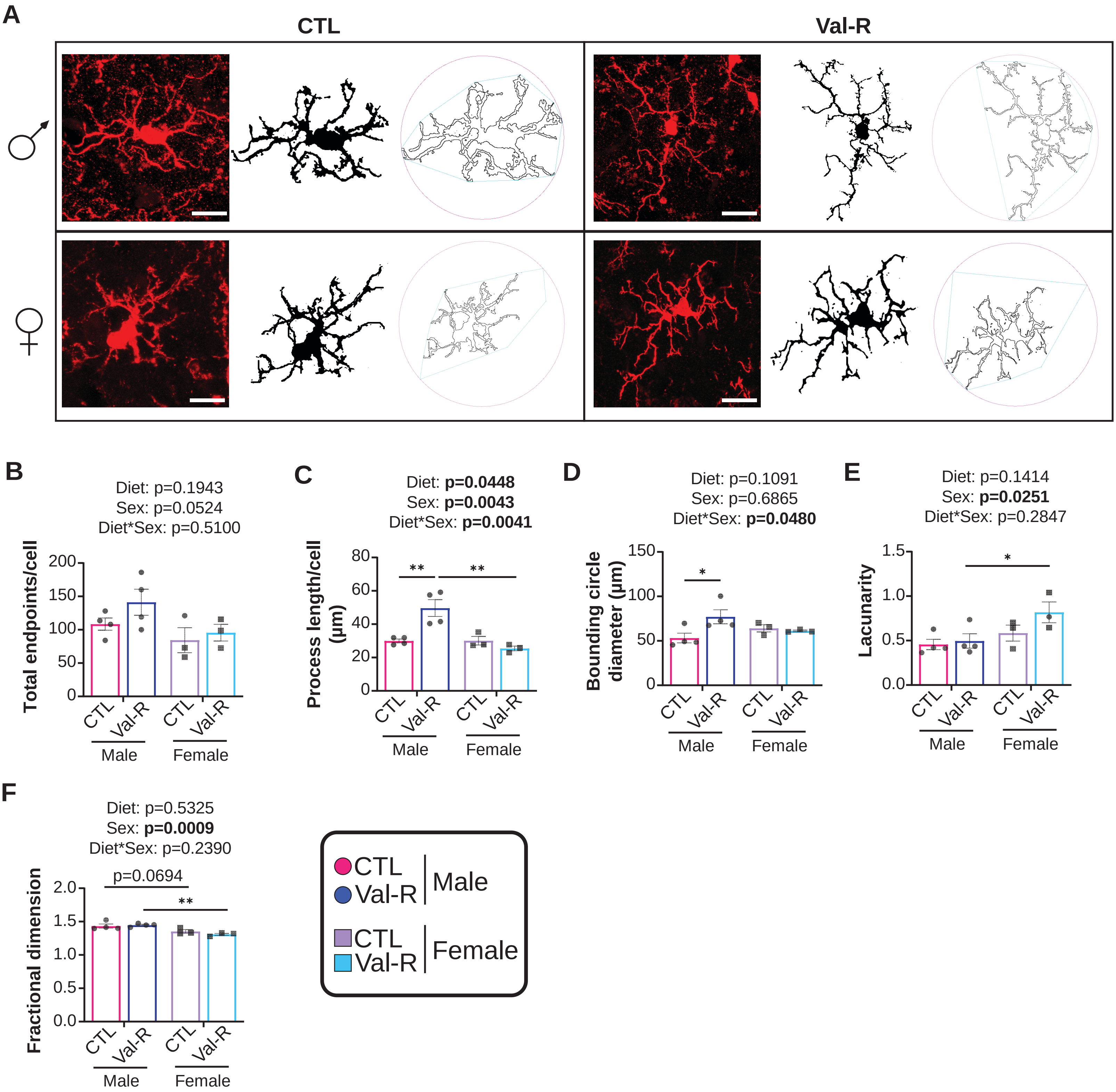

**Extended Figure 4: Val-R reduces activation of microglia in the hippocampus.**

(A) Representative images of immunofluorescent microglia with their stacked and reconstructed 3D images. Scale bar represents 10 µm. (B-F) Quantified total endpoints per cell (B), process length (C), bounding circle diameter (D), lacunarity (E) and fractional dimension (F). n=3-4 mice/group. Statistics for the overall effects of sex, diet, and the interaction represent the p value from a two-way ANOVA analysis; *$p<0.05$, **$p<0.01$, ****$p<0.0001$ from a Sidak's post-test examining the effect of parameters identified as significant in the two-way ANOVA. Data represented as mean ± SEM.

### Extended Figure 5

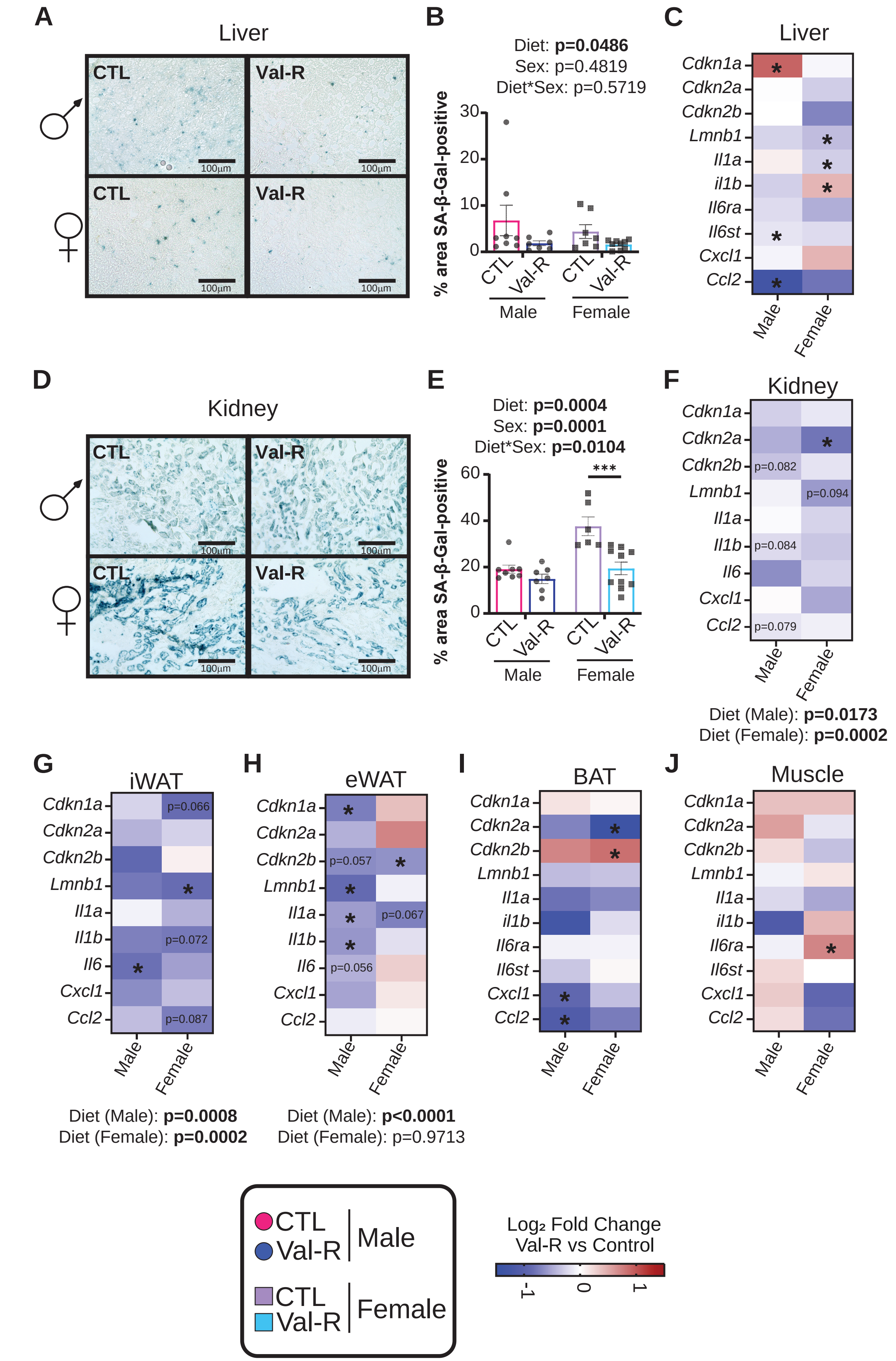

**Extended Figure 5: Val-R reduces senescence in multiple tissues.**

(A-C) Hepatic SA-β-Gal staining at 40X magnification (scale bar = 100 µm) (A) with quantification of SA-βGal-positive cells (B) and $\log_2$ fold-change from RNA sequencing analysis of pre-selected senescence and SASP related genes (C). (D-F) Kidney SA-β-Gal staining at 40X magnification (scale bar = 100 µm) (D) with quantification of SA-βGal-positive cells (E) and $\log_2$ fold-change of mRNA expression of senescence and SASP genes of kidney (F). (G-H) $\text{Log}_2$ fold-change of mRNA expression of senescence genes in the iWAT (G) and eWAT (H). (I-J) $\text{Log}_2$ fold-change from RNA sequencing analysis of pre-selected senescence and SASP related genes in the BAT (I) and muscle(J). (C-J) n=6-10 mice/group. (B, E, F-H) statistics for the overall effects of sex, diet, and the interaction represent the p value from a two-way ANOVA analysis; *$p<0.05$, ***$p<0.001$ from a Sidak's post-test examining the effect of parameters identified as significant in the two-way ANOVA. (F-H) *$p<0.05$, student's t-test. (C, I-J) *$p<0.05$, moderated t-test. Data represented as mean ± SEM.

### Exended Figure 6

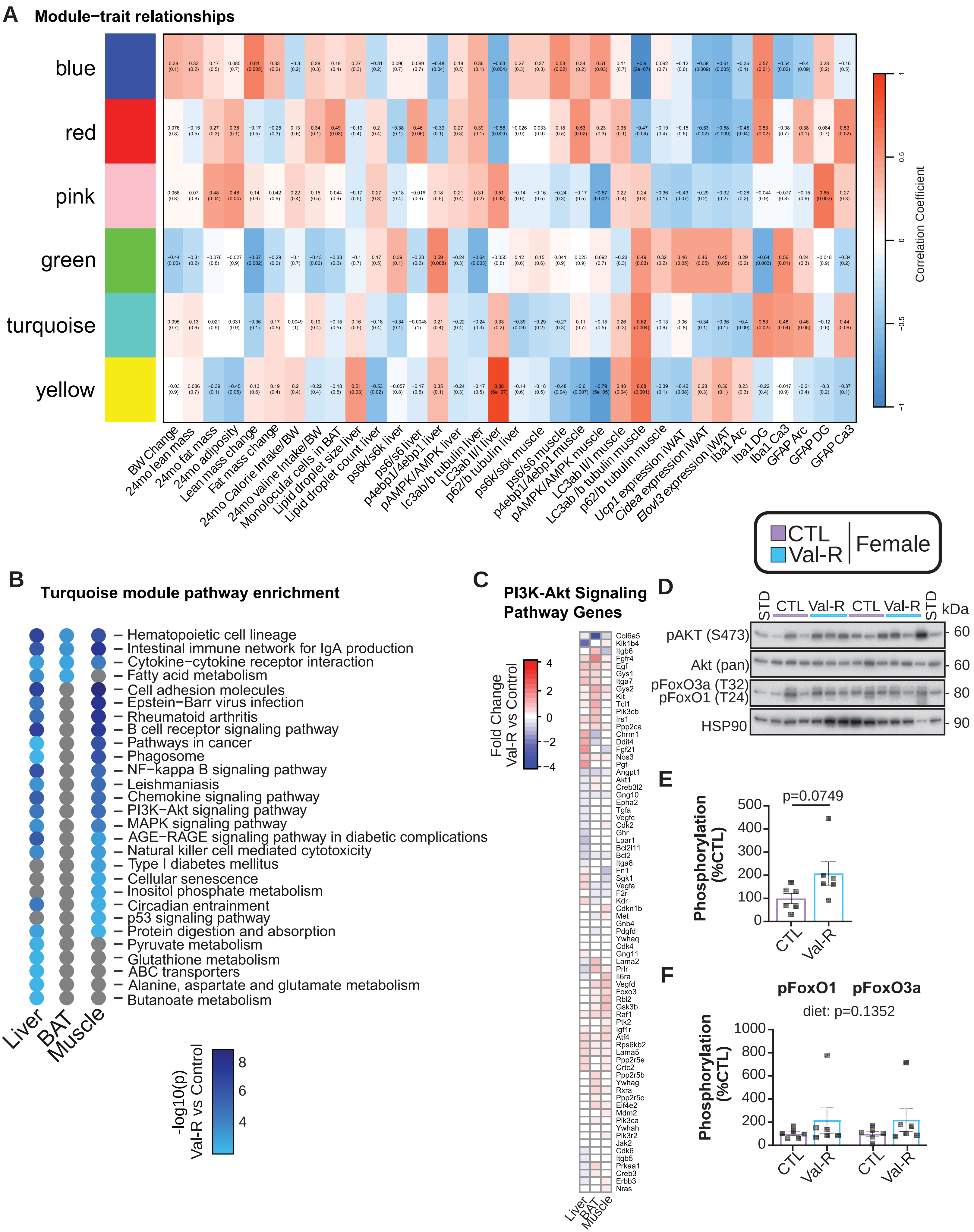

**Extended Figure 6: WGCNA analysis of selected modules and pathways in females.**

(A) Pearson correlation coefficient between the gene modules and selected phenotypic traits, numbers in brackets indicate the corresponding p values in females (n=6-10 mice/group). (B) Selected KEGG pathway enrichment of the blue module. Gray dots indicate no alterations in that pathway for that tissue. (C) Altered genes in the PI3K-Akt signaling pathway in the liver, muscle and BAT. Genes shown were significantly altered (Benjamini-Hochberg (BH) adjusted $p<0.05$) by Val-R in at least one tissue in males or females. (D) Western blots of the analyzed proteins in female livers. (E) Phosphorylation of AKT S473 normalized to the expression of AKT. (F) Phosphorylation of FoxO1 and FoxO3a normalized to HSP90. (E) n=6 mice/group; t-test. (F) Statistics for the overall effect of diet represent the p-value from a two-way ANOVA analysis. Data represented as mean ± SEM.

### Extended Figure 7

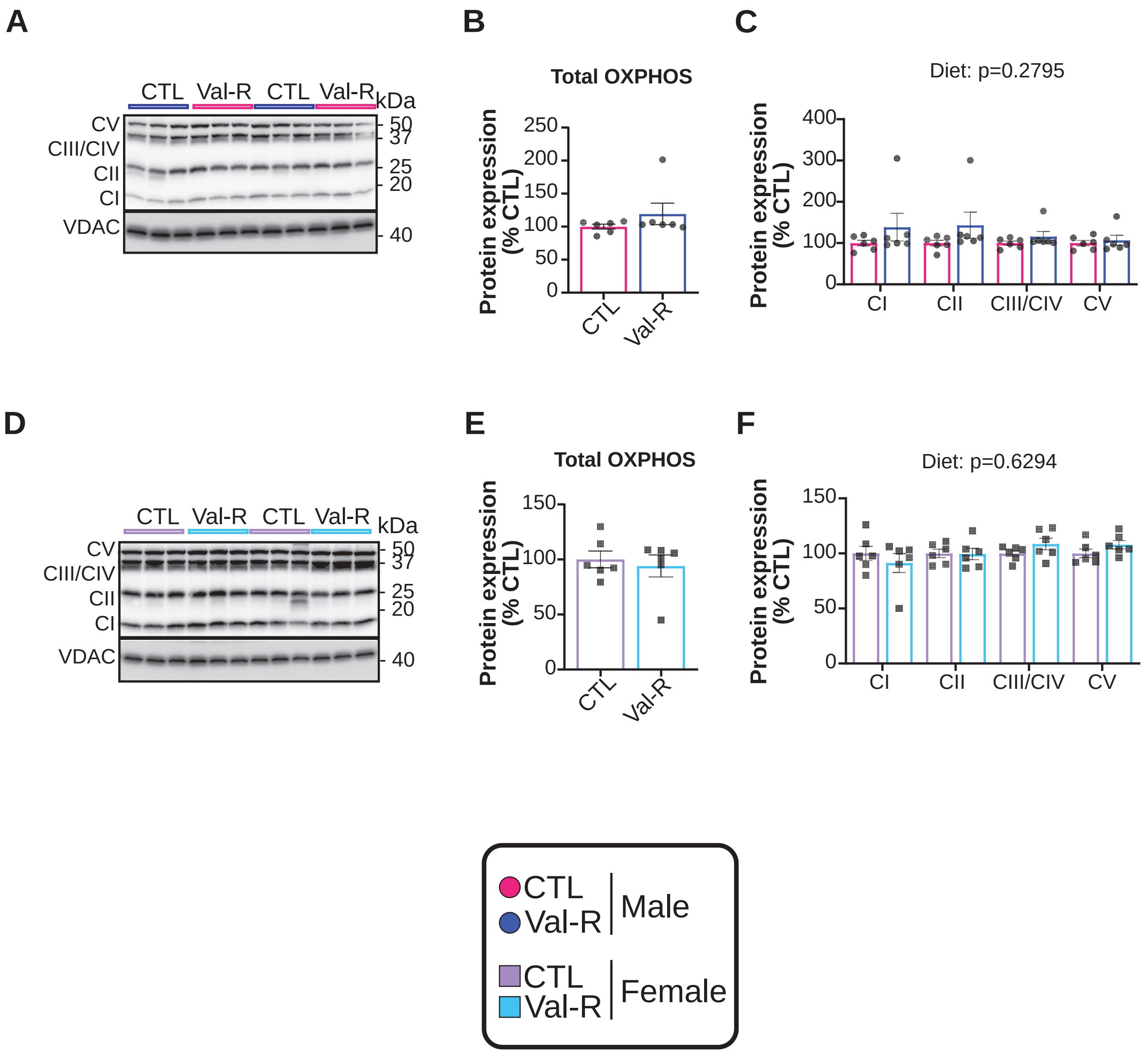

**Extended Figure 7: Val-R does not change OXPHOS protein abundance in male liver mitochondria.**

(A) Western blots of the analyzed proteins in male liver mitochondria. (B) Total OXPHOS normalized to VDAC. (C) Expression of CI, CII, CIII/CIV and CV normalized to VDAC. (D) Western blots of the analyzed proteins in female liver mitochondria. (E) Total OXPHOS normalized to VDAC. (F) Expression of CI, CII, CIII/CIV and CV normalized to VDAC. (B, E) n=6 mice/group; student's t-test. (C, F) Statistics for the overall effects of complex, diet, and the interaction represent the p value from a two-way ANOVA analysis; *$p<0.05$ from a Sidak's post-test examining the effect of parameters identified as significant in the two-way ANOVA. Data represented as mean ± SEM.
