## Supplemental Figures and Supplemental Table Legends for "Lifelong restriction of dietary valine has sex-specific benefits for health and lifespan in mice"

### Supplementary Figure 1

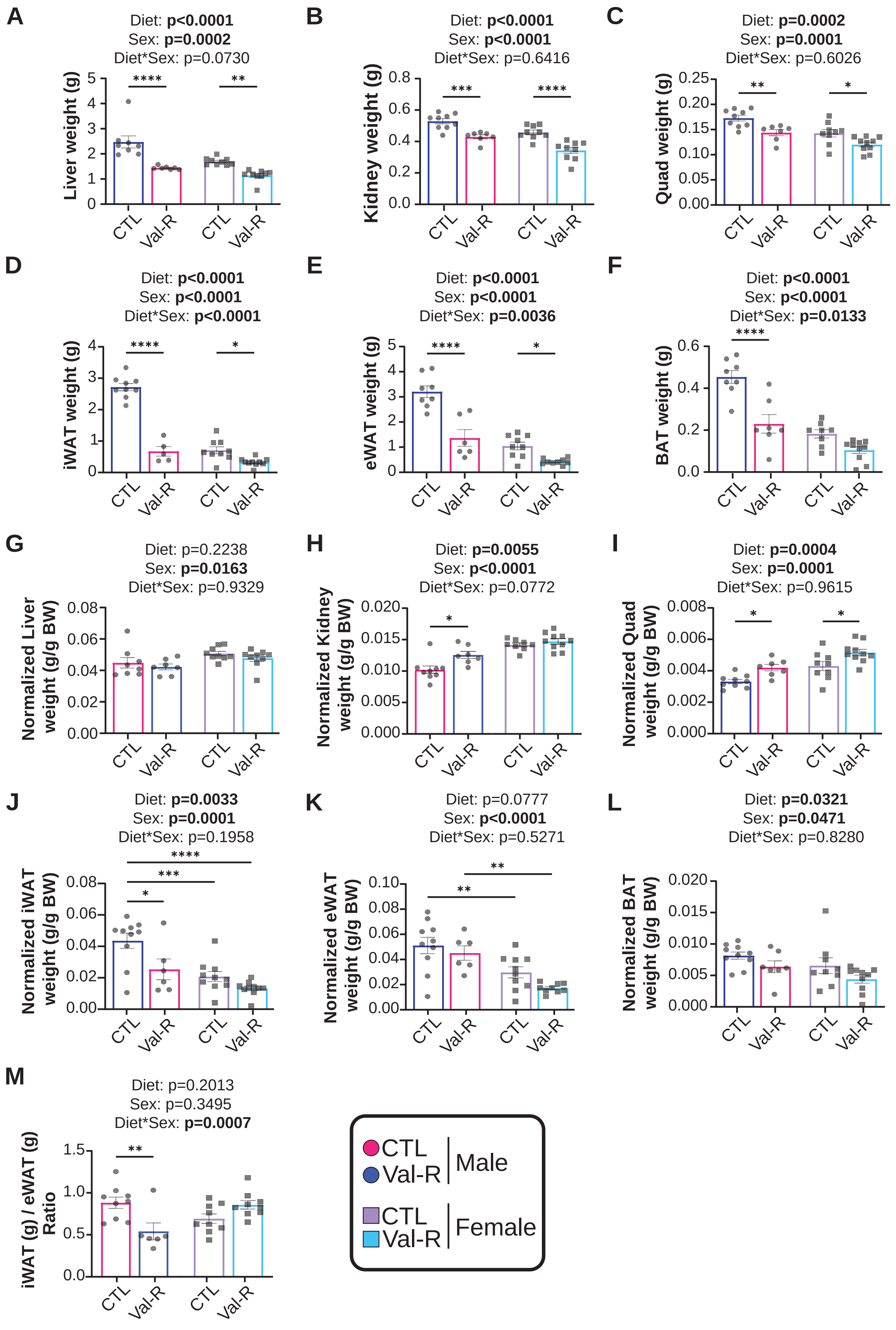

**Supplementary Figure Legends**

**Supplementary Figure 1: Val-R reduces adipose tissue.**

(A) Liver weight in grams. (B) Kidney weight in grams. (C) Quadricep muscle (quad) weight in grams. (D) Inguinal white adipose tissue (iWAT) weight in grams. (E) Epididymal white adipose tissue (eWAT) weight in grams. (F) Brown adipose tissue (BAT) weight in grams. (G) Liver weight in grams normalized to grams of body weight per mouse (g BW). (H) Kidney weight in grams normalized to g BW. (I) Quad weight in grams normalized to g BW. (J) iWAT weight in grams normalized to g BW. (K) eWAT weight in grams normalized to g BW. (L) BAT weight in grams normalized to g BW. (M) iWAT to eWAT ratio (A-M) n=6-10 mice/group; statistics for the overall effects of sex, diet, and the interaction represent the p value from a two-way ANOVA analysis; *p<0.05, **p<0.01, ***p<0.001, ****p<0.0001 from a Sidak's post-test examining the effect of parameters identified as significant in the two-way ANOVA. Data represented as mean ± SEM.

### Supplementary Figure 2

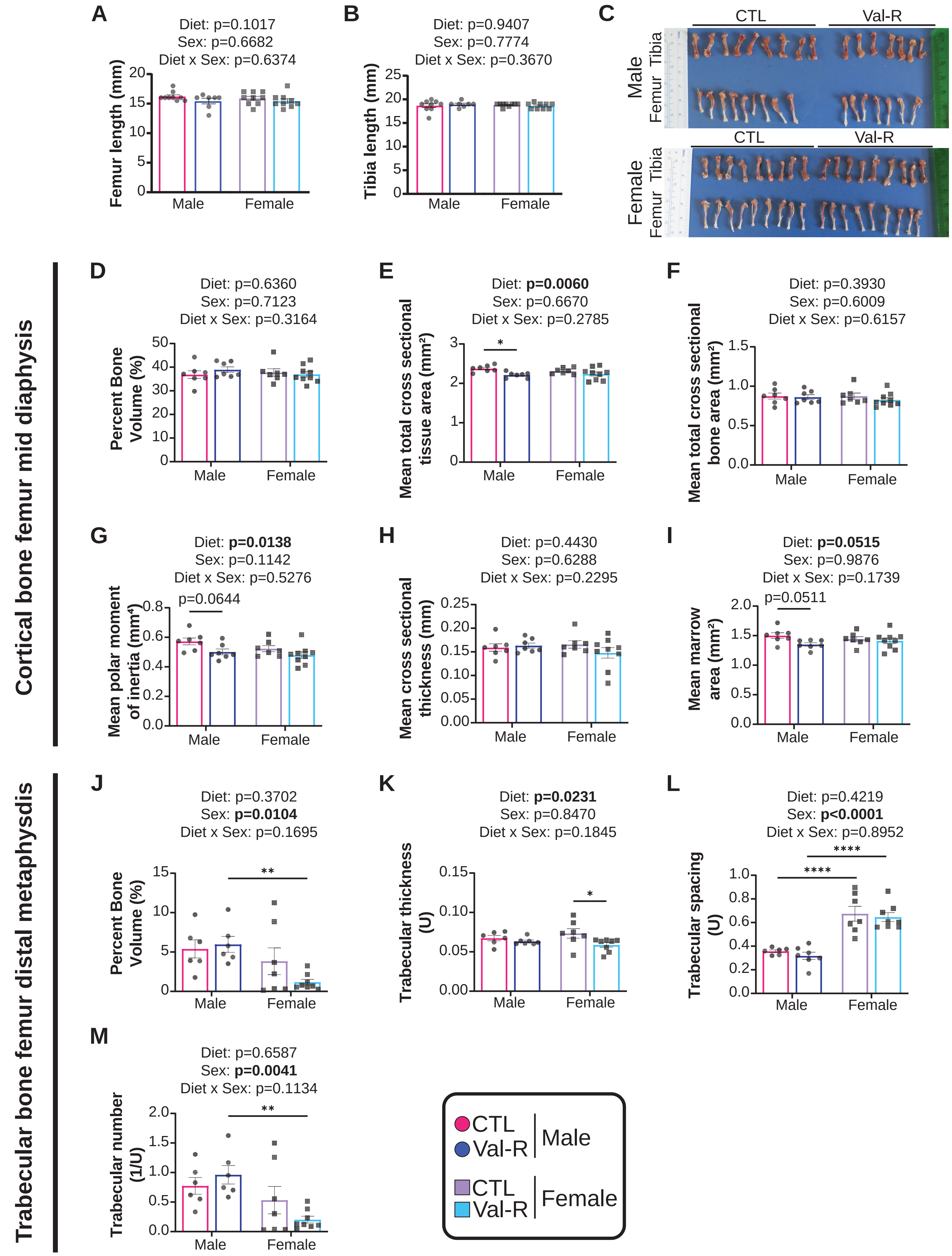

**Supplementary Figure 2: Val-R displays sex- and diet effects on femur bone morphology.**

(A) Femur bone length in millimeters. (B) Tibia bone length in millimeters. (C) Images of male and female tibia and femur length. (D-I) Mid diaphysis location of the cortical bone femur's percent bone volume (D), mean total cross-sectional tissue area in millimeters$^2$ (E), mean total cross-sectional bone area in millimeters$^2$ (F), mean polar moment in millimeters$^2$ (G), mean cross-sectional thickness in in millimeters (H), and mean marrow area in millimeters$^2$ (I). (J-M) Distal diaphysis location of the trabecular bone femur's percent bone volume (J), trabecular thickness in arbitrary units (K), trabecular spacing in arbitrary units (L), trabecular number in 1/arbitrary units (M). (A-B, D-M) n=6-10 mice/group; statistics for the overall effects of sex, diet, and the interaction represent the p value from a two-way ANOVA analysis; *$p<0.05$, **$p<0.01$, ****$p<0.0001$ from a Sidak's post-test examining the effect of parameters identified as significant in the two-way ANOVA. Data represented as mean ± SEM.

### Supplementary Figure 3

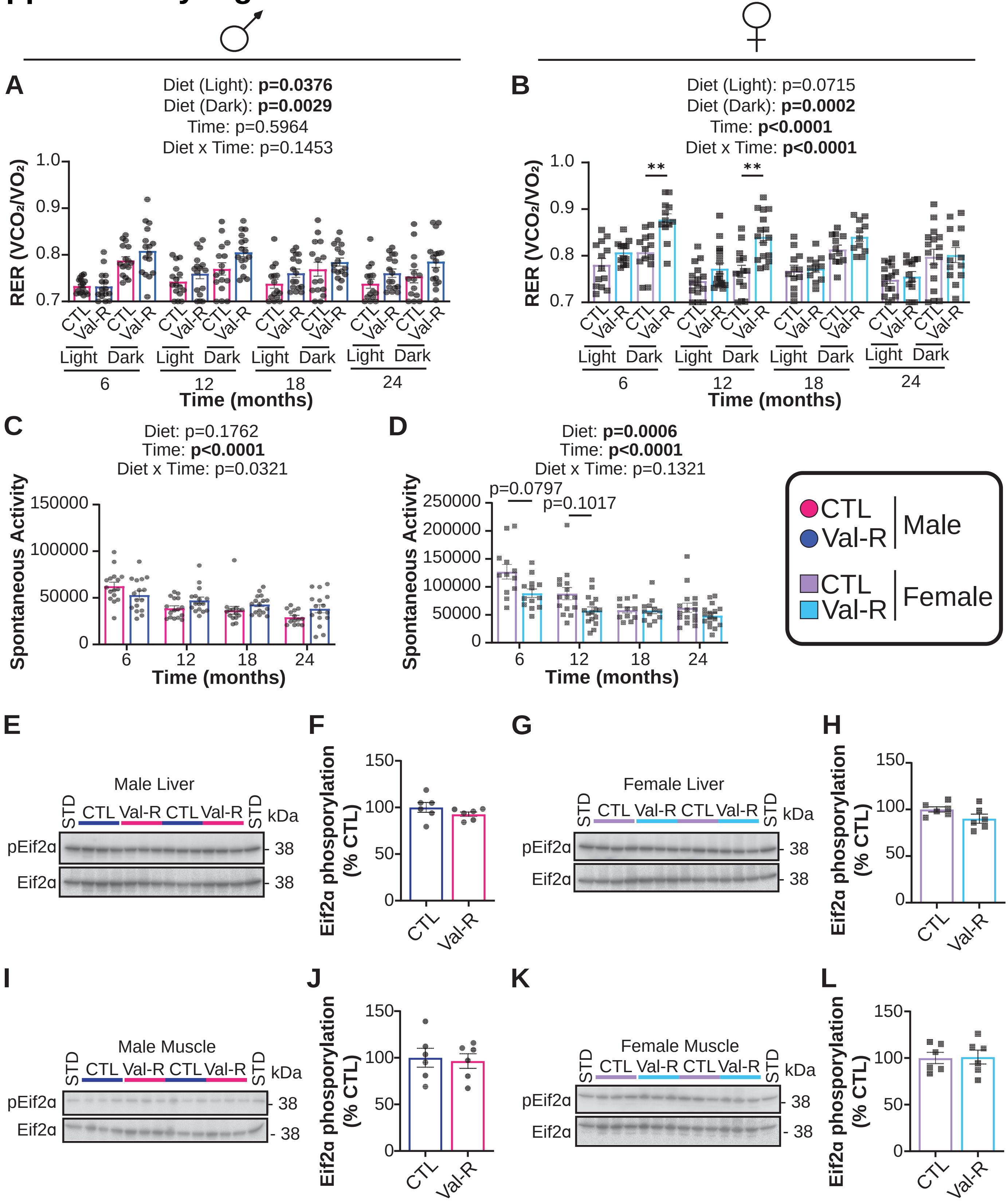

**Supplementary Figure 3: Val-R increases energy expenditure independently of RER, activity and Eif2α.**

(A-B) Respiratory exchange ratio ($VCO_2/VO_2$) in the light and dark phase at 6, 12, 18 and 24 months of age in male (A) and female (B) mice. (C-D) Spontaneous activity over a 24-hour period at 6, 12, 18 and 24 months of age in male (C) and female (D) mice. (E-F) Representative blots (E) and quantification (F) of phosphorylated Eif2α in male livers. (G-H) Representative blots (G) and quantification (H) of phosphorylated Eif2α in female livers. (I-J) Representative blots (I) and quantification (J) of phosphorylated Eif2α in male muscles. (K-L) Representative blots (K) and quantification (L) of phosphorylated Eif2α in female muscles. (A-D) n=15-16 mice/group; statistics for the overall effects of sex, diet (light and dark phase assess separately), and the interaction represent the p value from a two-way ANOVA analysis; *p<0.05, **p<0.01, ***p<0.001, ****p<0.0001 from a Sidak's post-test examining the effect of parameters identified as significant in the two-way ANOVA. (E-L) n=5-6 mice/group; student's t-test. Data represented as mean ± SEM.

### Supplementary Figure 4

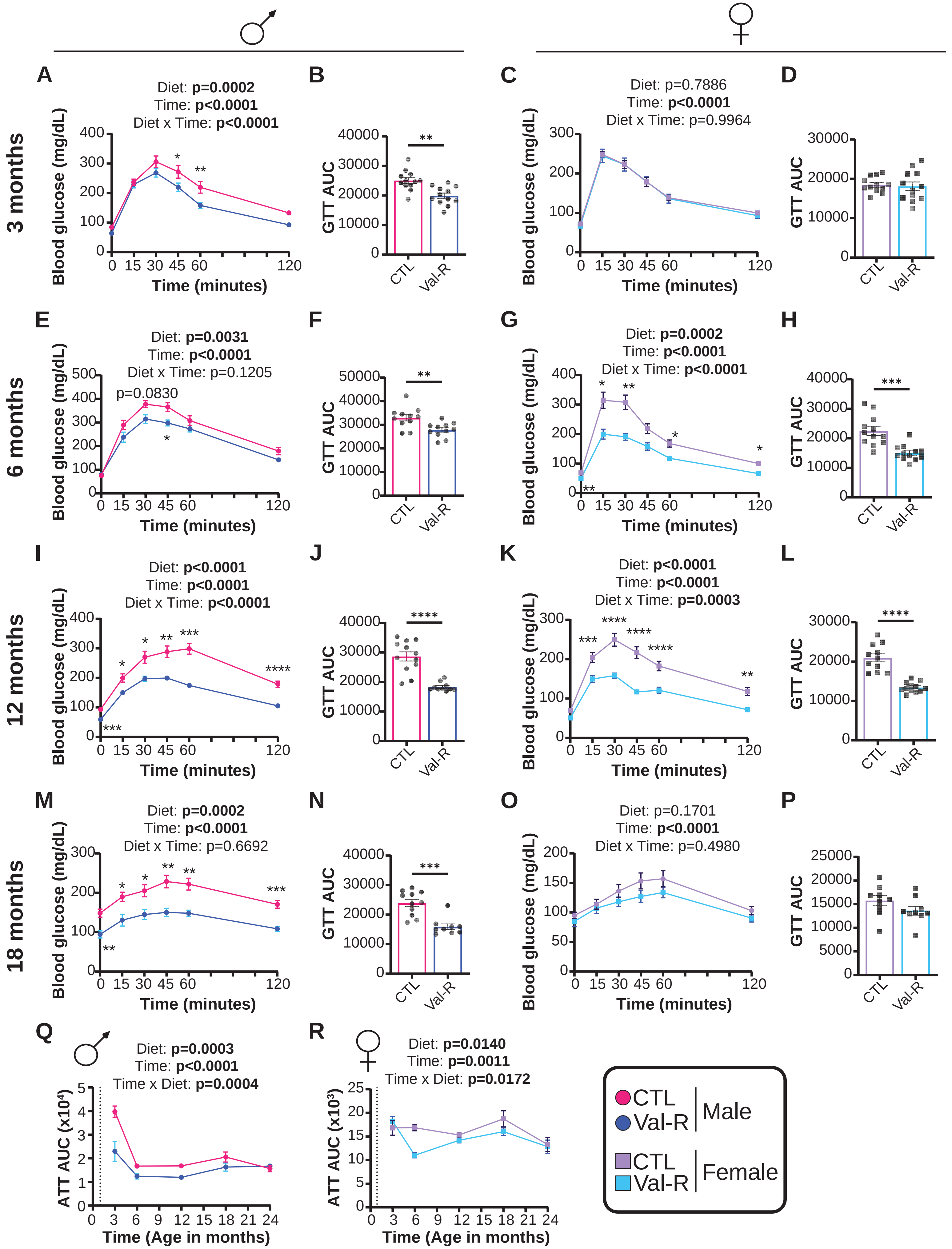

**Supplementary Figure 4: Val-R improves glucose tolerance at multiple ages.**

(A-P) Glucose tolerance test (GTT) and its associated area under the curve (AUC) in male and female mice conducted at 3, 6, 12 and 18 months of age. n=11-12 mice/group. (Q-R) Alanine tolerance test area under the curve (ATT AUC) over time in male (Q) and female (R) mice. (A, C, E, G, I, K, M, O, Q-R) statistics for the overall effects of time, diet, and the interaction represent the p value from a two-way ANOVA analysis; *p<0.05, **p<0.01, ***p<0.001, ****p<0.0001 from a Sidak's post-test examining the effect of parameters identified as significant in the two-way ANOVA. (B, D, F, H, J, L, N, P) student's t-test; **p<0.01, ***p<0.001, ****p<0.0001. Data represented as mean ± SEM.

### Supplementary Figure 5

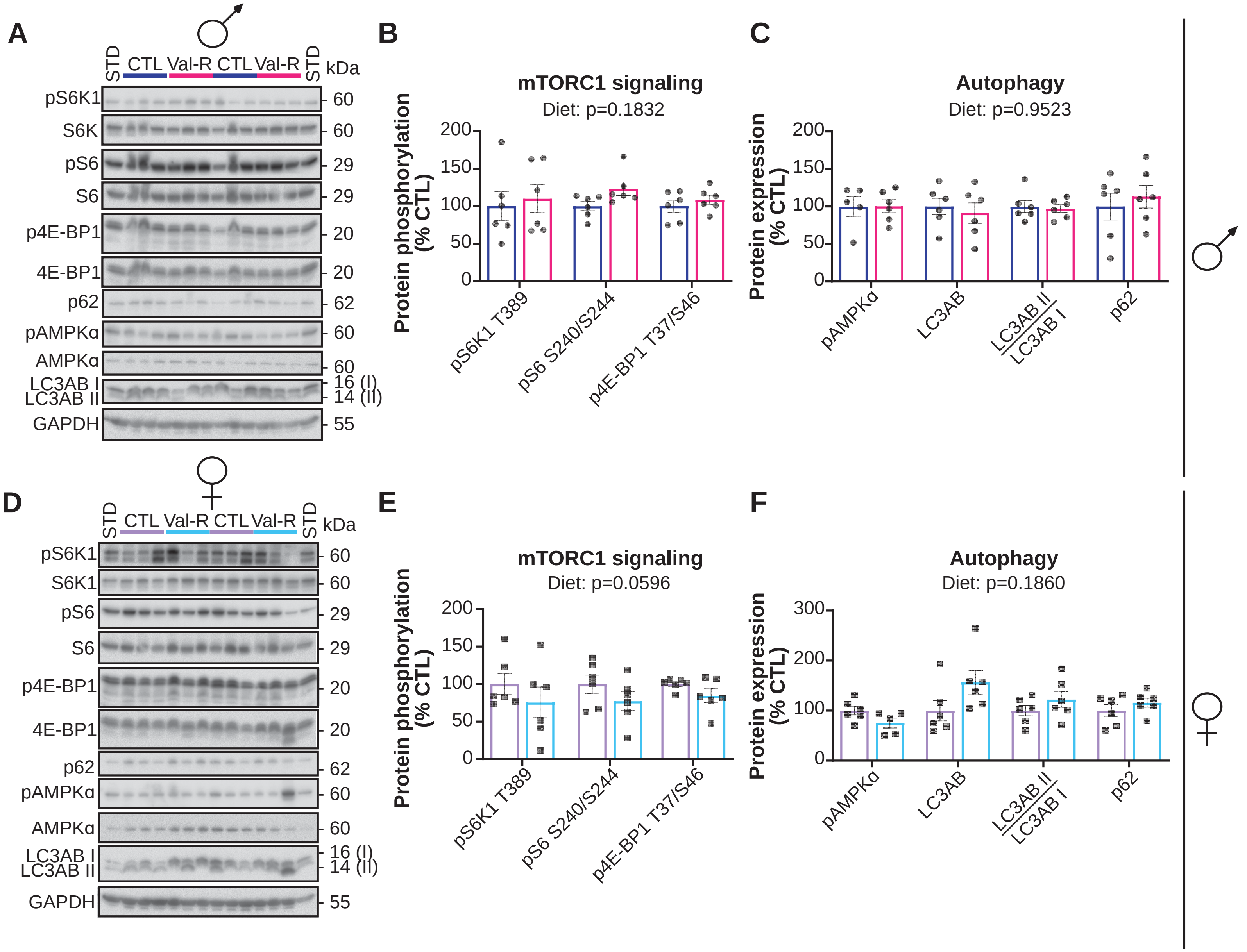

**Supplementary Figure 5: Val-R does not alter mTORC1 signaling in the muscle.**

(A) Western blots of the analyzed proteins in male quadricep muscle. (B) Phosphorylation of S6K1 T389, S6 S240/S244 and 4E-BP1 T37/S46 normalized to the expression of the respective protein. (C) Phosphorylation of AMPKα normalized to the expression of AMPKα and protein expression of LC3AB, Beclin-1 and p62 normalized to the expression of GAPDH (n=5-6 mice/group). (D) Western blots of the analyzed proteins in female muscles. (E) Phosphorylation of S6K1 T389, S6 S240/S244 and 4E-BP1 T37/S46 normalized to the expression of the respective protein. (F) Phosphorylation of AMPKα normalized to the expression of AMPKα and protein expression of LC3AB, Beclin-1 and p62 normalized to expression of GAPDH in female muscles (n=6 mice/group). (B-C, E-F) Statistics for the overall effects of gene or sex, diet, and the interaction represent the p value from a two-way ANOVA analysis; $^{**}p<0.01$, $^{****}p<0.0001$ from a Sidak's post-test examining the effect of parameters identified as significant in the two-way ANOVA. Data represented as mean ± SEM.

### Supplementary Figure 6

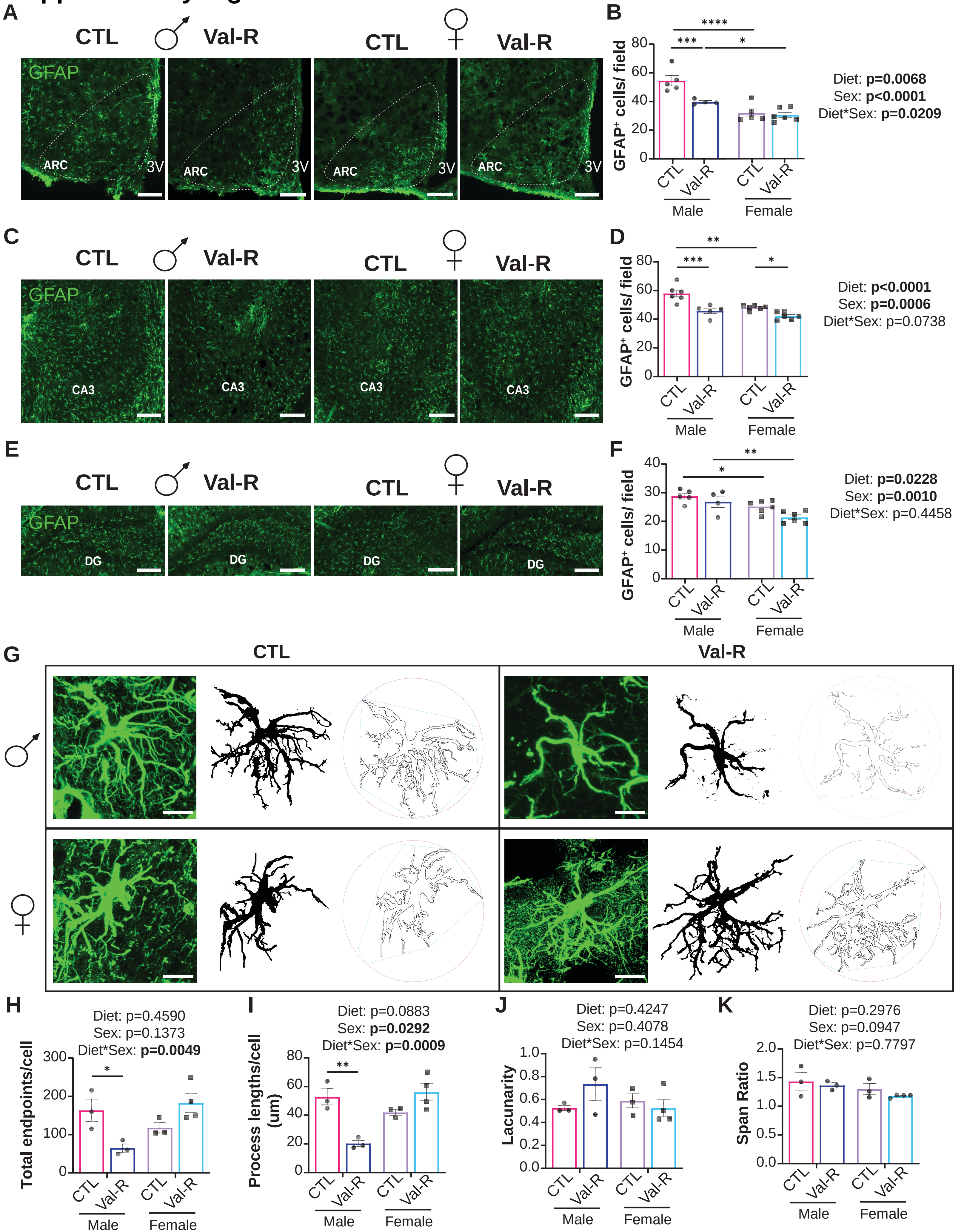

**Supplementary Figure 6: Val-R reduces GFAP and reactive astrocytes in the brain.**

(A, C, E) Representative images of GFAP staining in the Arc (A), CA3 (C) and DG (E) of the brain. Scale bar represents 200 µm. (B, D, F) Quantified staining of Iba1 in the Arc (B), CA3 (D) and DG (F). (G) Representative images of immunofluorescent astrocytes with their stacked and reconstructed 3D images. Scale bar represents 10 µm. (H-K) Quantified total endpoints per cell (H), process length (I), lacunarity (J) and span ratio (K). (B, D, F, H-K) n=3-6 mice/group. Statistics for the overall effects of sex, diet, and the interaction represent the p-value from a two-way ANOVA analysis; *p<0.05, **p<0.01, ***p<0.001, ****p<0.0001 from a Sidak's post-test examining the effect of parameters identified as significant in the two-way ANOVA. Data represented as mean ± SEM.

### Supplementary Figure 7

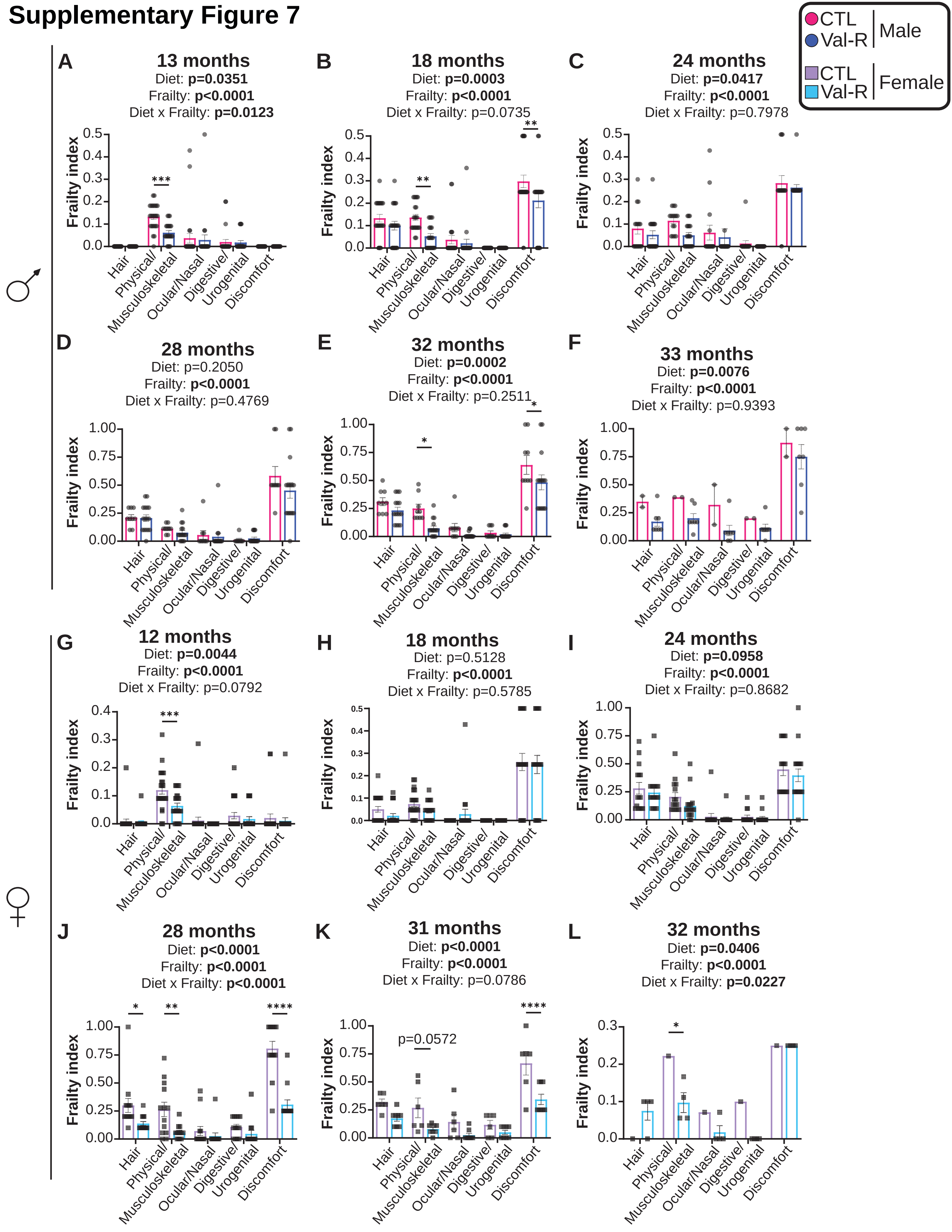

**Supplementary Figure 7: Val-R reduces physical/musculoskeletal and discomfort-related frailty scoring.**

(A-L) Frailty index scoring (0-1) separated into subsections hair, physical/musculoskeletal, ocular/nasal, digestive/urogenital and discomfort were scored for male (A-F) and female (G-L) mice at 12, 18, 24, 28, 31-32 and 32-33 months of age. n=20-25 mice/group; statistics for the overall effects of frailty, diet, and the interaction represent the p value from a two-way RM ANOVA analysis; *$p<0.05$, **$p<0.01$, ***$p<0.001$, ****$p<0.0001$ from a Sidak's post-test examining the effect of parameters identified as significant in the two-way ANOVA. Data represented as mean ± SEM.

### Supplementary Figure 8

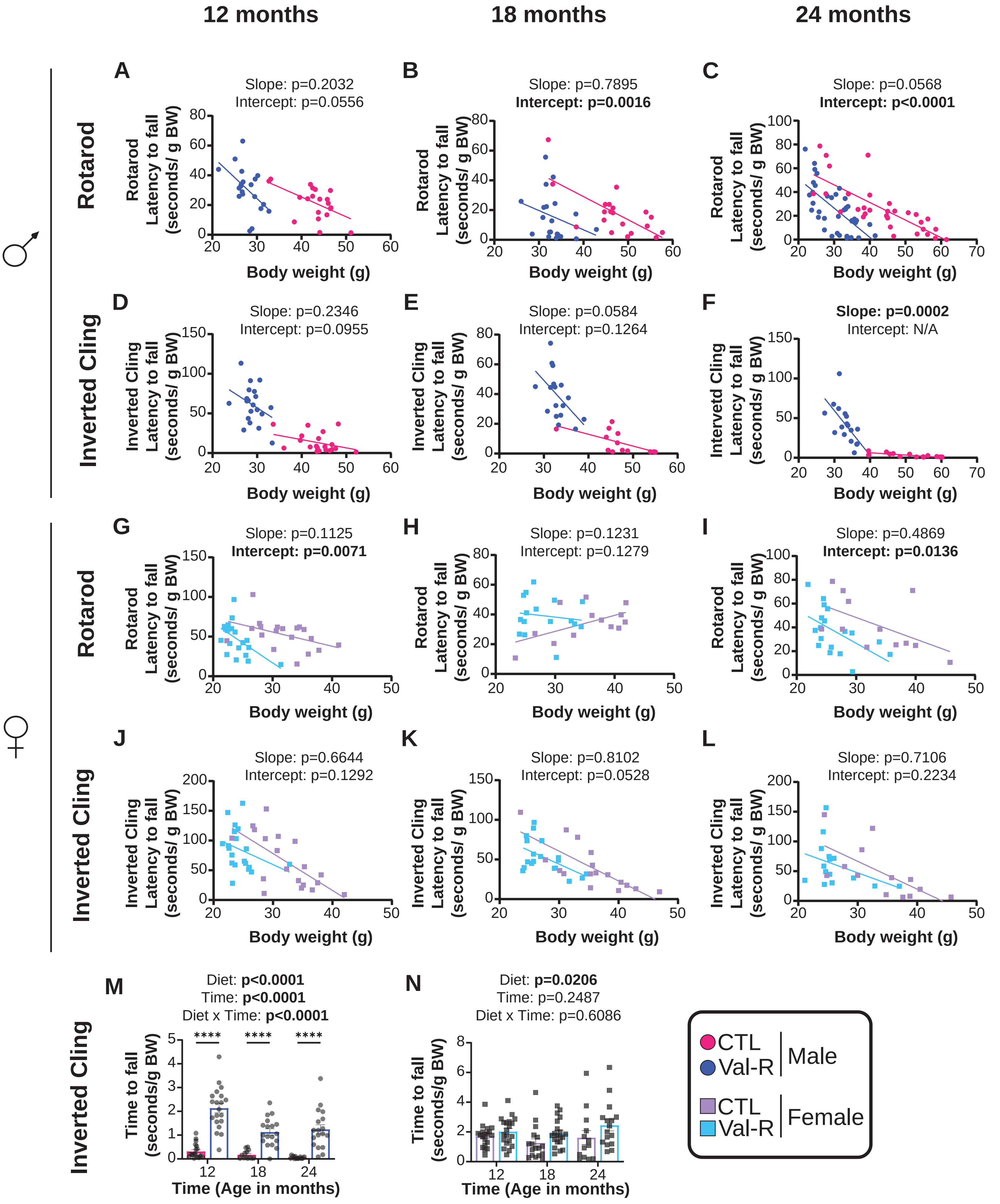

**Supplementary Figure 8: Val-R impacts rotarod and inverted cling performance.**

(A-C) Rotarod latency to fall in seconds of male mice as a function of body weight in (n=12-25 mice/group) at 12 (A), 18 (B), and 24 (C) months of age. (D-F) Inverted cling latency to fall in seconds of male mice as a function of body weight in (n=12-25 mice/group) at 12 (C), 18 (D), and 24 (E) months of age. (G-I) Rotarod latency to fall in seconds of female mice as a function of body weight in (n=12-25 mice/group) at 12 (G), 18 (H), and 24 (I) months of age. (J-L) Rotarod latency to fall in seconds of female mice as a function of body weight in (n=12-25 mice/group) at 12 (J), 18 (K), and 24 (L) months of age. (M-N) Inverted cling time to fall in seconds normalized to body weight in grams in (n=12-25 mice/group) in male (M) and female (N) mice. Statistics for the overall effects of time, diet, and interaction represent the p-value from a two-way ANOVA analysis; ****p<0.0001 from a Sidak's post-test examining the effect of parameters identified as significant in the two-way ANOVA. (A-L) data for each individual mouse is plotted; simple linear regression (ANCOVA) was calculated to determine if the slopes or elevations are equal; if the slopes are significantly different, differences in elevation cannot be determined. Data represented as mean ± SEM.

### Supplementary Figure 9

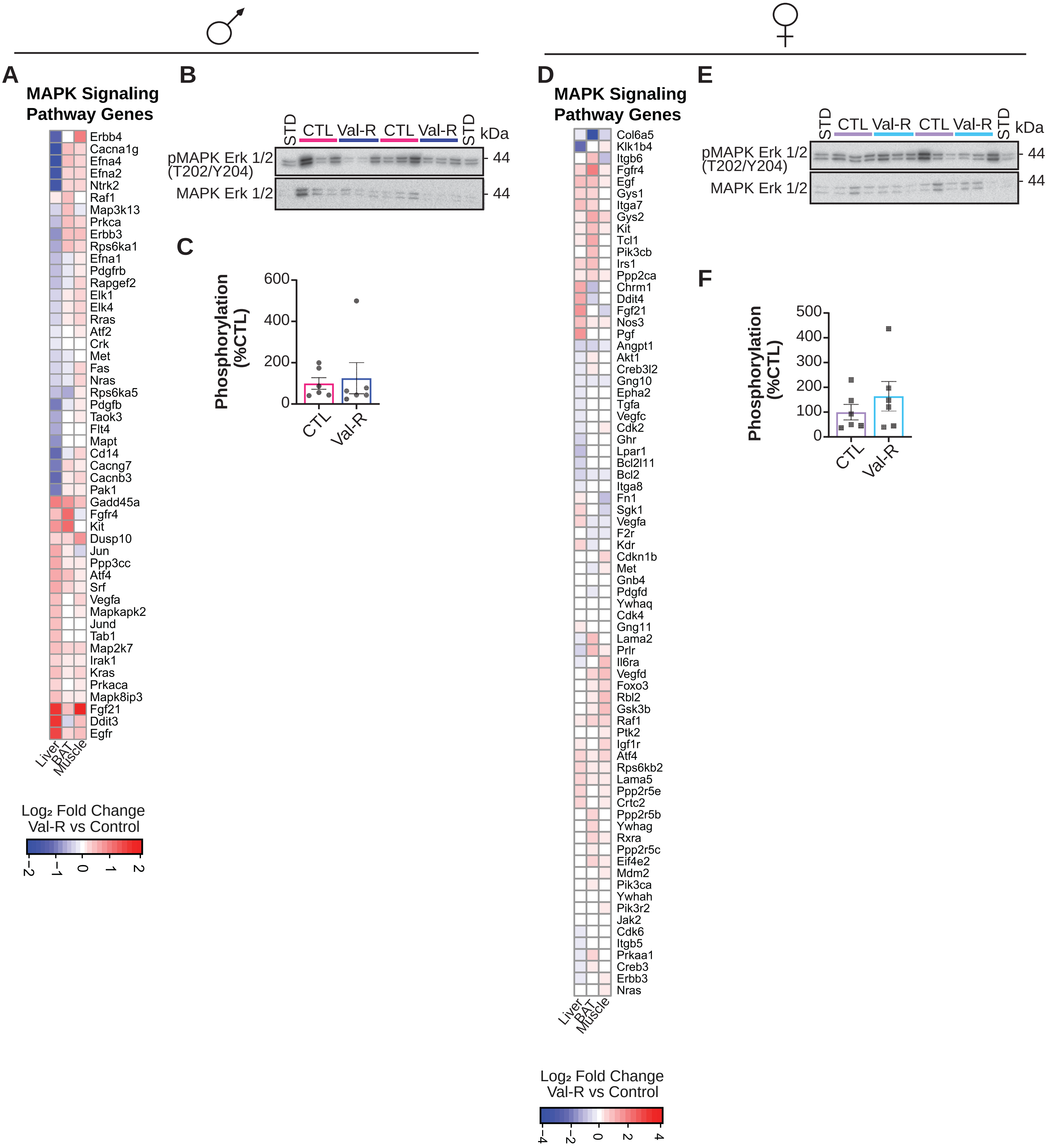

**Supplementary Figure 9: Val-R does not affect MAPK (ERK ½) signaling.**

(A) Altered genes in the MAPK signaling pathway in the liver, muscle and BAT of male mice. Genes shown were significantly altered (Benjamini-Hochberg (BH) adjusted $p<0.05$) by Val-R in at least one tissue in males. (B) Western blots of the analyzed proteins in male livers. (C) Phosphorylation of MAPK ERK 1/2 T202/Y204 normalized to the expression of MAPK ERK 1/2. (D) Altered genes in the MAPK signaling pathway in the liver, muscle and BAT of female mice. Genes shown were significantly altered (Benjamini-Hochberg (BH) adjusted $p<0.05$) by Val-R in at least one tissue in females. (E) Western blots of the analyzed proteins in female livers. (F) Phosphorylation of MAPK ERK 1/2 T202/Y204 normalized to the expression of MAPK ERK 1/2. (C, F) n=6 mice/group; student's t-test. Data represented as mean ± SEM.

### Supplementary Figure 10

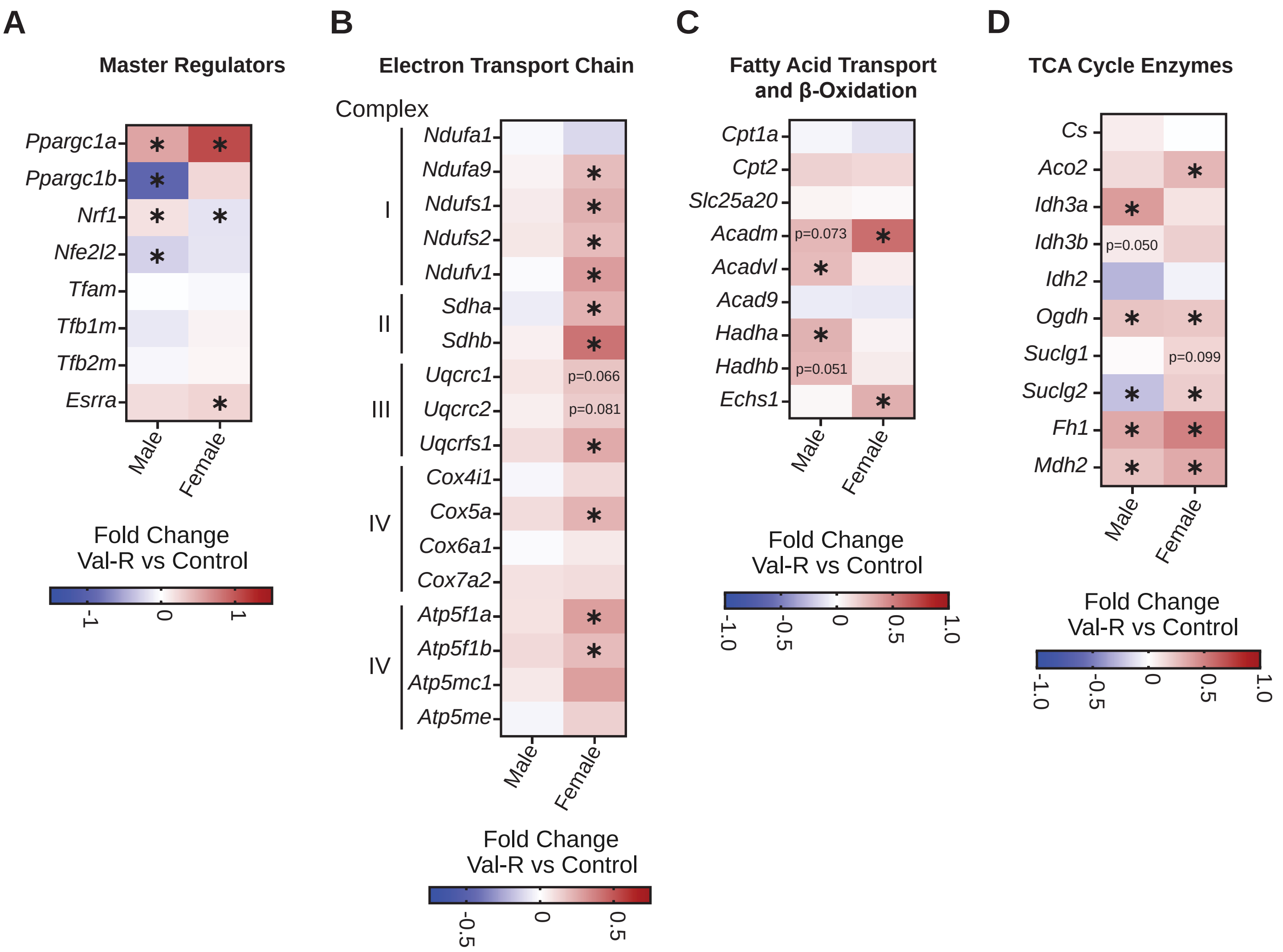

**Supplementary Figure 10: Val-R-fed females display larger mitochondrial metabolism changes in the liver.**

(A-D) $Log_2$ fold-change of gene expression from RNA sequencing analysis in the liver of pre-selected genes related to to mitochondrial master regulation (A), electron transport chain complexes (B), fatty acid metabolism (C) and TCA cycle (D). n=6-10 mice/group. *$p<0.05$, moderated t-test.

#### Supplementary Table Legends

**Supplementary Table 1: Experimental diets.** The composition and calorie content of the experimental diets used in this study.

**Supplementary Table 2: Differentially expressed gene expression.** The statistics from Ebayes method to determine differentially expressed genes.

**Supplementary Table 3: Brown adipose tissue KEGG pathways.** The statistics for the altered KEGG pathways of brown adipose tissue in male and female mice.

**Supplementary Table 4: Liver KEGG pathways.** The statistics for the altered KEGG pathways of the liver in male and female mice.

**Supplementary Table 5: Quadricep muscle KEGG pathways.** The statistics for the altered KEGG pathways of muscle in male and female mice.

**Supplementary Table 6: Mouse lifespan.** Lifespan in days of each mouse in the study.

**Supplementary Table 7: Male mouse frailty index.** Frailty scoring of each male mouse used throughout the study.

**Supplementary Figure 8: Female mouse frailty index.** Frailty scoring of each male mouse used throughout the study.

**Supplementary Figure 9: Full table of correlations and p-values of phenotypes to modules in males.** Pearson correlation coefficient between the gene modules and phenotypic traits, numbers in brackets indicate the corresponding p values in males.

**Supplementary Figure 10: Full table of altered KEGG pathways in the turquoise module.** KEGG pathway enrichment of turquoise module in males.

**Supplementary Figure 11: Full table of correlations and p-values of phenotypes to modules in females.** Pearson correlation coefficient between the gene modules and phenotypic traits, numbers in brackets indicate the corresponding p values in females.

**Supplementary Figure 12: Full table of altered KEGG pathways in the blue module.** KEGG pathway enrichment of blue module in females.

#### Source Data Legends

**Source Data 1**: Raw values for all graphs.

**Source Data 2.** Source Images for liver oil-red-O staining, Senescence-Associated Β-Galactosidase Staining and Western blots.

**Source Data 3.** Source Images for brain Iba1 and GFAP staining.

**Source Data 4**. Source images for microglia and astrocyte skeletal analysis as well as liver and iWAT western blots.
