## Supplementary figures and images for "Lifelong restriction of dietary valine has sex-specific benefits for health and lifespan in mice"

### Source Data 2

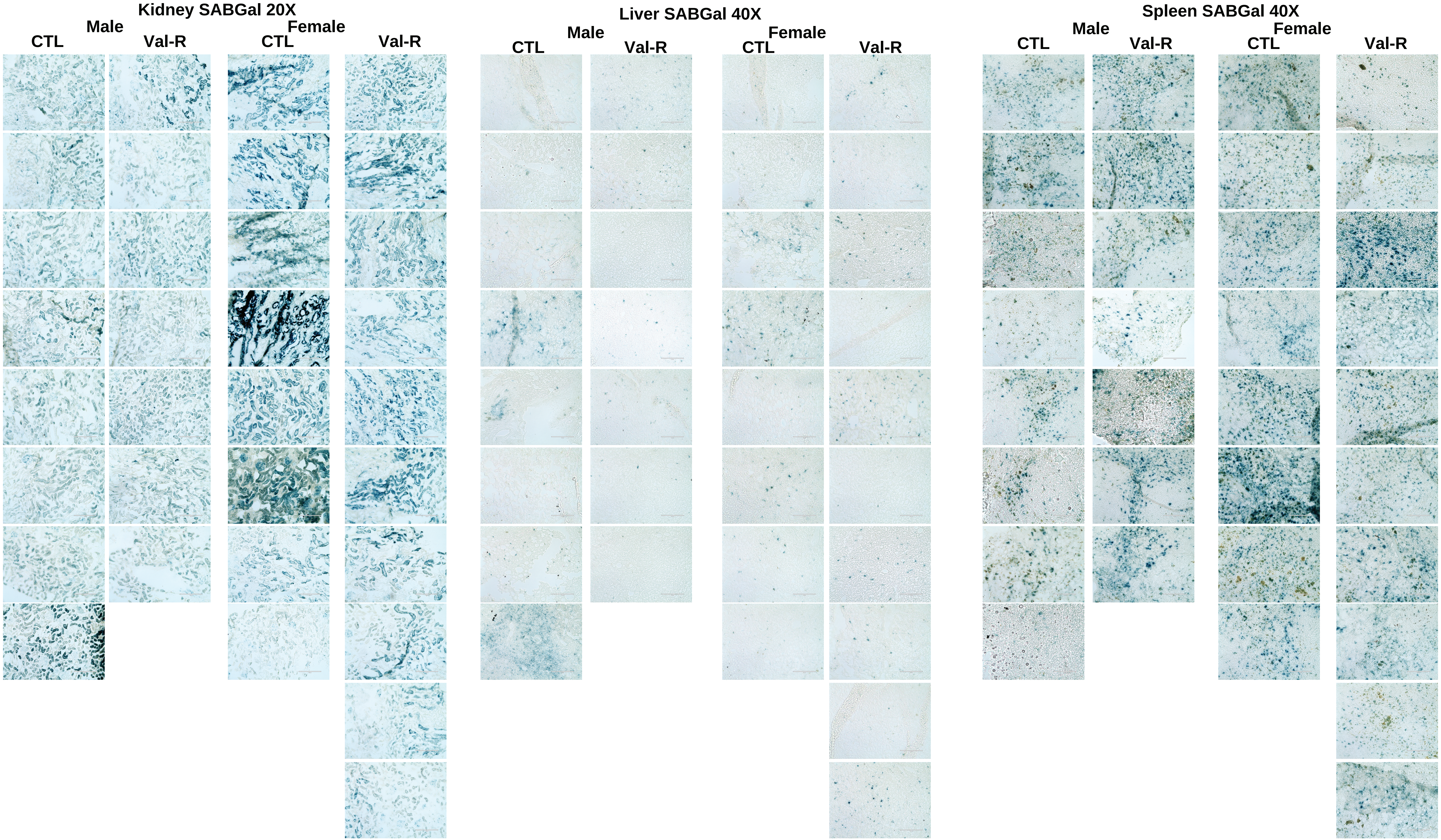

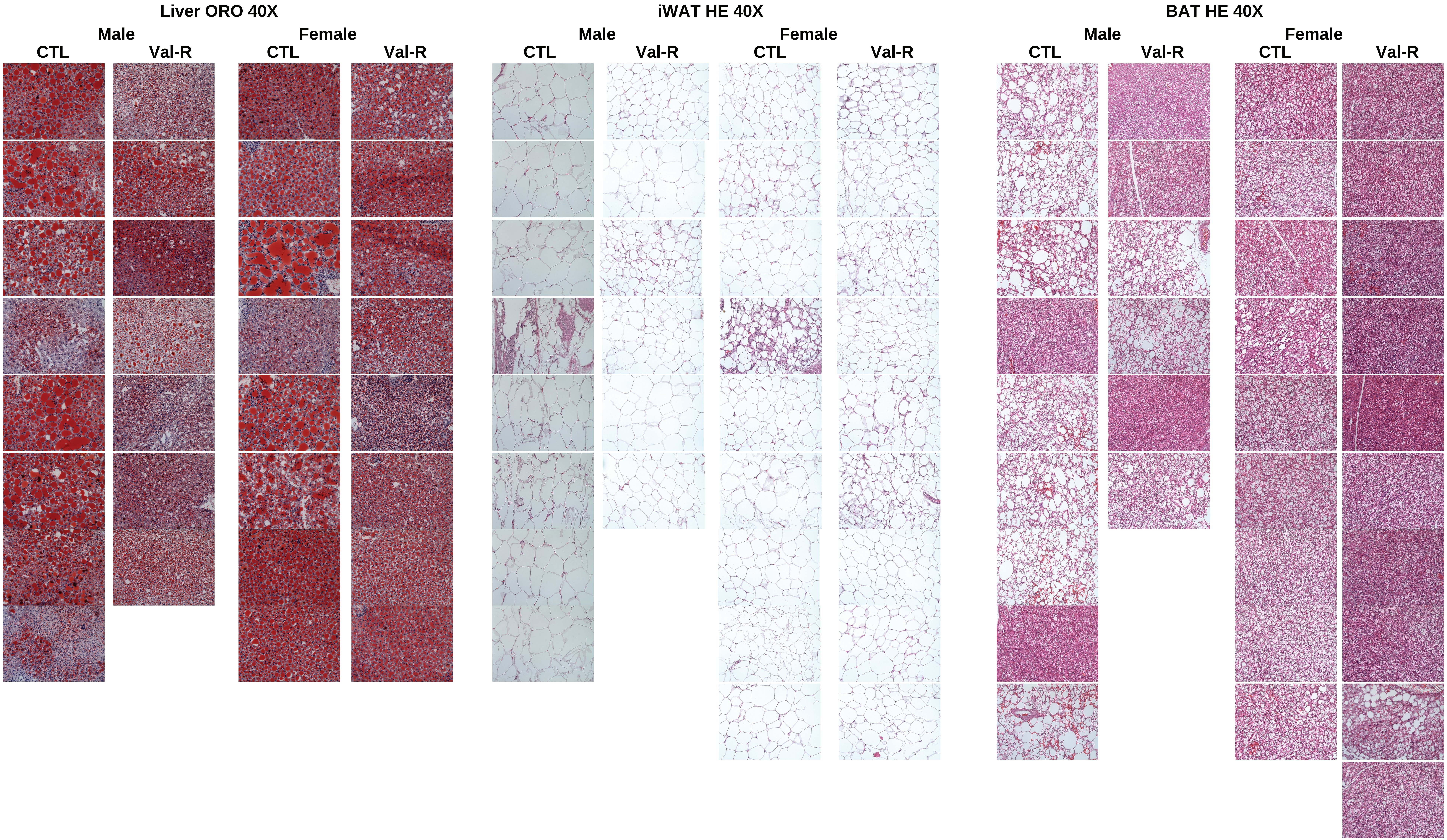

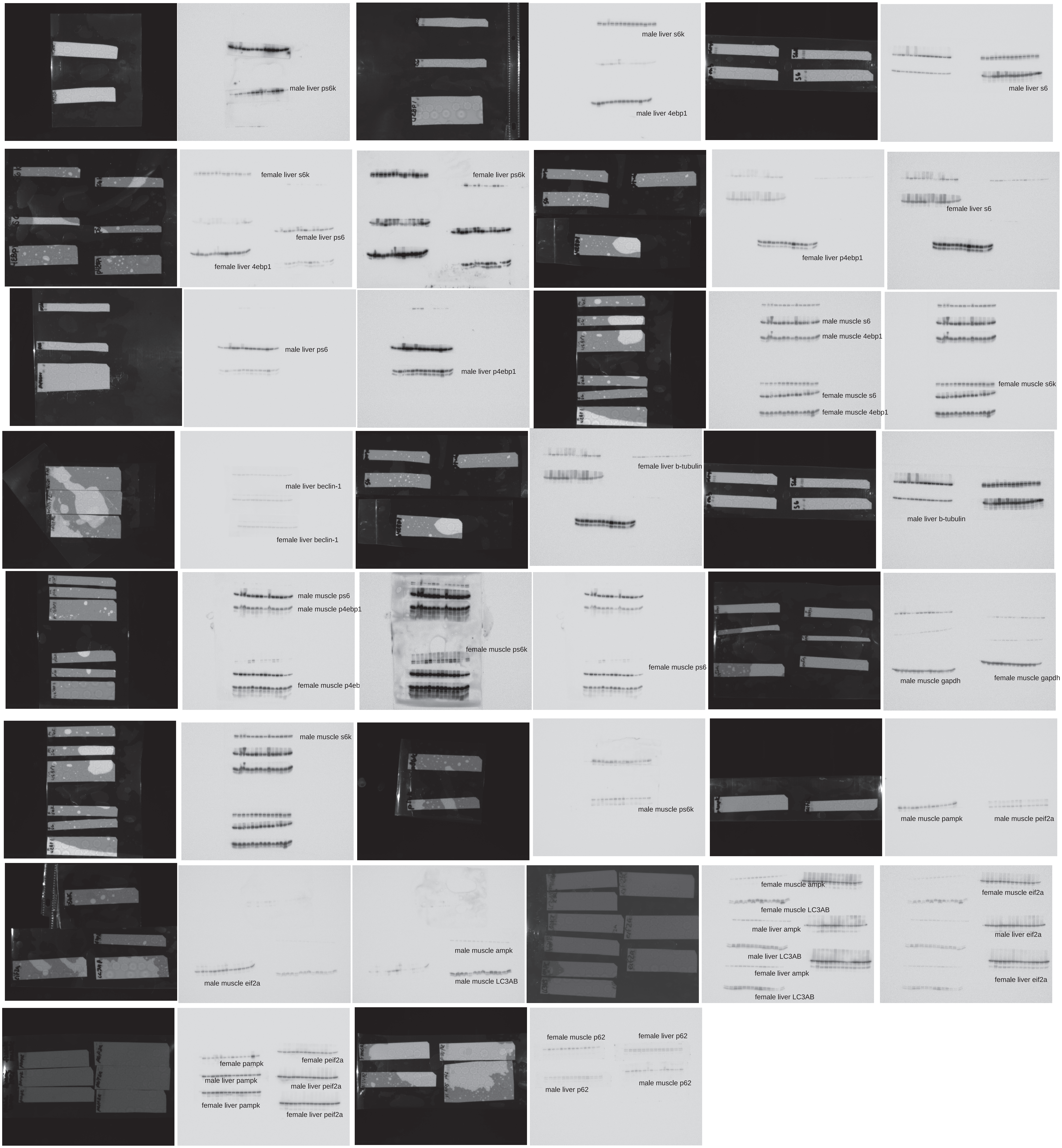

### Source Data 3

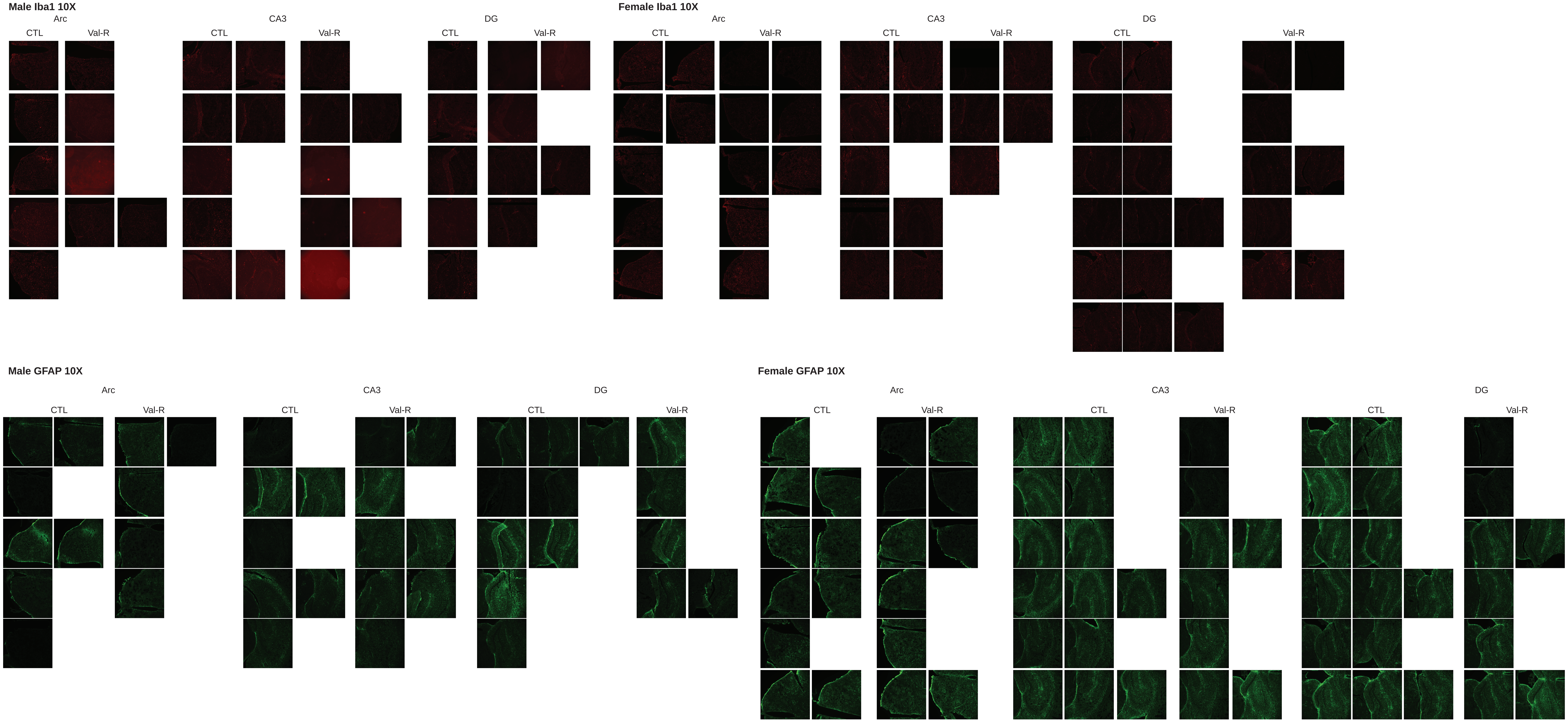

### Source Data 4

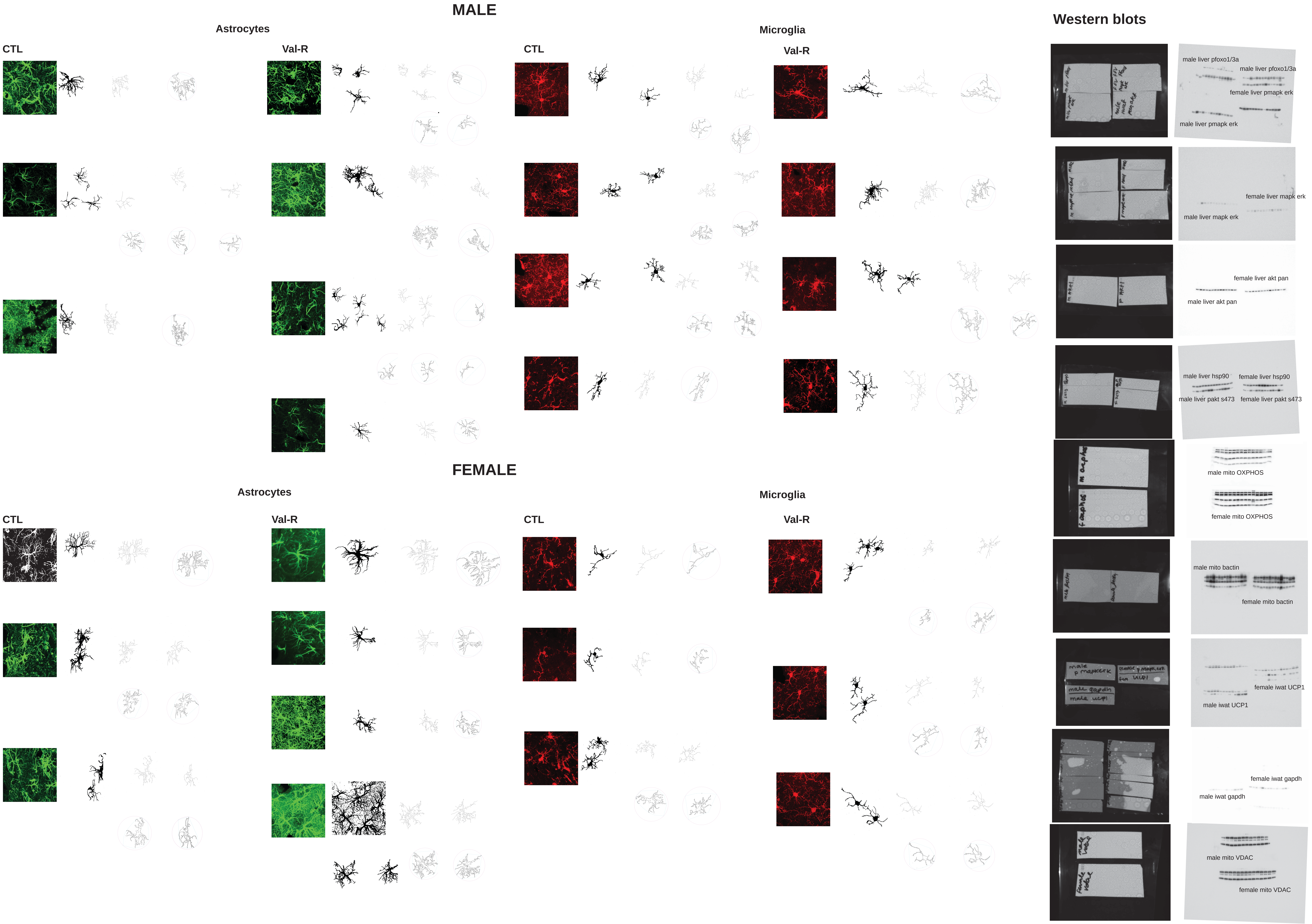
